# RND pumps across the *Acinetobacter* genus; AdeIJK is the ancestral efflux system

**DOI:** 10.1101/2022.10.19.512856

**Authors:** Elizabeth M. Darby, Vassiliy N. Bavro, Steven Dunn, Alan McNally, Jessica M. A. Blair

## Abstract

*Acinetobacter* are generally soil-dwelling organisms that can also cause serious human infections. *A. baumannii* is one of the most common causative agents of *Acinetobacter* infections and is extensively drug resistant. However, an additional 25 species within the genus have also been associated with infection. *A. baumannii* encodes 6 RND efflux pumps, the most clinically relevant class of efflux pumps for antibiotic export, however the distribution and types of RND efflux pumps across the genus is currently unknown. Sixty-three species making up the *Acinetobacter* genus were searched for RND systems within their genomes. We also developed a novel method using conserved RND residues to predict the total number of RND proteins including currently undescribed RND pump proteins. The total number of RND proteins differed both within a species and across the genus. Species associated with infection tended to encode more pumps. AdeIJK/AdeXYZ was found in all searched species of *Acinetobacter*, and through genomic, structural and phenotypic work we show that these genes are actually orthologues of the same system. This interpretation is further supported by structural analysis of the potential drug-binding determinants of the associated RND-transporters, which reveal their close similarity to each other, and distinctiveness from other RND-pumps in *Acinetobacter*, such as AdeB. Therefore, we conclude that AdeIJK is the fundamental RND system for species in the *Acinetobacter* genus. AdeIJK can export a broad range of antibiotics and provides crucial functions within the cell, for example lipid modulation of the cell membrane, therefore it is likely that all *Acinetobacter* require AdeIJK for survival and homeostasis. In contrast, additional RND systems, such as AdeABC and AdeFGH were only found in a subset of *Acinetobacter*, that are associated with infection. By understanding the roles and mechanisms of RND efflux systems in *Acinetobacter*, treatments for infections can avoid efflux-mediated resistance and improve patient outcomes.

**Impact statement:** Efflux pumps extrude antibiotics from within bacterial cells directly conferring antibiotic resistance and underpinning other mechanisms of resistance. By understanding the exact complement of efflux pumps and their roles across infection-causing organisms such as those within the *Acinetobacter* genus, it is possible to understand how cells become resistant to antibiotics and how this might be tackled. Efflux is an attractive target for inhibition to increase susceptibility to existing drugs and therefore, knowing which pumps are present in each species is important. Furthermore, we present a novel method using conserved RND residues to predict the total number of RND proteins including currently novel systems, within bacterial genomes.

**Data Summary:** This study made use of publicly available datasets downloaded from NCBI’s GenBank. A full list of accession numbers can be found in supplementary text 3. Bioinformatics software used in this study was previously published and listed in the methods section. The BLASTp conserved residue files are in S1 text 1 and 2.

The authors confirm all supporting data, code and protocols have been provided within the article or through supplementary data files.

## Introduction

Members of the *Acinetobacter* genus are Gram-negative bacteria commonly isolated from soil and water (1). However, many species are also important human pathogens and *Acinetobacter baumannii* is on the World Health Organisation’s priority pathogens list due to the number of drug resistant infections it causes (2). In addition to *A. baumannii*, a number of other *Acinetobacter* species are known to cause human infections for example *A. lwoffii*, which is the leading cause of *Acinetobacter-derived* bacteraemia in England and *A. nosocomialis* and *A. pittii*, which also cause nosocomial infections (3,4).

*A. baumannii* isolates are commonly multidrug resistant and this is mediated by a combination of molecular mechanisms including acquired resistance genes (e.g. *bla*_oxA-23_) (5), mutations in genes encoding the target of antibiotics, for example *gyrA* (6) and increased expression of multidrug efflux pump systems which actively pump antibiotic compounds out of the cell (7,8). Of particular importance are efflux pumps from the resistance nodulation division (RND) family. RND systems are broadly split into two categories based upon the substrates they export – hydrophobic and amphiphilic efflux pumps (HAE) which contribute to antimicrobial resistance and heavy metal efflux pumps (HME) (9). Typical RND systems are tripartite efflux pumps, which are built around an inner membrane H^+^/drug antiporter (9), the allosteric “pumping” of which allows the drug to be acquired from either the periplasmic space or the outer leaflet of the inner membrane and passed out of the cell *via* a conduit involving the partner outer membrane factor (OMF) channels and periplasmic adaptor proteins (PAPs) (10–13). The RND transporters themselves function as trimers, which contain three functionally interdependent protomers, cycling consecutively through the Loose (L), Tight (T) and Open (O) conformational states during cooperative catalysis (14,15). RND pumps exhibit a broad substrate specificity which is underpinned by the presence of distinct binding pockets within the transporter protomers. The principal binding pockets being known as the ‘Proximal Binding Pocket’ (PBP) and ‘Distal Binding Pocket’ (DBP), which have wide specificities, but are broadly associated with the processing of drugs of different molecular weight and are separated by the so-called gating-, or switch-loop (16–19).

To date, nine RND *genes* have been found in *Acinetobacter: adeJ, adeB, adeE, adeG, adeY, abeD, arpB, acrB* and *czcA* (7,20–28). AdeABC, AdeFGH and AdeIJK have all been characterised in *A. baumannii* and are known to export a broad range of compounds. AdeABC exports aminoglycosides, trimethoprim, chloramphenicol and fluoroquinolones (20,21). AdeFGH also exports trimethoprim, chloramphenicol and fluoroquinolones, but in addition exports tetracycline, tigecycline and clindamycin (7). Lastly, AdeIJK exports chloramphenicol, tetracycline, fluoroquinolones, trimethoprim as well as beta lactams, erythromycin, lincosamides, fusidic acid, novobiocin and rifampicin (23). Expression of AdeABC and AdeFGH can be increased in the presence of an antibiotic challenge, leading to reduced susceptibility to the drug (7,29,30). AdeIJK is constitutively expressed and provides intrinsic levels of resistance to antibiotics. Whilst small increases in expression have been characterised, leading to multi-drug resistance (MDR) phenotypes, increased expression of AdeIJK can be toxic to the cell, therefore increased expression of AdeABC and AdeFGH more commonly mediate MDR (31,32).

The number of RND efflux pumps present varies between bacterial species and also within members of the same species (10,24,33–35). For example, *Neisseria gonorrhoeae* has only one RND system, while *Pseudomonas aeruginosa* can have up to 12, showing that the number of RND genes in a given genome does not necessarily correlate with the ability of bacteria to cause human infections (36). However, it seems plausible that encoding more RND systems may allow a bacterium to adapt to a broader range of environmental stresses.

While a number of efflux pumps have been well-studied in *A. baumannii*, the range of RND systems across the *Acinetobacter* genus is not currently known. In this study we have developed a method to search available genomes for RND efflux systems and have used it to determine their number, type and distribution across the entire *Acinetobacter* genus, and have considered whether these correlate with species that commonly cause human infections. By mapping these data onto the phylogeny of the *Acinetobacter* genus, combined with structural modelling of these systems, we have shown that the pumps currently annotated as AdeIJK and AdeXYZ are actually orthologous RND systems. In addition, we show that AdeIJK is the ancestral pump found across all *Acinetobacter* species and that other RND systems, such as AdeABC have been acquired independently in specific *Acinetobacter* species.

## Methods

### Predicting the total number of RND proteins within a whole genome sequence

Amino acid sequences of characterised HAE proteins from 15 different Gram-negative bacterial species and HME RND proteins from 7 different species were aligned in separate files using MAFFT (v.7) (37). Alignments of the final HAE and HMD proteins used can be found in supplementary S1, text 1 and 2. The consensus sequence in >80% of the aligned sequences was taken from either alignment file to create conserved residue files, supplementary S3, texts 1-2, which can then be searched using BLASTp for other RND proteins within genomes from both *Acinetobacter* and other Gram-negative species (38). An e-value cutoff of 10 was used for the BLASTp command.

For the prediction of RND proteins across the genus, up to 4 reference sequences per *Acinetobacter* species were downloaded from NCBI, totalling 170 genomes. A full list of sequences and accession codes can be found in supplementary S2 table 1. At the time of analysis there were 64 *Acinetobacter* species fully validated by ICNP (https://lpsn.dsmz.de/genus/acinetobacter).

When determining the number of RND proteins in other Gram-negative species the following reference sequences were searched: *A. baumannii* AYE CU459141.1, *C. jejuni* NCTC 11168 GCA_900475265.1, *E. coli* K12 MG1655 NC_000913.3, *H. influenzae* NCTC 8143 GCA_001457655.1, *K. pneumoniae* ATCC 43816 CP064352.1, *N. gonorrhoeae* FA1090 NC_002946.2, *P. aeruginosa* PAO1 GCA_000006765.1, *S. enterica* SL1344 GCA_000210855.2 and *S. flexneri* 5a M90T CP037923.

Furthermore, 100 *A. baumannii* assemblies from NCBI were downloaded to determine if the number of RND proteins differs within a species, supplementary S3, text 3. These assemblies were quality checked using Quast (v.5.0.2) (39), where all assemblies had an N50 of >30,000. Furthermore, their average nucleotide identity across the genome compared to *A. baumannii* AYE (CU459141.1) was confirmed to be > 95% using fastANI (v.1.31) (40). The presence if duplicate *A. baumannii* genomes were detected using MASH (v.2.2.2) to confirm all genomes represented genetically distinct strains (41). The same sequences were used for recombination analysis of AdeABC, AdeFGH and AdeIJK below.

### Finding individual known RND genes across *Acinetobacter*

In addition to searching for the number of RND proteins within the genomes, the presence of known RND genes was also determined across the entire genus. The sequences of RND genes were searched using ABRicate (v.0.8.13) with a custom database comprised of PAP, RND and OMF encoding genes, S3 text 4 (42). Most reference gene sequences were from *A. baumannii* AYE (CU459141.1), apart from *adeDE* from *A. pittii* PHEA-2 (NC_016603.1), *adeXYZ* from *A. baylyi* ADP1 (CR543861.1) and *acrAB* from *A. nosocomialis* NCTC 8102 (CP029351.1). ABRicate cut-off values of >50% identity and >50% coverage were used to highlight orthologs in the different species.

The heatmap displaying the number of RND proteins and presence of RND genes was created using R packages gheatmap in ggtree, ggplot2 and treeio, where the phylogenetic tree and the metadata were visualised (43–46). A literature search was done to determine if a given *Acinetobacter* species had been documented to cause human infection by searching PubMed for the given species and “infection”. The phylogenetic tree of *Acinetobacter* was created using a core gene alignment generated by Panaroo (v.1.2.3) as an input for Fasttree (v.2.1.10) (47). Fasttree was implemented using the generalised time reversible model of evolution (48).

### Genomic context and recombination

To determine if *adeIJK* is found in the same genomic context in three species of *Acinetobacter, A. baumannii* AYE (CU459141.1), *A. lwoffii* 5867 (GCA_900444925.1) and *A. baylyi* ADP1 (CR543861.1), 10 Kb of sequence up and down stream of *adeIJK* was extracted and visualised in Easyfig (v.2.2.5), with tBLASTx homology annotated (38,49). To assess whether *adeABC, adeFGH* or *adeIJK* were found in a recombination hotspots, whole genome alignments of *A. baumannii* (n=100 assemblies, described above, mapped against *A. baumannii* AYE CU459141.1 reference) were created using Snippy (4.6.0) and Gubbins (v.3.1.3), where Gubbins highlighted areas of recombination (50,51). Recombination predictions were visualised in Phandango (52).

### Structural analysis and modelling of *Acinetobacter* RND pump components

Experimental structures of AdeJ from *A. baumannii* in both apo- and eravacycline-bound forms (7M4Q.pdb; and 7M4P.pdb respectively (53)) were used to perform homology modelling of the AdeJ from *A. lwoffii* (76.38% identity) and AdeY from *A. baylyi* (79.25% identity), using I-TASSER (54).

For the analysis of the properties of the drug-binding pockets, the experimental eravacycline-bound structure of *A. baumannii* AdeJ (PDB ID 7M4P, chain B), corresponding to the T-conformer was used, as well as the corresponding T-conformer structures of AcrB occupied by minocycline (PDB ID 4DX5.pdb chain B; (55)) and levofloxacin (PDB ID 7B8T, chain C; (56)). In addition, the L-conformer of the asymmetric AcrB (4DX5.pdb (55)); and the L-conformer of the AdeB in L*OO state (7B8Q.pdb (56)), were used to analyse the drug-binding pockets of AcrB and AdeB respectively, alongside the L-conformer and the ampicillin-bound T-conformer of the MtrD structure (6VKS.pdb; (57)). Sequence alignments have been performed with MAFFT (v.7) (37), and secondary structure visualised with ESpript 3 (58). All visualisations done with the PyMOL Molecular Graphics System, (v.1.8), Schrödinger, LLC.

### Cloning of *Acinetobacter* efflux genes

Efflux pump genes *adeIJK* from *A. baumannii* AYE and *adeXYZ* from *A. baylyi* ADP1 (supplementary S2, table 2) were cloned using NEB HiFi cloning into the expression vector pVRL2 (59). Briefly, the efflux genes were amplified using PCR and primers in supplementary S2, table 3. The vectors were digested using NotI-HF and XmaI restriction endonucleases (New England Biolabs) that left complementary overhangs to the PCR products. The PCR products and digested vectors were then ligated using HiFi assembly mix. Cloned vectors were transformed into *A. baumannii* ATCC 17978 *ΔadeAB ΔadeFGH ΔadeIJK* to determine function. The complete sequence of cloned vectors was determined by Plasmidsaurus, sequencing files in supplementary files S5 and S6 (60).

### Antimicrobial Susceptibility

The minimum inhibitory concentration was measured using broth micro-dilution method according to CLSI guidance with 1% arabinose (Acros Organics) (61,62). Compounds were chosen because they are exported by different Ade systems: Ampicillin (Sigma), Chloramphenicol (Sigma), Ciprofloxacin (Acros Organics), Clindamycin hydrochloride (TCI Chemicals), Ethidium bromide (Acros Organics), Rifampicin (Fisher) and Tetracycline (Sigma).

## Results

### Development of BLASTp database to detect RND proteins

The number of RND proteins is known to differ between different bacterial genera. Here, we developed a method to quantify the number of HAE and HME RND proteins within a genome sequence based upon conserved residues in characterised RND proteins. To do this, the sequences of 24 known RND genes from 16 species were aligned to determine the conserved residues which could be used to search for known and unknown RND genes in genome sequences of Gram-negative bacteria. Due to the degree of difference in sequences between HAE and HME pumps, two separate alignments were created. From each alignment residues that were the same in 80% of sequences or more were used to create a conserved residue file, with which BLASTp could search genomes to determine the number of each type of RND pump (supplementary S3, texts 1-2). To validate this method, the number of RND pumps in well characterised type strains of Gram-negative bacteria were determined, shown in table 1.

**Table 1:** Number of RND proteins within the genomes of Gram-negative bacteria, as determined by BLASTp of the conserved RND residues.

| Species | Accession | Number of predicted HAE proteins | Number of predicted HME proteins | Total number of proposed RND proteins in the literature |
| --- | --- | --- | --- | --- |
| *Acinetobacter baumannii* ACICU | GCA_000018445.1 | 5 | 1 | 6 (63) |
| *Acinetobacter baumannii* AYE | CU459141.1 | 4* | 1 | 6 (63) |
| *Campylobacter jejuni*<br>NCTC 11168 | GCA_900475265.1 | 2 | 0 | 2 (10) |
| *Escherichia coli* K12 | NC_000913.3 | 6 | 1 | 7 (64) |
| *Haemophilus influenzae* NCTC 8143 | GCA_001457655.1 | 1 | 0 | 1 (65) |
| *Klebsiella pneumoniae* ATCC 43816 | CP064352.1 | 8 | 1 | 9 (10,66,67) |
| *Neisseria gonorrhoeae* FA1090 | NC_002946.2 | 1 | 0 | 1 (68) |
| *Pseudomonas aeruginosa* PAO1 | GCA_000006765.1 | 10 | 1 | 11 (10) |
| *Shigella flexneri* 5a M90T | CP037923 | 5 | 1 | 6 (69) |
| *Salmonella enterica* serovar Typhimurium SL1344 | FQ312003.1 | 6 | 0 | 5 (10) |

\* AB AYE results were missing ArpB due to sequence dissimilarity with the RND proteins used in our alignments. This is discussed further below.
*S. enterica* Typhimurium is shown separately at the bottom because an additional RND efflux protein, to those described in the literature, was found.

For *A. baumannii, C. jejuni, E. coli, H. influenzae, N. gonorrhoeae, P. aeruginosa, K. pneumoniae* and *S. flexneri* the number of RND efflux pumps detected using our method matched that in the literature for the species tested, validating this approach. For example, *N. gonorrhoeae* is well known for encoding only one RND protein (68) and this was also true when FA1090 was tested using our method. For *E. coli* K12 all 7 known RND proteins were detected, including the HME protein CusA. CusA was identified in both the HAE and HMD RND protein searches, but only included once in table 1. Interestingly, we were also able to find an additional RND protein in *S. enterica* serovar Typhimurium, with homology to OqxB.

In *A. baumannii* AYE, however, the BLASTp searches only identified 5 out of 6 known RND proteins. Both HAE and HMD searches failed to highlight ArpB, an RND-like protein that is involved in opaque to translucent colony formation switching (27). ArpB was added to both the HAE and HME alignments, but due to differences in the sequences of the other RND proteins compared to it, the number of conserved residues reduced dramatically across the aligned proteins and rendered the method unable to then detect any RND proteins successfully via BLASTp. Phylogenetic trees based upon the alignments with ArpB are shown in supplementary S1, figures 3 and 4 and show that ArpB clusters separately from the other proteins. Interestingly, when using BLASTp to estimate the number of HAE RND proteins across 100 sequences of *A. baumannii (*including *Acinetobacter baumannii* ACICU), in 95% of the sequences all 5 proteins were detected, supplementary S2, table 4. This suggests that ArpB can be identified by the search and that the sequence of ArpB in *A. baumannii* AYE differs from that of other *A. baumannii* sequences. When directly comparing the amino acid sequence of ArpB from AYE and ArpB from ACICU, they are only 24.5% identical with a coverage of 94%.

Of the 100 *A. baumannii* sequences analysed 95% had 5 HAE RND proteins, 3% had 4, and 2% had 6/7. In the three sequences that had only 4 RND proteins, they were missing AdeB and a subsequent targeted BLASTn search for *adeB* provided no results. The additional RND protein in the other two sequences (GCA_000302135.1 and GCA_000301875.1) is currently not characterised and when looking more closely at the 7^th^ protein identified in GCA_000301875.1, it seems to be truncated RND protein with sequence similarity to AdeB and found next to ArpB.

In the 100 *A. baumannii* sequences the number of HME pumps also differed within the species. 67% of sequences had only 1 heavy metal pump protein – CzcA, supplementary S2, table 5. Although another inner membrane protein (CzcD) exists in the Czc system, its sequence is much shorter than the HME proteins in the alignment, so it isn’t found using the BLASTp search. The remaining sequences had an additional 1-3 proteins in their genomes. Where, 27% had a total of two HME proteins, CzcA and a protein annotated as CusA. Furthermore, 5% had 3 HME proteins where the third protein was also labelled as CzcA, but wasn’t found with CzcD, so it is likely to be annotated incorrectly and could represent a third distinct heavy metal efflux system in *A. baumannii*. Finally, one sequence had 4 HME proteins and three of them were annotated as CzcA and one as CusA. Of the three CzcA, one was found in the expected operon with other Czc proteins and the other two are presumably proteins with sequence similarity to CzcA.

Table 1 shows that not only does the method to identify RND proteins work across a broad range of Gram-negative species, it has also highlighted uncharacterised RND proteins within genomes. The method was further applied to determine the number of RND proteins in every species of the *Acinetobacter* genus, figure 1, column 1.

**Figure 1:**
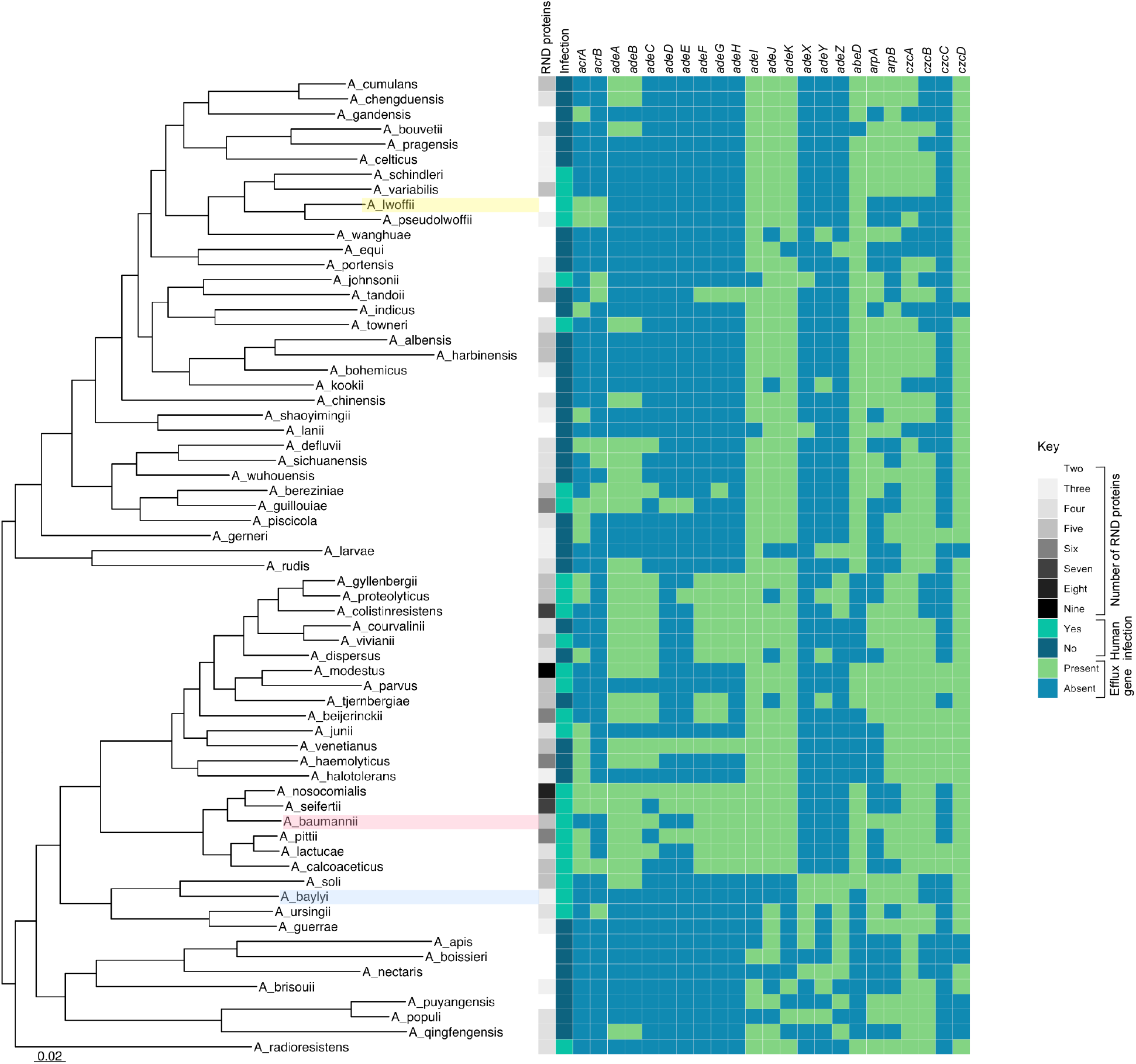
The presence of RND pumps across the Acinetobacter genus. Heatmap of *Acinetobacter* genus with characterised RND genes presence/absence and the number of RND proteins. For column 1, number of RNDs, the mean average of the total number of (both HME and HAE) RND efflux proteins for each species was determined, to 1 significant fgure place, where the greater the number of proteins, the darker grey the colour. For column 2, if a species has been shown in the literature to cause infection it is turquoise. Subsequent columns 3-25 are highlighted green if the efflux gene was found in the reference sequences using ABRicate. *A. Iwoffii* is highlighted yellow, *A. baumannii* is pink and *A. baylyi* is blue.

### The number of RND proteins differs across the *Acinetobacter* genus

The number of RND proteins in species of *Acinetobacter* ranged from 2 to 9, figure 1 column 1, and this correlated with whether that species is known to cause human infection. When doing a Pearson’s correlation between number of RND proteins and ability to cause infection the r^2^ value was 0.1411, and therefore 14% of the variance in infectivity can be explained by number of RNDs, which was statistically significant (p=0.002). Therefore, infection causing species generally encoded more RND genes than those that have not been reported to cause infection. Whilst there are likely to be more *Acinetobacter* species that have the capacity to cause human infection, the heatmap documents those published to date.

Previous work has shown that not all species have the same number of total RND components, for example in *A. baumannii* around 20% of all isolates are missing the OMF AdeC and up to 25-30% have been shown to be missing the RND protein AdeB (24,33,70). This is also evident in our data; whilst the number of RND proteins differs across the *Acinetobacter* genus, the number of RND proteins also differs between members of the same species. For example, in *A. colistiniresistens* sequences there were between 3 and 5 HAE RND proteins and between 1 and 4 HME RND proteins. The mean average of the total RND proteins, 7, highlighted in the four *A. colistiniresistens* sequences tested is shown in figure 1. Furthermore, this variation is seen in other species for example *A. bereziniae*, where 3-4 HAE and 1-3 HME proteins were highlighted as well as in *A. haemolyticus, A. pittii, A. proteolyticus, A. tandoii* and above in the 100 *A. baumannii* sequences.

### Presence of RND efflux pumps across the *Acinetobacter* genus

In order to get a broader picture of what RND genes, including periplasmic adaptor protein (PAP) genes and outer membrane factor (OMF) genes, are present in the *Acinetobacter* genus, genomes from all validated *Acinetobacter* species were searched for the presence of these genes using a custom database of *Acinetobacter* RND genes in ABRicate, supplementary S3, text 4. Figure 1 shows a heatmap, where the presence (green) and absence (blue) of RND genes is mapped onto a phylogenetic tree of the genus. The average number of RND proteins, as determined by the novel RND residue BLASTp search, is also plotted in greyscale, column 1. The average number of RND proteins sometimes over or underestimates the number compared to the RND genes found by ABRicate, this is because ABRicate searched only one reference sequence and the number of RND proteins is the average of up to four sequences searched by BLASTp. When directly comparing the same sequence using BLASTp to search for proteins with conserved RND residues and ABRicate to highlight all characterised RND genes, BLASTp finds the same or more efflux pump proteins compared to ABRicate in 38 species. In the remaining 27 species, BLASTp found 75% of the RND proteins highlighted by ABRicate. Therefore, a combination approach of both methods provides the best resolution when looking for RND genes and proteins. In total BLASTp found 274 proteins with conserved RND residues and ABRicate found 272 characterised RND genes.

Parts of the metal ion efflux system, Czc, are also common across the genus where *czcD* and *czcA*, coding for the inner membrane proteins, are found in almost all species. In contrast, other RND systems, such as *adeABC* and *adeFGH*, are commonly found in a clade comprised of *A. baumannii* and closely related species, which have a higher propensity to cause human infection.

Notably, almost all *Acinetobacter* species encode *adeIJK* and it is striking, that those that do not, encode the *adeXYZ* operon instead. This is true in all but three species, *A. colistiniresistens, A. gyllenbergii* and *A. proteolyticus*, which encode genes that are similar to *both adeK* and *adeZ* according to the ABRicate search. Indeed, when looking more closely at these, it seems that they have a full *adeIJK* operon, but also an additional RND operon with an OMF that is 69-71% identical to *adeK*, found with a PAP and RND protein.

### *A. baumanii adeIJK* and *A. baylyi adeXYZ* are orthologous efflux systems

Originally, *adeXYZ* was described in *A. baylyi* and stated to have high sequence similarity to *adeIJK* from *A. baumannii* (24,71). The fact that the absence of *adeIJK* in figure 1 seems to match almost perfectly to the presence of *adeXYZ* suggested that these pumps may be divergent examples of the same pump, rather than distinct systems. Therefore, the sequence, genomic location, function and structure of *adeIJK* from *A. baumannii* AYE, and *adeXYZ* from *A. baylyi* ADP1 were compared. In addition, another *adeIJK* from a more phylogenetically distant *Acinetobacter* (*A. lwoffii* 5867) was included for context.

In *A. baumannii* AYE, *A. lwoffii* 5867 and *A. baylyi* ADP1, the *adeIJK* and *adeXYZ* operons are found in the same genomic location, with conserved regions up and immediately downstream of the operon, figure 2. The genes flanking the *adeIJK/XYZ* operons encode PAP2 phosphatase family proteins and YbjQ family proteins. Despite the high level of conservation around the operons, downstream from the OMF the sequences differ dramatically. Neighbouring genes that could be annotated by Prokka (72) are also included in figure 2, however there are discrepancies where orthologous genes are annotated differently in different species.

**Figure 2:**
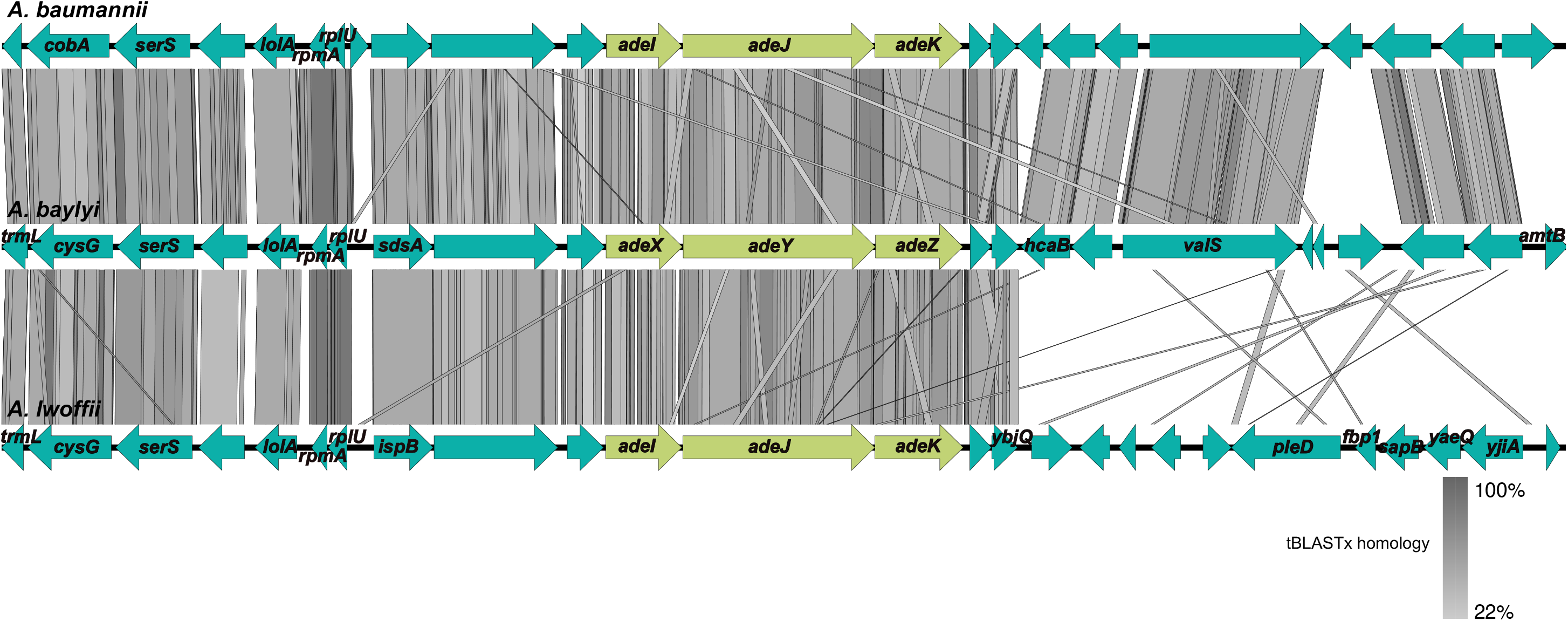
The genomic context of *adeIJK* and *adeXYZ* is identical. Figure was created in Easyfig (v.2.2.5) using *A. baumannii* AYE, *A. baylyi* ADP1 and *A. lwoffii* 5867 sequences plus and minus 10 Kb from the *ade* operons, annotated using Prokka (72). The grey scale shows tBLASTx homology and arrows refer to coding regions within the genome. Immediately around *adeIJK/XYZ* is conserved but differs further downstream of the OMF. *A. baumannii* and *A. baylyi* are more similar downstream of the OMF, compared to *A. lwoffii*. The two genes immediately after each OMF are genes which encode YbjQ family proteins and this is conserved in all three species (*Abau:* HKO16_14475,HKO16_14480, *Abay:* KJPEBFEI_02742, KJPEBFEI_02743, *Alwo:* NCTC5867_02643*, ybjQ*. The gene immediately upstream of AdeI/X is a gene encoding a PAP2 phosphatase family protein (*Abau:*HKO16_14455, *Abay:*KJPEBFEI_02738, *Alwo:*NCTC5867_02647).

The periplasmic adaptor proteins, *adeI* and *adeX*, in *A. baumannii, A. lwoffii and A. baylyi* are between 62-70% similar. The sequence of *adeI* from *A. baumannii* is more similar to *adeX* from *A. baylyi* than *adeI* from *A. lwoffii*. The same pattern can be seen for the RND (*adeJ/Y*) and OMF (*adeK/Z*) genes, where the nucleotide sequences from *A. baumannii* for each component of the tripartite pump are more like the sequences from *A. baylyi* than *A. lwoffii*, despite them sharing the nomenclature with *A. lwoffii*, figure 3.

**Figure 3:**
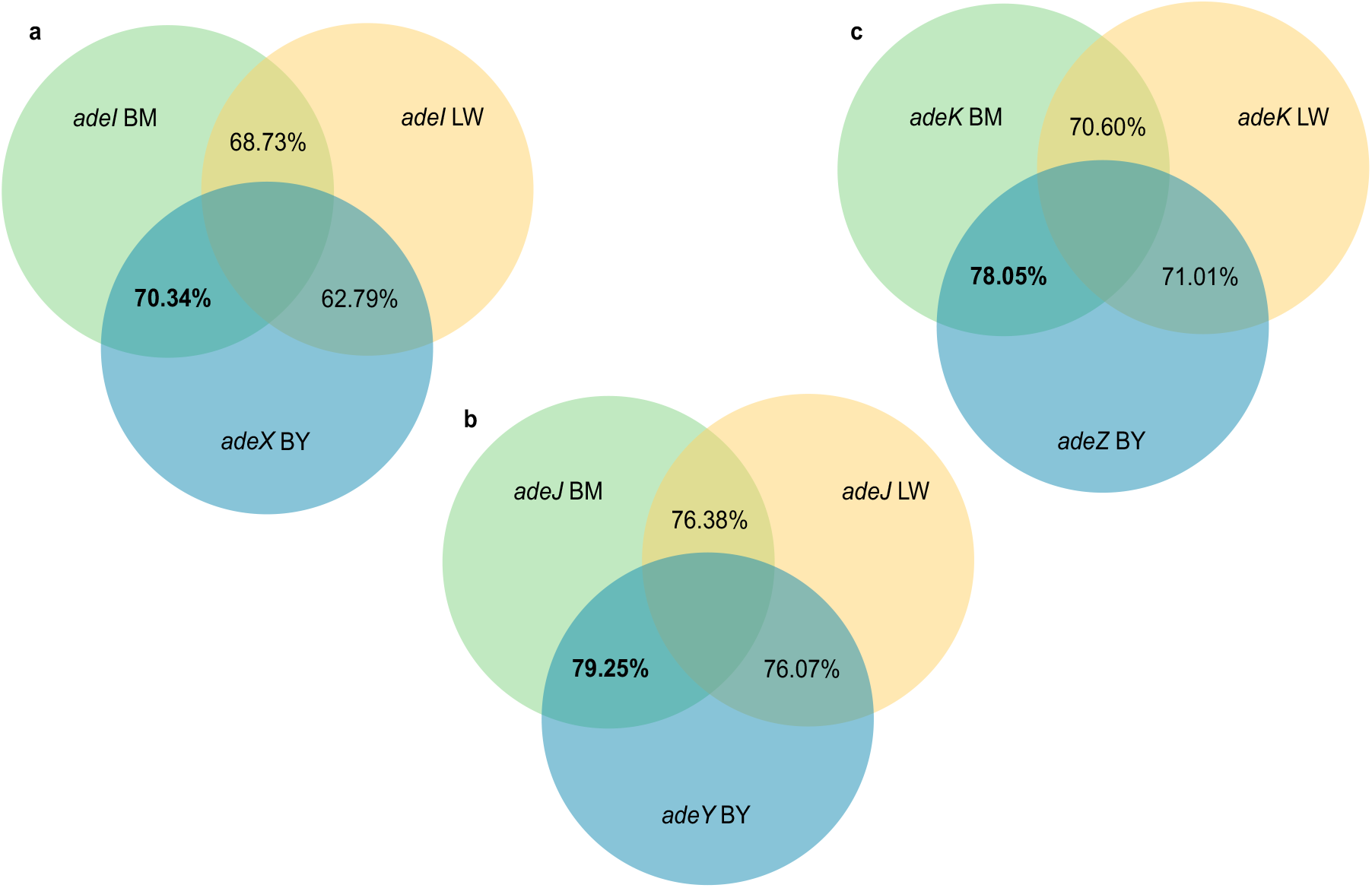
Venn diagrams of percentage identity between *Acinetobacter* RND genes. Percentage identity was determined by a MAFFT (v.7) alignment of the PAP, RND and OMF genes, which was analysed by Sequence Manipulation Suite (73). BM- *A. baumannii* (green) LW-*A. lwoffii* (yellow) BY*-A. baylyi* (blue) a – PAP b – RND c- OMF. In bold are most similar pair for PAP, RND and OMF based upon nucleotide % identity.

Given that *adeIJK/XYZ* is found in all *Acinetobacter*, but *adeABC* and *adeFGH* are found only in a subset of species, the recombination levels and polymorphisms in and around all three *ade* operons were analysed. To determine if there were any recombination hotspots and polymorphisms around *adeIJK*, Gubbins was used to infer recombination levels across the whole genomes of 100 *A. baumannii* sequences. Figure 4 shows the recombination predictions for *adeABC* (e and f)*, adeFGH* (c and d) and *adeIJK* (a and b) across the sequences. Of the three systems, there is no signature of recombination seen around the *adeIJK* genes across these sequences indicating this is an ancestral operon common across all genomes studied here. High levels of recombination are seen around *adeABC* and since it is found in *A. baumannii* and other infection-causing species (figure 1) but not all *Acinetobacter*, it is likely that this operon is in a recombination hotspot where genes are acquired by horizontal gene transfer. Polymorphisms are also shown in figure 4 (red and blue blocks), where more are seen around AdeABC. Further to this, the level of recombination and polymorphisms is also low around *adeFGH*.

**Figure 4:**
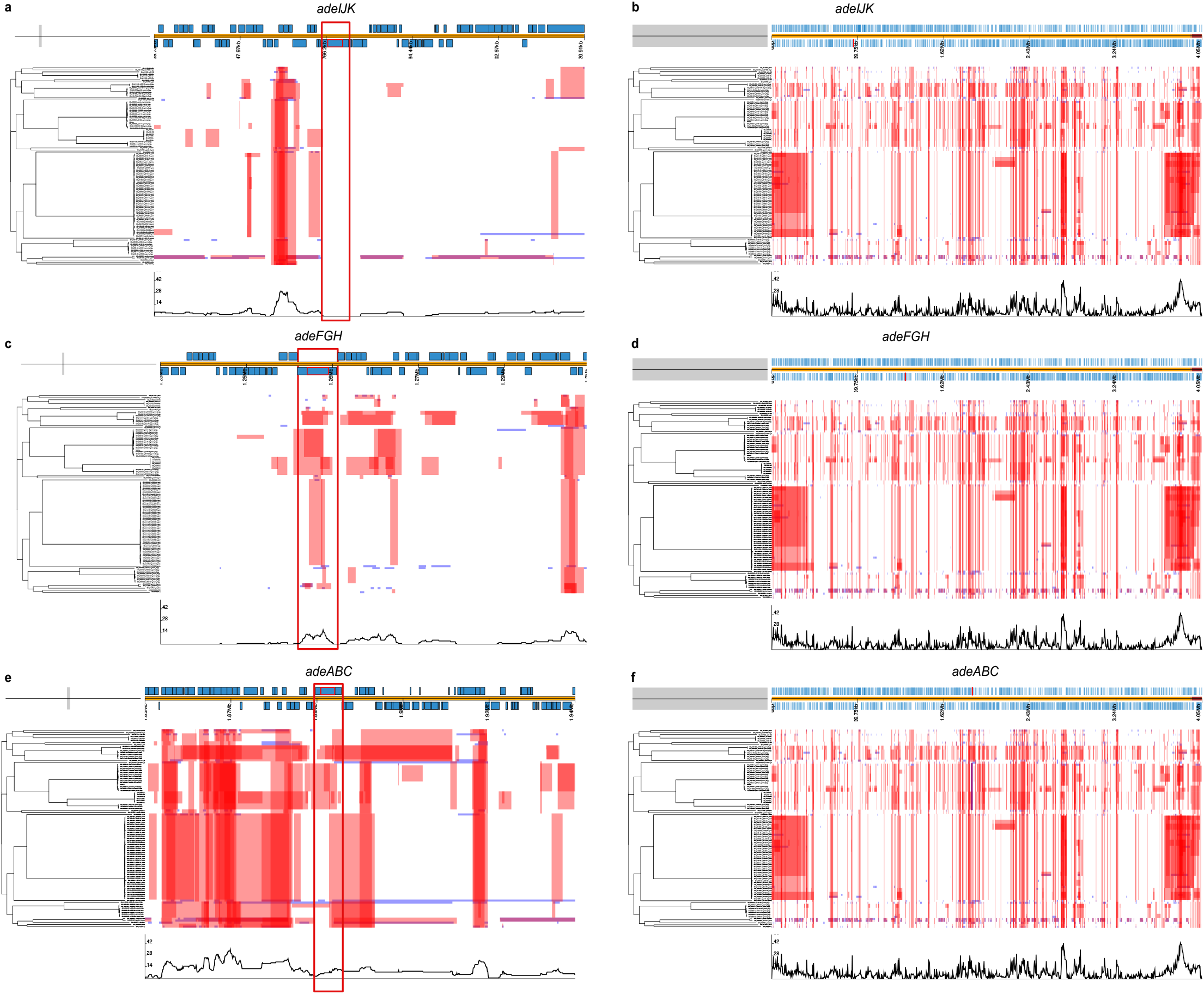
AdeIJK has fewer SNPs within it and lower levels of recombination surrounding it than AdeABC or AdeFGH. 100 *A. baumannii* genomes were aligned against reference *A. baumannii* AYE and the presence of polymorphisms and recombination was determined using Gubbins. Parts a, c and e are zoomed in parts of the genome at each ade operon, showing the levels of SNPs (red and blue squares, red are ancestral SNPs) and recombination levels (the black line on the bottom, the higher the peak the more recombination). The right hand side, b, d and f, show the entire genome and the position of each different *ade* operon, which is highlighted red underneath the label. All figures have an associated phylogenetic tree created by Snippy to show the relatedness of the *A. baumannii* sequences. AdeIJK (a) has fewer SNPs and recombination than AdeABC (e) and AdeFGH (c) indicating it is highly conserved.

### The phenotypic impact of *adeIJK* and *adeXYZ* expression is the same

To determine whether the substrate profile of AdeIJK and AdeXYZ are the same, *adeIJK* from *A. baumannii* AYE and *adeXYZ* from *A. baylyi* ADP1 were cloned and expressed in *A. baumannii* ATCC 17978 *ΔadeAB ΔadeFGH ΔadeIJK* and susceptibility to known substrates of different Ade systems was measured. The effect of expression of AdeIJK and AdeXYZ in ATCC 17978 *ΔadeAB ΔadeFGH ΔadeIJK* was identical with both conferring decreased susceptibility to chloramphenicol, ciprofloxacin, clindamycin, tetracycline and rifampicin.

Initially, these plasmids were expressed in ATCC 17978 lacking only *adeIJK (ΔadeIJK*), which increased the ethidium bromide (EtBr) MIC from 1 to 64 μg/mL (data not shown). However, in the triple pump deletion background *ΔadeAB ΔadeFGH ΔadeIJK*, expression of neither AdeIJK nor AdeXYZ altered susceptibility to EtBr (Table 2) suggesting AdeIJK does not export EtBr as shown previously (23). The basis for this difference in relation to strain background is not fully understood but has been reported previously (23).

**Table 2:**
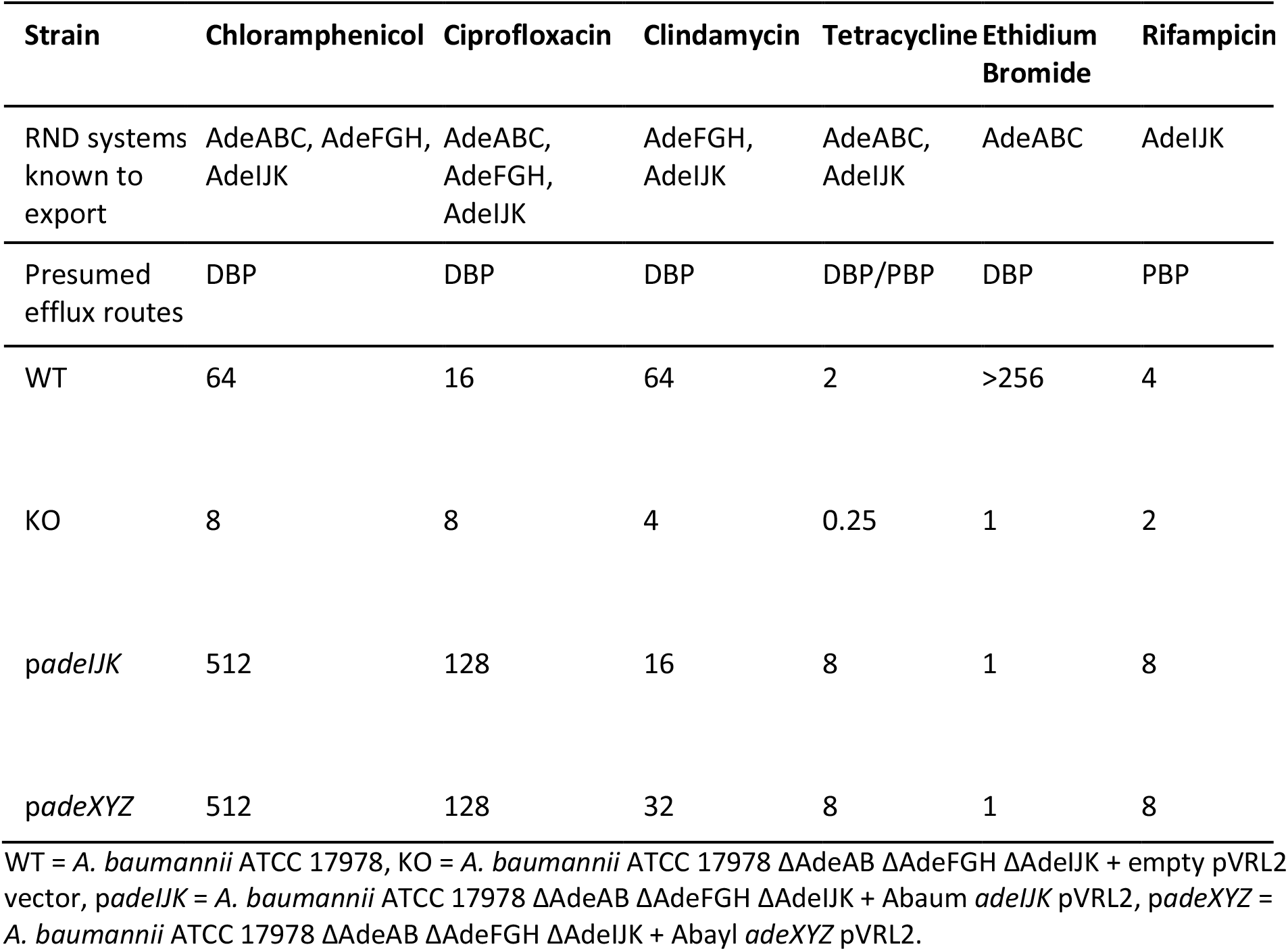
Broth microdilution MIC results (μg/mL) for AdeIJK and AdeXYZ when expressed in *A. baumannii* ATCC 17978 *ΔadeAB ΔadeFGH ΔadeIJK*.

| Strain | Chloramphenicol | Ciprofloxacin | Clindamycin | Tetracycline | Ethidium Bromide | Rifampicin |
| --- | --- | --- | --- | --- | --- | --- |
| RND systems known to export | AdeABC, AdeFGH, AdeIJK | AdeABC, AdeFGH, AdeIJK | AdeFGH, AdeIJK | AdeABC, AdeIJK | AdeABC | AdeIJK |
| Presumed efflux routes | DBP | DBP | DBP | DBP/PBP | DBP | PBP |
| WT | 64 | 16 | 64 | 2 | >256 | 4 |
| KO | 8 | 8 | 4 | 0.25 | 1 | 2 |
| *padeIJK* | 512 | 128 | 16 | 8 | 1 | 8 |
| *padeXYZ* | 512 | 128 | 32 | 8 | 1 | 8 |

WT = *A. baumannii* ATCC 17978, KO = *A. baumannii* ATCC 17978 $\Delta AdeAB \Delta AdeFGH \Delta AdeIJK$ + empty pVRL2 vector, *padeIJK* = *A. baumannii* ATCC 17978 $\Delta AdeAB \Delta AdeFGH \Delta AdeIJK$ + *Abaum adeIJK* pVRL2, *padeXYZ* = *A. baumannii* ATCC 17978 $\Delta AdeAB \Delta AdeFGH \Delta AdeIJK$ + *Abayl adeXYZ* pVRL2.

### AdeJ from *A. baumannii* and AdeY from *A. baylyi* are structurally similar

Since the AdeIJK from *A. baumanii* and AdeXYZ from *A. baylyi* are genetically and functionally similar, we next examined whether their respective RND-transporters also shared similar structural features using comparative analysis of the experimentally available structures and making homology models of the missing ones. Again, *A. lwoffii* AdeJ was included for context.

The high-fidelity homology modelling of the AdeJ from *A. lwoffii* (76.38% identity) and AdeY from *A. baylyi* (79.25% identity) was enabled by recent determination of the experimental structures of AdeJ from *A. baumannii* (*Ab*AdeJ) in both apo- and eravacycline-bound forms (7M4Q.pdb; and 7M4P.pdb respectively (53)). The amino acid sequence of AdeJ from *A. baumannii* (*Ab*AdeJ), *A. lwoffii* (*Al*AdeJ) and *A. baylyi* AdeY align without any gaps, allowing for one-to-one positional correspondence between them (figure 5), with the sole exception of a single residue insertion after position 602 (*Ab*AdeJ numbering) in both *A. lwoffii* AdeJ and AdeY, which maps to the protein surface (PC1 sub-domain), and thus should not directly affect drug binding. There is also a high level of conservation with other members of the RND-transporter family, including AdeB (56,74), AcrB (55,75) and MtrD (57). Indeed, reflecting it is the very close geometry of these transporters, with the superposition of the AdeJ structure and the ligand-occupied T-conformers AcrB (7M4P:B and 4DX5:B respectively) yielding a strikingly low RMSD of 1.17Å, allowing for direct comparison and interpretation of their binding pockets (supplementary S4, figure 5). As our current knowledge of substrate recognition within RND pumps derives primarily from AcrB, this close relation allows for an unambiguous assignment of the binding determinants between the pumps. Accordingly, the analysis of the residues lining the proximal and the distal binding pockets, which are implicated in the processing of the high-molecular weight (including macrolides and rifampins) and lower-molecular weight/planar compounds (e.g. tetracyclines, fluoroquinolones and beta-lactams) respectively (55,56,76,77), identified high levels of positional conservation of the residues previously described as forming the recognition determinants of these binding pockets (19,55,76) (annotated in figure 5) between the AdeJ and AdeY, fitting with the antimicrobial susceptibility data in table 2 and suggesting an identical substrate profile. A detailed description of the residue conservation within the respective binding pockets and comparison to other RND-transporters is provided in the supplementary S4 text 4 and table 6.

**Figure 5:**
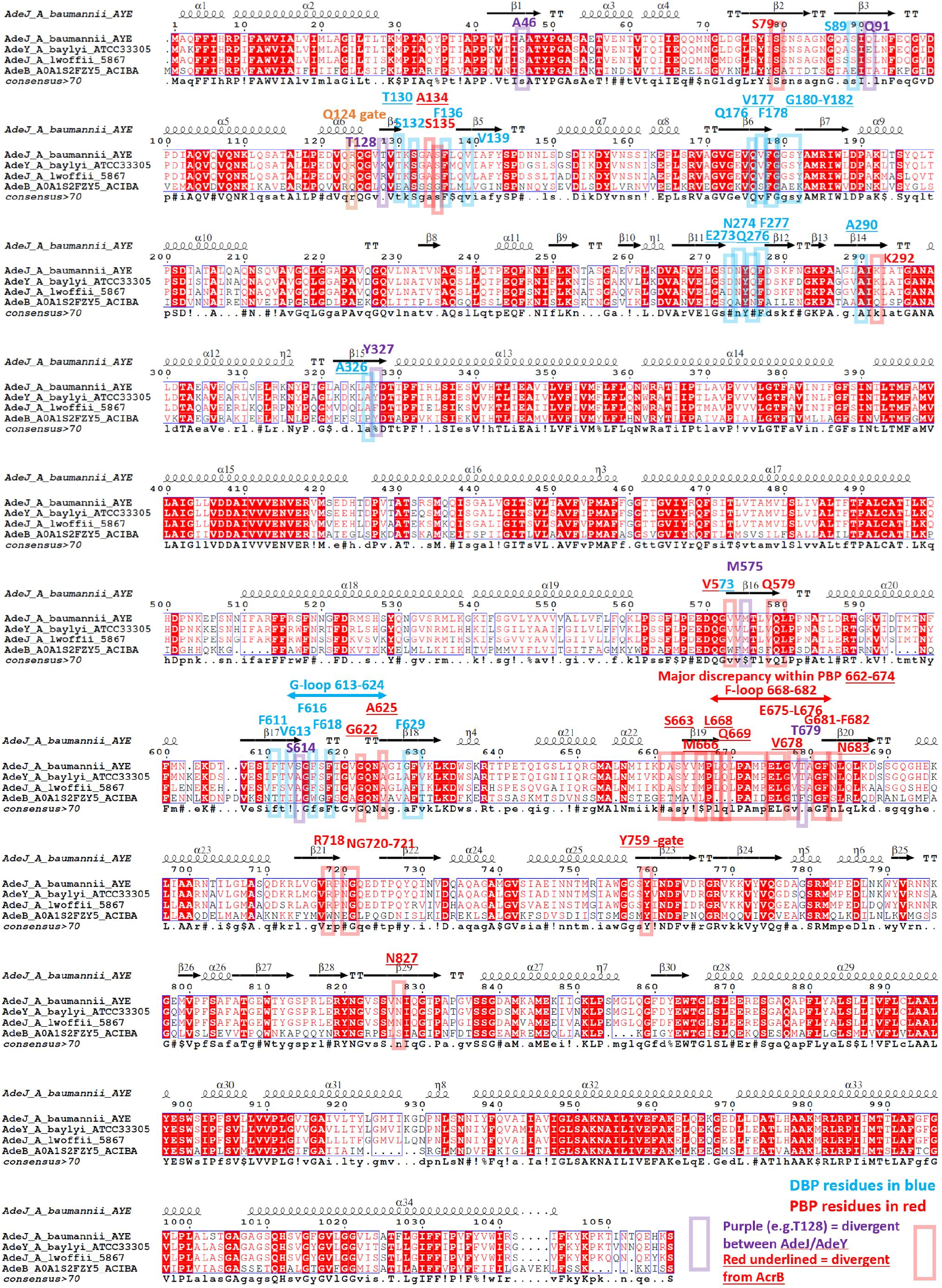
Sequence alignment of the AdeJ/Y and *Ab*AdeB, highlighting the key binding determinants in the DBP (blue boxes/font) and PBP (red boxes/font). The alignment shows the close relationship between AdeJ and AdeY, and positional equivalence between the members of AdeJ/Y clade. Residues within the drug-binding pockets that are divergent within the AdeJ/Y clade are highlighted in purple. Secondary structural elements derived from the experimental *Ab*AdeJ structure 7M4P.pdb. Consensus sequence displayed as a bottom row. Figure prepared with ESpript 3.0 (58).

The PBP is generally relatively conserved amongst the RND transporters, except for the residue range 660-688 (*Ab*AdeJ numbering), which forms the bottom section of the PBP and covers the so-called F-loop (figure 5). Strikingly, while this region displays near-complete conservation between the AdeJ and AdeY (the only minor exception of T679, which in *A. lwoffii* is represented by a conservative substitution to Serine), it shows major deviation from both the *Ab*AdeB (53,74,78) and *E. coli* and *Salmonella* AcrB (79,80). The AdeJ/Y PBP also features a diagnostic V573, which is strictly conserved amongst them, but not present in other RND-transporters, providing a clear differentiation of the AdeJ and AdeY pumps. Another prominently conserved residue within the AdeJ/Y-subfamily is found in the front of the PBP, corresponding to the R718 in *Ab*AdeJ (supplementary S4 figure 6, panel B). It is notable, that the R718 is conserved in AcrB/MtrD/MexB transporters, but not in the paralogous AdeB, which suggests closer relation of the AdeJ/Y to the former. Taken together the above conservation analysis suggests a common mode of substrate recognition in the PBP of the AdeJ and AdeY, once again highlighting their close relationship to each other.

The PBP and DBP are separated by the flexible G-loop (covering residue range 613-624 in *Ab*AdeJ), which contains a conserved phenylalanine (F618), which is involved in drug-binding in the DBP of both the newly-determined AdeJ fluorocycline-bound structures (53,81), and in AdeB (74). Amongst the AdeJ/Y the whole of the G-loop is strictly conserved, which yet again suggest similar binding properties in the upper part of the PBP and the front part of the DBP.

Moving to the DBP, the *Ab*AdeJ residues F136, F178, Y327, V613, F618, and F629 are universally conserved across AdeJ/Y, AcrB/MtrD and AdeB-transporters. Again, amongst the AdeJ/Y family members, there are very few substitutions (figure 6), but it is striking that three out of four that are present (A46, Q91 and T128), are clustered together at the back of the pocket (supplementary S4, figure 6, panel C), forming a plausible interaction site, hinted by the apparent covariation of the residues occupying positions 46 and 128. While in *A. baumannii* this pair is represented by A46/T128, in *A. lwoffii* and *A. baylyi* it is instead a S46 in combination with either R128 or K128 respectively. The last DBP residue to show variation within the AdeJ/Y subfamily is Y327 (F327 in *A. lwoffii*), and is found at the bottom of the pocket, opposite side across from the A46/T128 pair (supplementary S4, figure 6, panel C). Intriguingly, it is in direct contact with M575, itself a variable residue, forming the front of the PBP, and also interacting with the variable PBP residue T679 mentioned above.

**Figure 6:**
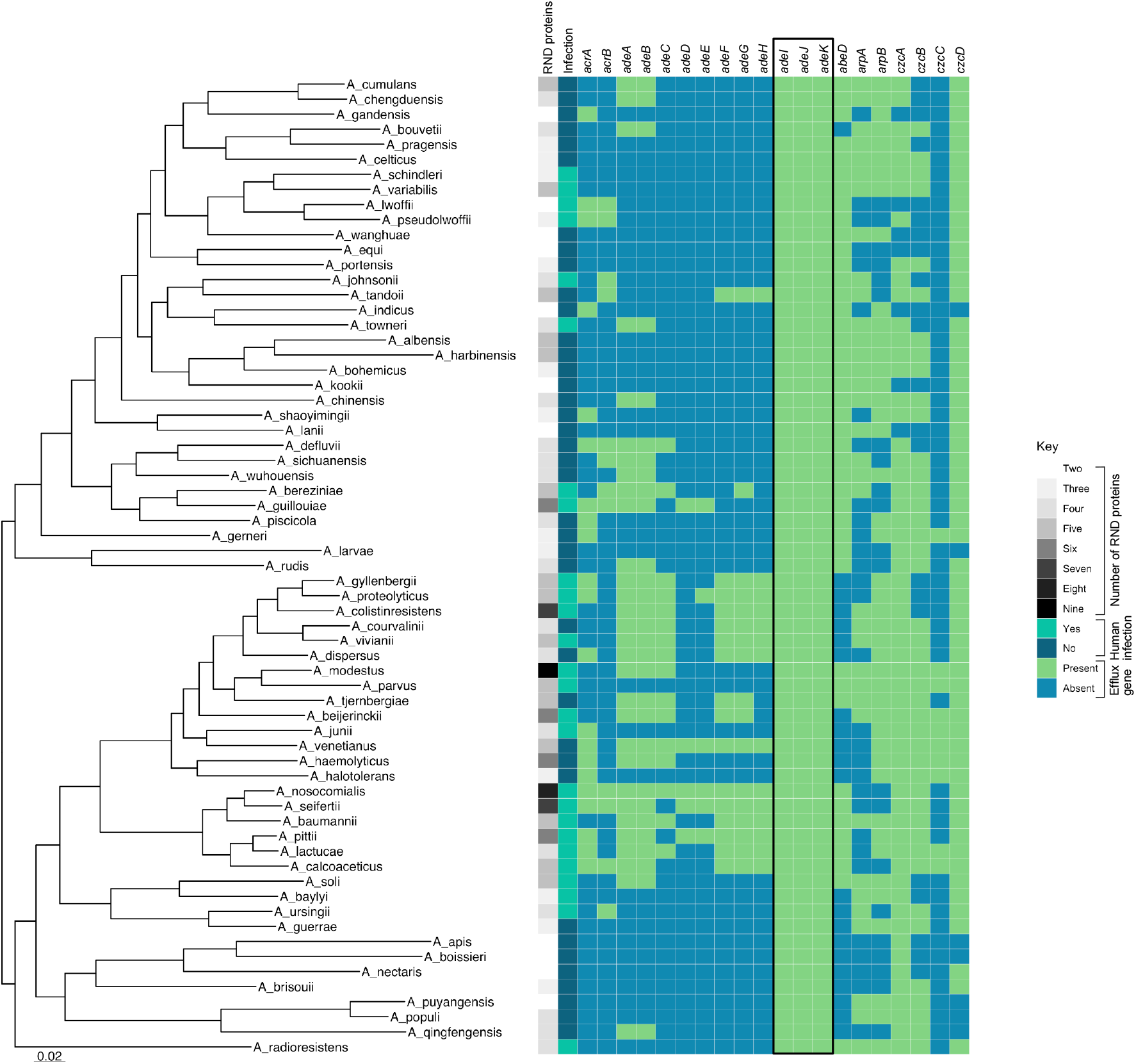
AdeIJK is the ancestral efflux pump in the *Acinetobacter* genus. Heatmap of *Acinetobacter* genus with characterised RND genes presence/absence and the number of RND proteins. For column 1, number of RNDs, the mean average of the total number of (both HME and HAE) RND efflux proteins for each species was determined, to 1 significant figure, where the greater the number of proteins, the darker grey the colour. For column 2, if a species has been shown in the literature to cause infection it is turquoise. Subsequent columns 3-22 are highlighted green if the efflux gene was found in the reference sequences using ABRicate. *adeIJK* and *adeXYZ* columns were amalgamated (black box) in this heatmap because they are the orthologs, for clarity.

In summary, the structural analysis of the AdeJ and AdeY transporters reveals that they share a number of distinguishing features that are common across them, but distinctive enough to set them apart as a separate group within the wider RND-family. These include major changes in the PBP, including the entire F-loop region, which is very much divergent from other RND transporters. At the same time, the overall high conservation of the DBP, combined with the presence of the R718 residue, helps to explain their identical substrate profile (Table 2), while suggesting a closer functional alignment to AcrB/MtrD/MexB transporters than to the paralogous AdeB.

Thus, based upon the genetic, structural and phenotypic similarities between *adeIJK* and *adeXYZ*, we propose than *adeXYZ* should be named *adeIJK* and have hence amalgamated the two systems in the heatmap in figure 6, which now shows the presence of *adeIJK* in every described *Acinetobacter* species.

## Discussion

The number of RND efflux genes per genome differs at both the genus and species level. Novel RND pumps are being characterised continually and it is likely there are many yet to be discovered. Due to the broad roles that RND pumps have within cells, fully understanding the complement of systems within a given species is important, especially when trying to understand MDR phenotypes in pathogens. We have presented a simple way to screen for HAE and HME RND proteins from bacterial genomes based upon conserved residues, which highlighted both known RND proteins and some uncharacterised ones in a variety of Gram-negative species. There are some limitations to the method as very divergent systems, such as ArpB, may not be detected. A combination approach of searching for known efflux genes and using the conserved residue RND protein search provides the best insight into the complement of RND systems in a given species.

The number of RND proteins differs between species of *Acinetobacter* and also within species. In *Acinetobacter*, there were between two and nine RND proteins in any given genome sequence, where *A. nosocomialis* and *A. modestus* had the most RND genes. Species that cause human infection encoded more RND proteins. Previous work has shown that by over-expressing the RND genes *adeABC, A. baumannii* is more virulent in the lungs of a mouse (82) and the role of RND pumps in infections has also been noted in other Gram-negative bacteria including *N. gonorrhoeae* (MtrCDE), *Salmonella enterica* (AcrAB-TolC), *Pseudomonas aeruginosa* (MexAB-OprM*), Campylobacter jejuni* (CmeABC) and *Vibrio cholerae* (VexB/D/K) (83–87). Subsequent work is needed to fully elucidate if there is a link between the number of RND proteins and a species’ ability to cause infection in humans across the *Acinetobacter* genus. In this study sixty-four *Acinetobacter* species were characterised at the time the analyses were done, however since then further *Acinetobacter* species have been described. Additionally, when looking at RND proteins within the *A. baumannii* species, almost all sequences had the five characterised RND proteins but there was some variation, with some sequences lacking AdeB, which has been seen previously in *A. baumannii* isolates (24,33).

The RND system *adeIJK* was found to be present across *Acinetobacter* and isolates without it encoded *adeXYZ*, which led us to investigate if these systems are actually the same. The MIC data shows that both AdeIJK and AdeXYZ can export the same compounds including tetracycline, chloramphenicol and clindamycin which have previously been identified as substrates as AdeIJK (63). Exact values were not directly comparable to previous studies as different expression systems were used. The pVRL vectors are high copy number plasmids explaining why some MICs increased above that of the wild-type strain. Very high levels of expression of AdeIJK have previously been shown to be toxic in both *A. baumannii* and *E. coli* (23) and correspondingly we were unable to successfully express AdeIJK in *E. coli* despite testing a range of vectors.

Homology modelling supported the data suggesting identical substrate profiles as the structure of AdeJ and AdeY, and in particular their binding pockets, were highly similar.

Our structural analysis also confirmed the common structural features of AdeJ/Y, which justifies isolating them as a separate subfamily of RND transporters. Indeed, while the overall architecture, and correspondingly the structure of the drug-binding pockets is conserved across AdeJ/Y, they show unique features clearly distinguishing them even from the closely related RND relatives. In interests of space, the detailed discussion of residue conservation is provided in supplementary S4, text 4, but a few noteworthy elements are highlighted below.

The differentiating features of AdeJ/Y include the organisation of the principal drug-binding pockets, with, in particular the base of the PBP displaying clear differences from other members of RND family, including the so-called F-loop (figure 6, supplementary S4, figure 6B). Indeed, the F-loop is different not only between the AdeJ/Y and AdeB, but also between AdeJ/Y and AcrB/MtrD. It seems likely this discrepancy may be contributing to the differential substrate efflux efficiencies between AdeB and AdeJ/Y reported earlier (88).

The PBP also features a diagnostic V573, which is only present within the AdeJ/Y subfamily, providing differentiation from other transporters. In addition, AdeJ/Y also display some hybrid features, linking them to the canonical MDR transporters. These include the presence of an arginine (R718 in AdeJ/Y), which forms front part of the PBP (substrate channel 2 exit) (77,89), and which is conserved across AcrB/MtrD/MexB (R717 in AcrB) (57,90,91), where it has been implicated in the binding of the macrolides and rifamicyns (57,95,96). This critical residue is not conserved in AdeB however, suggesting that AdeJ/Y likely process their substrates more similarly to the AcrB/MtrD than to AdeB. This is further supported by our analysis of DBP, where only 4 residues have been shown to have limited variability within the analysed AdeJ/Y structures. As mentioned above, three out of four variable residues (A46, Q91 and T128) are clustered together at the back of the pocket and display positional covariation (supplementary S4, figure 6, panel C), suggesting functional interaction between them and formation of a plausible interaction site, which could be responsible for distinctive DBP ligand coordination and warrant further investigation beyond the remit of the current study. The additional conservation of the residues within the DBP, including the critical ligand-binding residues such as AcrB F610 (corresponding to *Ab*AdeJ F611), across AcrB/MtrD/MexB, but not AdeB once again suggests closer relation of AdeJ/Y to the former.

The clear distinction of AdeJ/Y from AdeB, and the closeness of AdeJ/Y instead to AcrB/MtrD/MexB, suggest that the AdeJ/Y represent the basal efflux pumps within the genus, while AdeB paralogues may have been acquired within the *Acinetobacter* genus.

Together, the genetic, structural and phenotypic data presented shows that AdeIJK and AdeXYZ are in fact divergent orthologues of the same system and we propose than *adeXYZ* should be named *adeIJK*. As shown in figure 6, this highlights the presence of *adeIJK* in every *Acinetobacter* species studied to date indicating it is under high selection pressure providing an important function. The reason for the presence of *adeIJK* across all species in the genus might be due to the wide roles that *adeIJK* carries out in *Acinetobacter*, for example its ability to protect against antibacterial host-associated fatty acids and modulate the bacterial cell membrane (88). Furthermore, *adeIJK* plays a role in virulence, biofilm formation, surface motility and can export a broad range of compounds, including clinically relevant antibiotics, providing intrinsic resistance (82,92,93). It is therefore possible to say that *adeIJK* is the defining *Acinetobacter* pump, and to belong to the genus a species will encode a version of *adeIJK*. Important disease-causing *Acinetobacter* may also encode *adeABC*, which when overexpressed increases antibiotic resistance in *A. baumannii* (8). Therefore, it is advantageous in some species to have the combination of *adeIJK*, providing intrinsic resistance, and *adeABC*, synergistically providing even higher resistance. It is uncommon for *adeIJK* to be overexpressed, implying the functions it carries out are important and need to be tightly regulated (8,23).

## Supporting information

Supplemental 4 - structure

pVRL2 AdeXYZ fasta

pVRL2 AdeIJK fasta

## Author Statements

### Author contributions

E. M. D. and J.M.A.B. conceptualised and designed the study. E.M.D. created the BLASTp database, carried out genomic analyses and phenotypic work. V. N. B. provided protein modelling, analysis and discussion on structure. S. D. contributed to the gubbins recombination analysis. The manuscript was written by E. M. D, J. M. A. B and V. N. B, with input from A. M and S.D.

### Ethical approval

This study did not require ethical approval.

## Acknowledgements

The authors thank Massimiliano Lucidi for providing the cloning plasmid pVRL2 for this study, Ayush Kumar for providing *A. baumannii* ATCC 17978 *ΔadeAB ΔadeFGH ΔadeIJK* and Laura Piddock for providing *A. baumannii* ATCC 17978.

