## Supplemental 1 - figures for "RND pumps across the *Acinetobacter* genus; AdeIJK is the ancestral efflux system"

### Supplementary S1:

**Supplementary S1, figure 1**: HAE protein alignment for BLASTp.

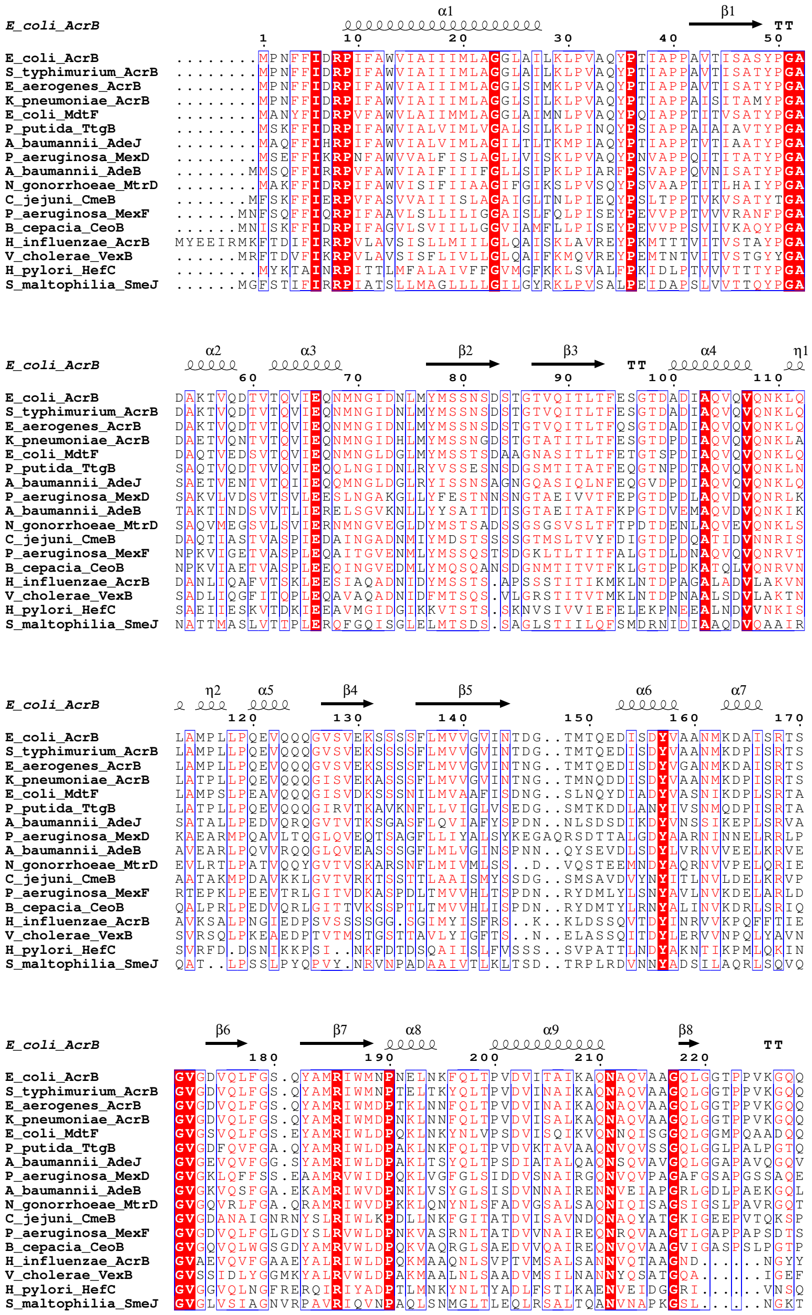

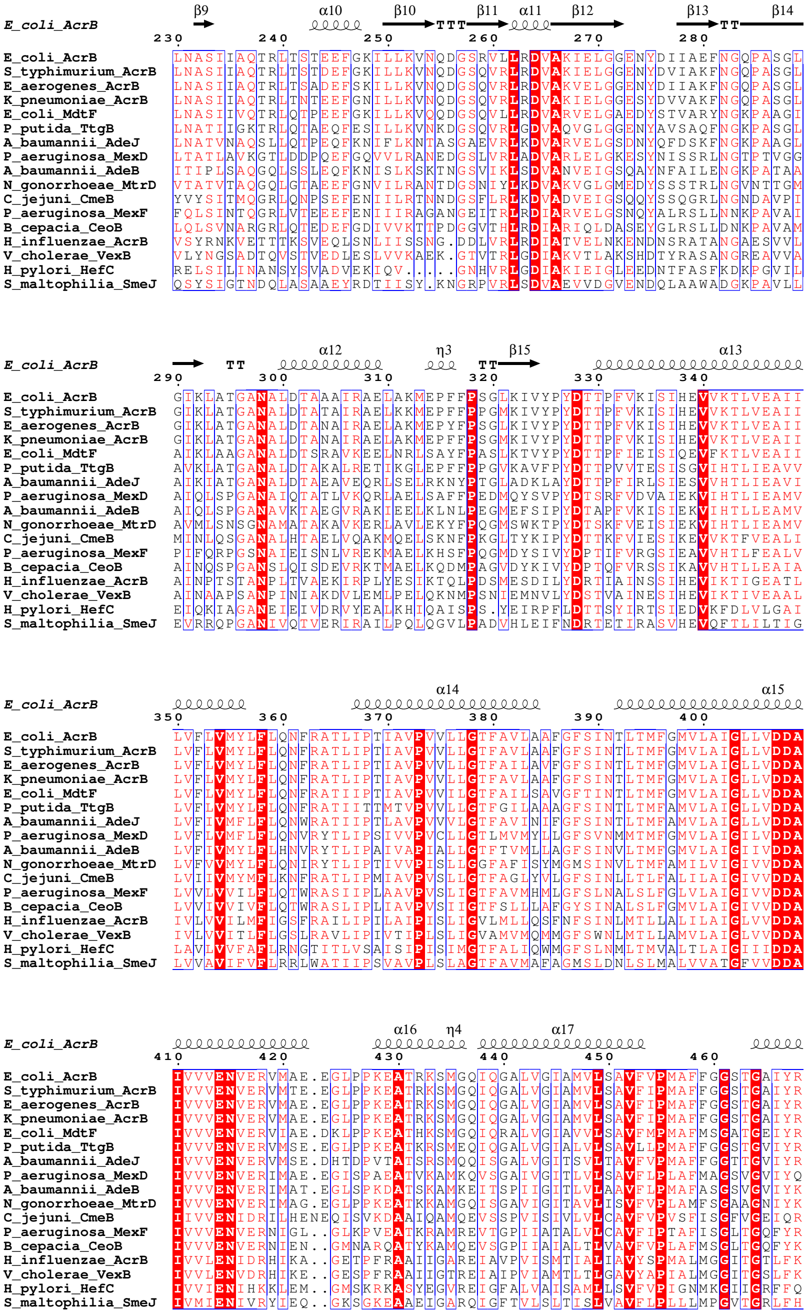

E_coli_AcrB
S_typhimurium_AcrB
E_aerogenes_AcrB
K_pneumoniae_AcrB
E_coli_MdtF
P_putida_TtgB
A_baumannii_AdeJ
P_aeruginosa_MexD
A_baumannii_AdeB
N_gonorrhoeae_MtrD
C_jejuni_CmeB
P_aeruginosa_MexF
B_cepacia_CeoB
H_influenzae_AcrB
V_cholerae_VexB
H_pylori_HefC
S_maltophilia_SmeJ
β9 α10 β10 β11 α11 β12 β13 β14
α12 η3 β15 α13
α14 α15
α16 η4 α17

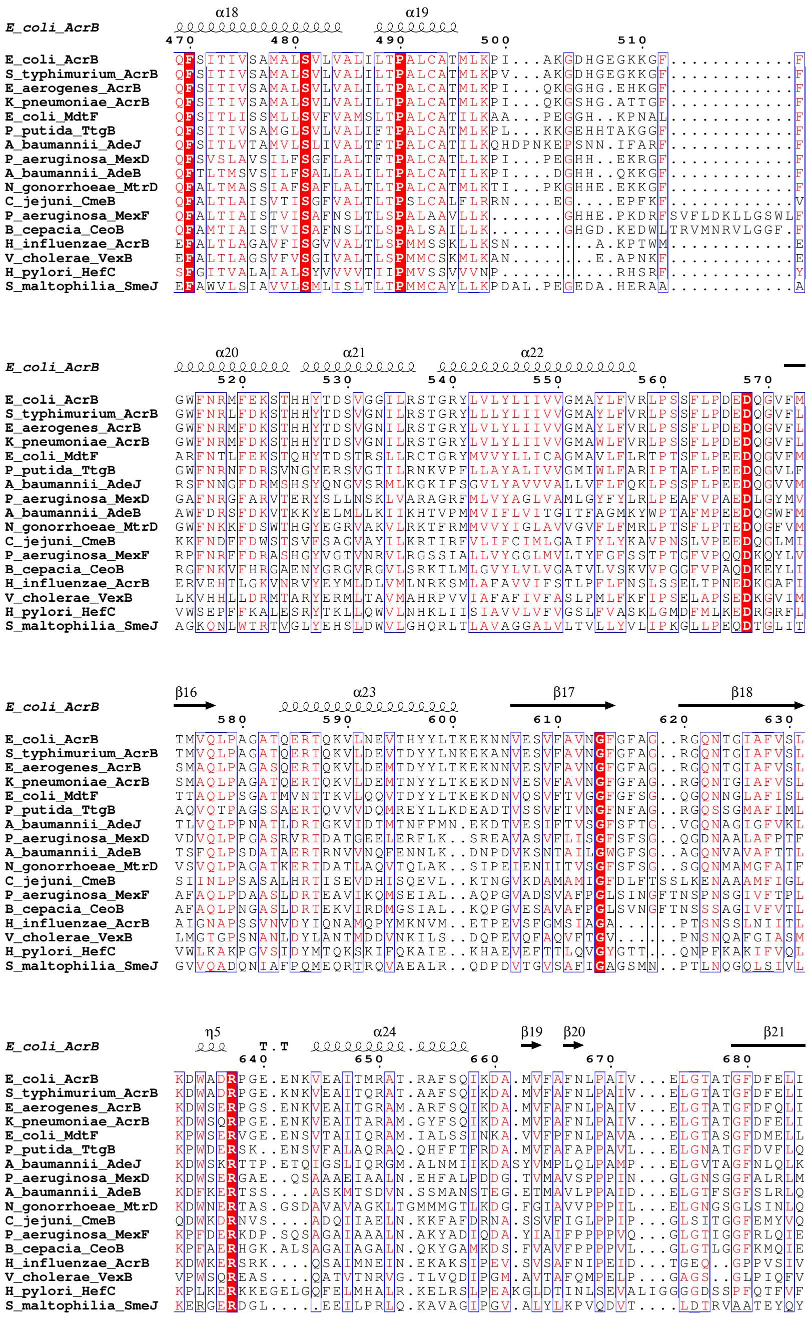
E_coli_AcrB α18 α19
470 480 490 500 510
E_coli_AcrB QFSITIVSAMALSVLVALILTPALCATMLKPI...AKGDHGEGKKGF...........F
S_typhimurium_AcrB QFSITIVSAMALSVLVALILTPALCATMLKPV...AKGDHGEGKKGF...........F
E_aerogenes_AcrB QFSITIVSAMALSVLVALILTPALCATMLKPI...QKGGHG.EHKGF...........F
K_pneumoniae_AcrB QFSITIVSAMALSVLVALILTPALCATMLKPI...QKGSHG.ATTGF...........F
E_coli_MdtF QFSITLISSMLLSVFVAMSLTPALCATILKAA...PEGGH..KPNAL...........F
P_putida_TtgB QFSITIVSAMGLSVLVALIFTPALCATMLKPL...KKGEHHTAKGGF...........F
A_baumannii_AdeJ QFSITLVTAMVLSLIVALTFTPALCATILKQHDPNKEPSNN.IFARF...........F
P_aeruginosa_MexD QFSVSLAVSILFSGFLALTFTPALCATLLKPI...PEGHH..EKRGF...........F
A_baumannii_AdeB QFTLTMSVSILFSALLALILTPALCATILKPI....DGHH..QKKGF...........F
N_gonorrhoeae_MtrD QFALTMASSIAFSAFLALTLTPALCATMLKTI...PKGHHE.EKKGF...........F
C_jejuni_CmeB QFALTLAISVTISGFVALTLTPSLCALFLRRN....EG....EPFKF...........V
P_aeruginosa_MexF QFALTIAISTVISAFNSLTLSPALAAVLLK.......GHHE.PKDRFSVFLDKLLGSWLF
B_cepacia_CeoB QFAMTIAISTVISAFNSLTLSPALSAILLK.......GHGD.KEDWLTRVMNRVLGGF.F
H_influenzae_AcrB EFALTLAGAVFISGVVALTLSPMMSSKLLKSN.......A.KPTWM...........E
V_cholerae_VexB EFALTLAGSVFVSGIVALTLSPMMCSKMLKAN.......E.APNKF...........E
H_pylori_HefC SFGITVALAIALSYVVVVTIIPMVSSVVNP..........RHSRF...........Y
S_maltophilia_SmeJ EFAWVLSIAVVLSMLISLTLTPMMCAYLLKPDAL.PEGEDA.HERAA...........A

E_coli_AcrB α20 α21 α22
520 530 540 550 560 570
E_coli_AcrB GWFNRMFEKSTHHYTDSVGGILRSTGRYLVLYLIIVVGMAYLFVRLPSSFLPDEDQGVFM
S_typhimurium_AcrB GWFNRLFDKSTHHYTDSVGNILRSTGRYLLLYLIIVVGMAYLFVRLPSSFLPDEDQGVFL
E_aerogenes_AcrB GWFNRMFDKSTHHYTDSVGNILRSTGRYLVLYLIIVVGMAYLFVRLPSSFLPDEDQGVFL
K_pneumoniae_AcrB GWFNRMFDKSTHHYTDSVGNILRSTGRYLVLYLIIVVGMAWLFVRLPSSFLPDEDQGVFL
E_coli_MdtF ARFNTLFEKSTQHYTDSTRSLLRCTGRYMVVYLLICAGMAVLFLRTPTSFLPEEDQGVFM
P_putida_TtgB GWFNRNFDRSVNGYERSVGTILRNKVPFLLAYALIVVGMIWLFARIPTAFLPEEDQGVLF
A_baumannii_AdeJ RSFNNGFDRMSHSYQNGVSRMLKGKIFSGVLYAVVVALLVFLFQKLPSSFLPEEDQGVVM
P_aeruginosa_MexD GAFNRGFARVTERYSLLNSKLVARAGRFMLVYAGLVAMLGYFYLRLPEAFVPAEDLGYMV
A_baumannii_AdeB AWFDRSFDKVTKKYELMLLKIIKHTVPMMVIFLVITGITFAGMKYWPTAFMPEEDQGWFM
N_gonorrhoeae_MtrD GWFNKKFDSWTHGYEGRVAKVLRKTFRMMVVYIGLAVVGVFLFMRLPTSFLPTEDQGFVM
C_jejuni_CmeB KKFNDFFDWSTSVFSAGVAYILKRTIRFVLIFCIMLGAIFYLYKAVPNSLVPEEDQGLMI
P_aeruginosa_MexF RPFNRFFDRASHGYVGTVNRVLRGSSIALLVYGGLMVLTYFGFSSTPTGFVPQQDKQYLV
B_cepacia_CeoB RGFNKVFHRGAENYGRGVRGVLSRKTLMLGVYLVLVGATVLVSKVVPGGFVPAQDKEYLI
H_influenzae_AcrB ERVEHTLGKVNRVYEYMLDLVMLNRKSMLAFAVVIFSTLPFLFNSLSSELTPNEDKGAFI
V_cholerae_VexB LKVHHLLDRMTARYERMLTAVMAHRPVVIAFAFIVFASLPMLFKFIPSELAPSEDKGVIM
H_pylori_HefC VWSEPFFKALESRYTKLLQWVLNHKLIISIAVVLVFVGSLFVASKLGMDFMLKEDRGRFL
S_maltophilia_SmeJ AGKQNLWTRTVGLYEHSLDWVLGHQRLTLAVAGGALVLTVLLYVLIPKGLLPEQDTGLIT

E_coli_AcrB β16 α23 β17 β18
580 590 600 610 620 630
E_coli_AcrB TMVQLPAGATQERTQKVLNEVTHYYLTKEKNNVESVFAVNGFGFAG..RGQNTGIAFVSL
S_typhimurium_AcrB TMVQLPAGATQERTQKVLDEVTDYYLNKEKANVESVFAVNGFGFAG..RGQNTGIAFVSL
E_aerogenes_AcrB SMAQLPAGASQERTQKVLDEMTDYYLTKEKNNVESVFAVNGFGFAG..RGQNTGIAFVSL
K_pneumoniae_AcrB SMAQLPAGATQERTQKVLDEMTNYYLTKEKDNVESVFAVNGFGFAG..RGQNTGIAFVSL
E_coli_MdtF TTAQLPSGATMVNTTKVLQQVTDYYLTKEKDNVQSVFTVGGFGFSG..QGQNNGLAFISL
P_putida_TtgB AQVQTPAGSSAERTQVVVDQMREYLLKDEADTVSSVFTVNGFNFAG..RGQSSGMAFIML
A_baumannii_AdeJ TLVQLPPNATLDRTGKVIDTMTNFFMN.EKDTVESIFTVSGFSFTG..VGQNAGIGFVKL
P_aeruginosa_MexD VDVQLPPGASRVRTDATGEELERFLK..SREAVASVFLISGFSFSG..QGDNAALAFPTF
A_baumannii_AdeB TSFQLPSDATAERTRNVVNQFENNLK..DNPDVKSNTAILGWGFSG..AGQNVAVAFTTL
N_gonorrhoeae_MtrD VSVQLPAGATKERTDATLAQVTQLAK..SIPEIENIITVSGFSFSG..SGQNMAMGFAIF
C_jejuni_CmeB SIINLPSASALHRTISEVDHISQEVL..KTNGVKDAMAMIGFDLFTSSLKENAAAMFIGL
P_aeruginosa_MexF AFAQLPDAASLDRTEAVIKQMSEIAL..AQPGVADSVAFPGLSINGFTNSPNSGIVFTPL
B_cepacia_CeoB AFAQLPNGASLDRTEKVIRDMGSIAL..KQPGVESAVAFPGLSVNGFTNSSSAGIVFVTL
H_influenzae_AcrB AIGNAPSSVNVDYIQNAMQPYMKNVM..ETPEVSFGMSIAGA......PTSNSSLNIITL
V_cholerae_VexB LMGTGPSNANLDYLANTMDDVNKILS..DQPEVQFAQVFTGV......PNSNQAFGIASM
H_pylori_HefC VWLKAKPGVSIDYMTQKSKIFQKAIE..KHAEVEFTTLQVGYGTT...QNPFKAKIFVQL
S_maltophilia_SmeJ GVVQADQNIAFPQMEQRTRQVAEALR..QDPDVTGVSAFIGAGSMN..PTLNQGQLSIVL

E_coli_AcrB η5 T.T α24 β19 β20 β21
640 650 660 670 680
E_coli_AcrB KDWADRPGE.ENKVEAITMRAT.RAFSQIKDA.MVFAFNLPAIV...ELGTATGFDFELI
S_typhimurium_AcrB KDWADRPGE.KNKVEAITQRAT.AAFSQIKDA.MVFAFNLPAIV...ELGTATGFDFELI
E_aerogenes_AcrB KDWSERPGS.ENKVEAITGRAM.ARFSQIKDA.MVFAFNLPAIV...ELGTATGFDFQLI
K_pneumoniae_AcrB KDWSQRPGE.ENKVEAITARAM.GYFSQIKDA.MVFAFNLPAIV...ELGTATGFDFELI
E_coli_MdtF KPWSERVGE.ENSVTAIIQRAM.IALSSINKA.VVFPFNLPAVA...ELGTASGFDMELL
P_putida_TtgB KPWDERSK..ENSVFALAQRAQ.QHFFTFRDA.MVFAFAPPAVL...ELGNATGFDVFLQ
A_baumannii_AdeJ KDWSKRTTP.ETQIGSLIQRGM.ALNMIIKDASYVMPLQLPAMP...ELGVTAGFNLQLK
P_aeruginosa_MexD KDWSERGAE..QSAAAEIAALN.EHFALPDDG.TVMAVSPPPIN...GLGNSGGFALRLM
A_baumannii_AdeB KDFKERTSS....ASKMTSDVN.SSMANSTEG.ETMAVLPPAID...ELGTFSGFSLRLQ
N_gonorrhoeae_MtrD KDWNERTAS.GSDAVAVAGKLTGMMMGTLKDG.FGIAVVPPPIL...ELGNGSGLSINLQ
C_jejuni_CmeB QDWKDRNVS....ADQIIAELN.KKFAFDRNA.SSVFIGLPPIP...GLSITGGFEMYVQ
P_aeruginosa_MexF KPFDERKDP.SQSAGAIAAALN.AKYADIQDA.YIAIFPPPPVQ...GLGTIGGFRLQIE
B_cepacia_CeoB KPFAERHGK.ALSAGAIAGALN.QKYGAMKDS.FVAVFPPPPVL...GLGTLGGFKMQIE
H_influenzae_AcrB KDWKERSRK....QSAIMNEIN.EKAKSIPEV.SVSAFNIPEID...TGEQ..GPPVSIV
V_cholerae_VexB VPWSQREAS....QATVTNRVG.TLVQDIPGM.AVTAFQMPELP...GAGS..GLPIQFV
H_pylori_HefC KPLKERKKEGELGQFELMHALR.KELRSLPEAKGLDTINLSEVALIGGGGDSSPFQTFVF
S_maltophilia_SmeJ KERGERDGL.....EEILPRLQ.KAVAGIPGV.ALYLKPVQDVT....LDTRVAATEYQY

```
                            α25                        β22      β23       α26
E_coli_AcrB         ► ooooooooooooooooo            →        →        oooooo
                      690       700       710       720       730       740
E_coli_AcrB         DQAGLGHEKLTQARNQLLAEAAKHPDML...TSVRPNGLEDTPQFKIDIDQEKAQALGVS
S_typhimurium_AcrB  DQAGLGHEKLTQARNQLFGEVAKYPDLL...VGVRPNGLEDTPQFKIDIDQEKAQALGVS
E_aerogenes_AcrB    DQGGLGHEKLTQARNQLFGMVAQHPDLL...VGVRPNGLEDTPQFKIDIDQEKAQALGVS
K_pneumoniae_AcrB   DQGGLGHEKLTQARNQLFGMVAQHPDVL...TGVRPNGLEDTPQFKIDIDQEKAQALGVS
E_coli_MdtF         DNGNLGHEKLTQARNELLSLAAQSPNQV...TGVRPNGLEDTPMFKVNVNAAKAEAMGVA
P_putida_TtgB       DRGGVGHAKLMEARNQFLAKAAQSKIL....SAVRPNGLNDEPQYQLTIDDERASALGVT
A_baumannii_AdeJ    DSSGQGHEKLIAARNTILGLASQDKRL....VGVRPNGQEDTPQYQINVDQAQAGAMGVS
P_aeruginosa_MexD   DRSGVGREALLQARDTLLGEIQTNPKF....LYAMMEGLAEAPQLRLLIDREKARALGVS
A_baumannii_AdeB    DRANLGMPALLAAQDELMAMAAKNKKF....YMVWNEGLPQGDNISLKIDREKLSALGVK
N_gonorrhoeae_MtrD  DRNNTGHTGIAGEGNELIQKMRASGLFDP..STVRAGGLEDSPQLKIDINRAAAAAQGIS
C_jejuni_CmeB       NKSGKSYDEIQKDVNKLVAAANQRKEL....SRVRTTLDTTFPQYKLIIDRDKLKHYNLN
P_aeruginosa_MexF   DRGNQGYEELFKQTQNIITKARALPELEP..SSVFSSYQVNVPQIDADIDREKAKTHGVA
B_cepacia_CeoB      DRGAVGYAKLSDATNDFIKRAQQAPEL....GPLFTSYQINVPQLNVDLDRVKAKQLGVP
H_influenzae_AcrB   LKTAQDYKSLANTAEKFLSAMKASGKF....IYTNLDLTYDTAQMTISVDKEKAGTYGIT
V_cholerae_VexB     ITTPSNFESLFTIATDVLTEVKANPMF....VYSDLDLNFDSATMKINIDKDKAGAYGVT
H_pylori_HefC       SHS...QEAVDKSVENLRKFLLESPELKGKVESYHTSTSESQPQLQLKILRQNANKYGVS
S_maltophilia_SmeJ  SLSDVDSATVATQATRLTEALRKRPEL....ADVDNNLSNQGRALELNIDRDKASVLGVP

                          α27           β24       β25        η6    η7    β26
E_coli_AcrB         oooooooooooo     →TT→                   ooo  oooo   →    T.......
                          750       760       770       780       790
E_coli_AcrB         INDINTTLGAAW.GGSYVNDFIDRGRVKKVYVMSEAKYRMLPDDIGDWYVRAA.......
S_typhimurium_AcrB  ISDINTTLGAAW.GGSYVNDFIDRGRVKKVYVMSEAKYRMLPDDINDWYVRGS.......
E_aerogenes_AcrB    ISDINTTLGAAW.GGSYVNDFIDRGRVKKVYVMSEAKYRMLPEDIGNWYVRGS.......
K_pneumoniae_AcrB   ISDINTTLGAAW.GGSYVNDFIDRGRVKKVYIMSEAKYRMLPEDIGKWYVRGS.......
E_coli_MdtF         LSDINQTISTAF.GSSYVNDFLNQGRVKKVYVQAGTPFRMLPDNINQWYVRNA.......
P_putida_TtgB       IADINNTLSIAL.GASYVNDFIDRGRVKKVYIQGEPSARMSPEDLQKWYVRNG.......
A_baumannii_AdeJ    IAEINNTMRIAW.GGSYINDFVDRGRVKKVYVQGDAGSRMMPEDLNKWYVRNN.......
P_aeruginosa_MexD   FETISGTLSAAF.GSEVINDFTNAGRQQRVVIQAEQGNRMTPESVLELYVPNA.......
A_baumannii_AdeB    FSDVSDIISTSM.GSMYINDFPNQGRMQQVIVQVEAKSRMQLKDILNLKVMGS.......
N_gonorrhoeae_MtrD  FADIRTALASAL.SSSYVSDFPNQGRLQRVMVQADEDARMQPADILNLTVPNK.......
C_jejuni_CmeB       MQDVFNTMNATI.GTYYVNDFSMLGKNFQVNIRAKGDFRNTQDALKNIFVRSN.......
P_aeruginosa_MexF   ISDIFDTLQVYL.GSLYANDFNRFGRTYQVNVQAEQQFRLEPEQIGQLKVRNN.......
B_cepacia_CeoB      VTDVFNTMQVYL.GSLYVNDFNRFGRVYQVRVQADAPFRQRADDILQLKTRND.......
H_influenzae_AcrB   MQQISNTLGSFL.SGATVTRVDVDGRAYKVISQVKRDDRLSPESFQNYYLTAS.......
V_cholerae_VexB     MQDIGITLSTMM.ADGYVNRIDLNGRSYEVIPQVERKWRLNPESMNSYYVRAA.......
H_pylori_HefC       AQTIGSVVSSAFSGTSQASVFKEDGKEYDMIIRVPDDKRVSVEDIKRLQVRNK.......
S_maltophilia_SmeJ  MQTIDDTLYDAF.GQRQISTIFTELNQYRVVLEVAPEFRTSTALMEQLAVASNGAGALTG

                                                      β27 η8    β28      β29
E_coli_AcrB         ...............................T   → ooo    →        →TT—
                                                      800       810       820
E_coli_AcrB         ...............................DGQMVPFSAFSSSRWEYGSPRLERYNGLPS
S_typhimurium_AcrB  ...............................DGQMVPFSAFSSSRWEYGSPRLERYNGLPS
E_aerogenes_AcrB    ...............................DGQMVPFSAFSTSHWEYGSPRLERYNGLPS
K_pneumoniae_AcrB   ...............................DGQMVPFSAFSTSRWEYGSPRLERYNGLPS
E_coli_MdtF         ...............................SGTMAPLSAYSSTEWTYGSPRLERYNGIPS
P_putida_TtgB       ...............................AGEMVPFSSFAKGEWTYGSPKLSRYNGVEA
A_baumannii_AdeJ    ...............................KGEMVPFSAFATGEWTYGSPRLERYNGVSS
P_aeruginosa_MexD   ...............................AGNLVPLSAFVSVKWEEGPVQLVRYNGYPS
A_baumannii_AdeB    ...............................SGQLVSLSEVVTPQWNKAPQQYNRYNGRPS
N_gonorrhoeae_MtrD  ...............................SGVAVPLSTIATVSWENGTEQSVRFNGYPS
C_jejuni_CmeB       ...............................DGKMIPLDSFLTLQRSSGPDDVKRFNLFPA
P_aeruginosa_MexF   ...............................LGEMVPLASFIKVSDTSGPDRVMHYNGFIT
B_cepacia_CeoB      ...............................KGEMVPLSSLVTVTPTFGPEMVVRYNGYTA
H_influenzae_AcrB   ...............................NGQSVPLSSVISMKLETQPTSLPRFSQLNS
V_cholerae_VexB     ...............................DGKVIPLGSLVTIDVVAEPRSLPHFNQLNS
H_pylori_HefC       ...............................YDKLMFLDALVEITETKSPSSISRYNRQRS
S_maltophilia_SmeJ  TNATSFGQVTSSNSSTATGIGAQNTGITVGAGNIIPLSALAEGKVSSAPLVVSHQQQLPA

                    β30               α28                 β31
E_coli_AcrB         →            ooooooooooooo       T....T→                oooo
                          830       840       850       860       870
E_coli_AcrB         MEILGQA..APGKSTGEAMELMEQLASK..LPT....GVGYDWTGMSYQERLSGNQAPSL
S_typhimurium_AcrB  MEILGQA..APGKSTGEAMAMMEELASK..LPS....GIGYDWTGMSYQERLSGNQAPAL
E_aerogenes_AcrB    MEILGQA..APGRSTGEAMAMMEQLASK..LPS....GVGYDWTGMSYQERLSGNQAPAL
K_pneumoniae_AcrB   LEILGQA..APGKSTGEAMALMEELAGK..LPS....GIGYDWTGMSYQERLSGNQAPAL
E_coli_MdtF         MEILGEA..AAGKSTGDAMKFMADLVAK..LPA....GVGYSWTGLSYQEALSSNQAPAL
P_putida_TtgB       MEILGAP..APGYSTGEAMAEVERIAGE..LPS....GIGFSWTGMSYEEKLSGSQMPAL
A_baumannii_AdeJ    VNIQGTP..APGVSSGDAMKAMEEIIGK..LPSMGLQGFDYEWTGLSLEERESGAQAPFL
P_aeruginosa_MexD   IRIVGDA..APGFSTGEAMAEMERLASQ..LPA....GIGYEWTGLSYQEKVSAGQATSL
A_baumannii_AdeB    LSIAGIP..NFDTSSGEAMREMEQLIAK..LPK....GIGYEWTGISLQEKQSESQMAFL
N_gonorrhoeae_MtrD  MKLSASP..ATGVSTGQAMEAVQKMVDE..LGG....GYSFEWGGQSSEEAKGGSQTLIL
C_jejuni_CmeB       AQVQGQP..APGYTSGQAIEAIAQVAKE.TLGD....DYSIAWSGSAYQEVSSKGTASYA
P_aeruginosa_MexF   AELNGAP..AAGYSSGQAQAAIEKLLKE.ELPN....GMTYEWTELTYQQILAGNTALFV
B_cepacia_CeoB      ADINGGP..APGFSSGQAQAAVERIAHE.TLPR....GVRFEWTDLTYQQILAGDSAMWV
H_influenzae_AcrB   AEISAVP..MPGTSSGDAIAWLQQQAND.NLPQ....GYTYDFKSEARQLVQEGNALTTT
V_cholerae_VexB     ATVGAVP..APGTAMGDAINWFENLASS.KLPK....GYSHDYMGEARQYVTEGSALYAT
H_pylori_HefC       VTVLAEPNRNAGVSLGEILTQVSKNTKE.WLVE....GANYRFTGEADNAKESNGEFLIA
S_maltophilia_SmeJ  VTVSFNV..APGYSLSEAVRAIQETKDSLDMPA....HLHAEFIGKAAEFTGSQTDVVWL
```

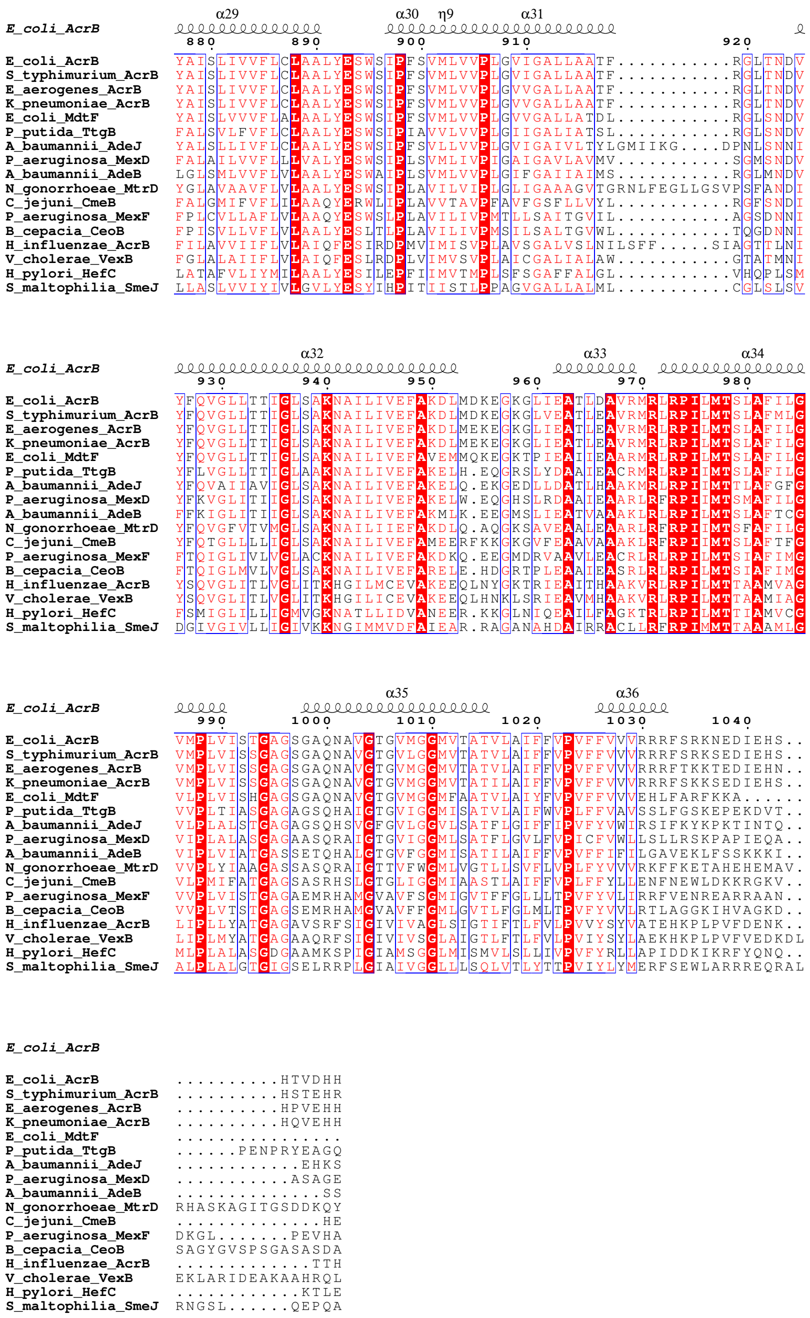


RND HAE proteins from different Gram-negative bacteria were aligned using MAFFT (v.7) against *E. coli* AcrB (P31224) reference sequence. The sequences aligned were *S. enterica* Typhimurium AcrB (Q8ZRA7), *E. aerogenes* AcrB (Q9AEG1), *K. pneumoniae* AcrB (W9B4M6), *E. coli* MdtF (P37637), *P. putida* TtgB (O52248), *A. baumanii* AdeJ (Q2FD94), *P. aeruginosa* MexD (Q9HVI9), *A. baumannii* AdeB (Q2FD70), *N. gonorrhoeae* MtrD (Q51073), *C. jejuni* CmeB (Q8RTE4), *P. aeruginosa* MexF (Q9I0Y8), *B. cepacia* CeoB (B4EJQ0), *H. influenzae* AcrB (A0A224AVD5), *V. cholerae* VexB (A0A085RXY1), *H. pylori* HefC (B6JLJ0), *S. maltophilia* SmeJ (A0A0U5D5D4). The alignment is shown using ESPript 3.

**Supplementary S1, figure 2:** HME protein alignment for BLASTp

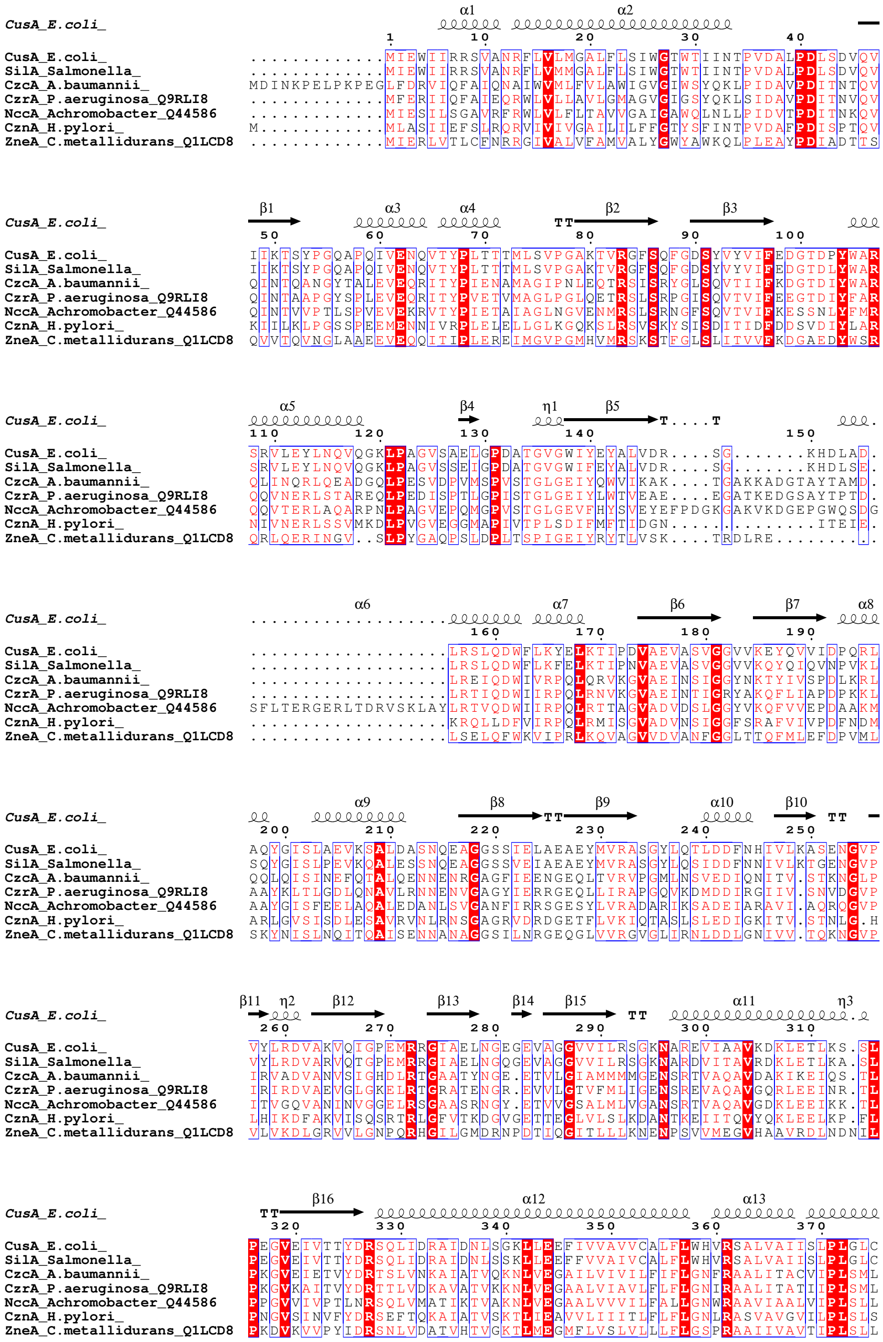
CusA_E.coli_
SilA_Salmonella_
CzcA_A.baumannii_
CzrA_P.aeruginosa_Q9RLI8
NccA_Achromobacter_Q44586
CznA_H.pylori_
ZneA_C.metallidurans_Q1LCD8

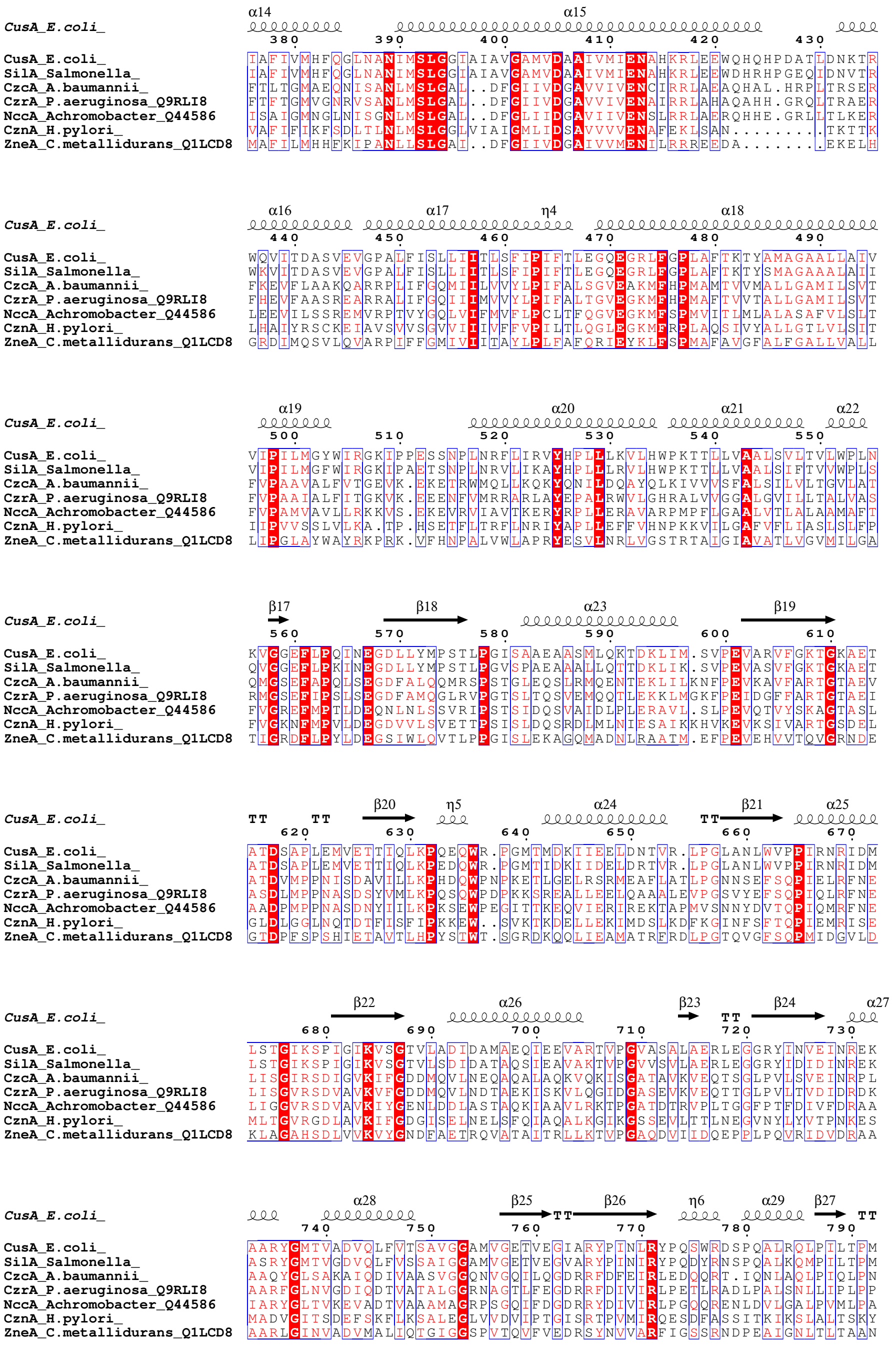
CusA_E.coli_ α14 α15
380 390 400 410 420 430
CusA_E.coli_ IAFIVMHFQGLNANIMSLGGIAIAVGAMVDAAIVMIENAHKRLEEWQHQHPDATLDNKTR
SilA_Salmonella_ IAFIVMHFQGLNANIMSLGGIAIAVGAMVDAAIVMIENAHKRLEEWDHRHPGEQIDNVTR
CzcA_A.baumannii_ FTLTGMAEQNISANLMSLGAL..DFGIIVDGAVVIVENCIRRLAEAQHAL.HRPLTRSER
CzrA_P.aeruginosa_Q9RLI8 FTFTGMVGNRVSANLMSLGAL..DFGIIVDGAVVIVENAIRRLAHAQAHH.GRQLTRAER
NccA_Achromobacter_Q44586 ISAIGMNGLNISGNLMSLGAL..DFGLIIDGAVIIVENSLRRLAERQHHE.GRLLTLKER
CznA_H.pylori_ VAFIFIKFSDLTLNLMSLGGLVIAIGMLIDSAVVVVENAFEKLSAN.........TKTTK
ZneA_C.metallidurans_Q1LCD8 MAFILMHHFKIPANLLSLGAI..DFGIIVDGAIVVMENILRRREEDA.........EKELH
CusA_E.coli_ α16 α17 η4 α18
440 450 460 470 480 490
CusA_E.coli_ WQVITDASVEVGPALFISLLIITLSFIPIFTLEGQEGRLFGPLAFTKTYAMAGAALLAIV
SilA_Salmonella_ WKVITDASVEVGPALFISLLIITLSFIPIFTLEGQEGRLFGPLAFTKTYSMAGAAALAII
CzcA_A.baumannii_ FKEVFLAAKQARRPLIFGQMIILVVYLPIFALSGVEAKMFHPMAMTVVMALLGAMILSVT
CzrA_P.aeruginosa_Q9RLI8 FHEVFAASREARRALIFGQIIIMVVYLPIFALTGVEGKMFHPMAFTVVTALLGAMILSVT
NccA_Achromobacter_Q44586 LEEVILSSREMVRPTVYGQLVIFMVFLPCLTFQGVEGKMFSPMVITLMLALASAFVLSLT
CznA_H.pylori_ LHAIYRSCKEIAVSVVSGVVIIIVFFVPILTLQGLEGKMFRPLAQSIVYALLGTLVLSIT
ZneA_C.metallidurans_Q1LCD8 GRDIMQSVLQVARPIFFGMIVIITAYLPLFAFQRIEYKLFSPMAFAVGFALFGALLVALL
CusA_E.coli_ α19 α20 α21 α22
500 510 520 530 540 550
CusA_E.coli_ VIPILMGYWIRGKIPPESSNPLNRFLIRVYHPLLLKVLHWPKTTLLVAALSVLTVLWPLN
SilA_Salmonella_ VIPILMGFWIRGKIPAETSNPLNRVLIKAYHPLLLRVLHWPKTTLLVAALSIFTVVWPLS
CzcA_A.baumannii_ FVPAAVALFVTGEVK.EKETRWMQLLKQKYQNILDQAYQLKIVVVSFALSILVLTGVLAT
CzrA_P.aeruginosa_Q9RLI8 FVPAAIALFITGKVK.EEENFVMRRARLAYEPALRWVLGHRALVVGGALGVILLTALVAS
NccA_Achromobacter_Q44586 FVPAMVAVLLRKKVS.EKEVRVIAVTKERYRPLLERAVARPMPFLGAALVTLALAAMAFT
CznA_H.pylori_ IIPVVSSLVLKA.TP.HSETFLTRFLNRIYAPLLEFFVHNPKKVILGAFVFLIASLSLFP
ZneA_C.metallidurans_Q1LCD8 LIPGLAYWAYRKPRK.VFHNPALVWLAPRYESVLNRLVGSTRTAIGIAVATLVGVMILGA
CusA_E.coli_ β17 β18 α23 β19
560 570 580 590 600 610
CusA_E.coli_ KVGGEFLPQINEGDLLYMPSTLPGISAAEAASMLQKTDKLIM.SVPEVARVFGKTGKAET
SilA_Salmonella_ QVGGEFLPKINEGDLLYMPSTLPGVSPAEAAALLQTTDKLIK.SVPEVASVFGKTGKAET
CzcA_A.baumannii_ QMGSEFAPQLSEGDFALQQMRSPSTGLEQSLRMQENTEKLILKNFPEVKAVFARTGTAEV
CzrA_P.aeruginosa_Q9RLI8 RMGSEFIPSLSEGDFAMQGLRVPGTSLTQSVEMQQTLEKKLMGKFPEIDGFFARTGTAEI
NccA_Achromobacter_Q44586 FVGREFMPTLDEQNLNLSSVRIPSTSIDQSVAIDLPLERAVL.SLPEVQTVYSKAGTASL
CznA_H.pylori_ FVGKNFMPVLDEGDVVLSVETTPSISLDQSRDLMLNIESAIKKHVKEVKSIVARTGSDEL
ZneA_C.metallidurans_Q1LCD8 TIGRDFLPYLDEGSIWLQVTLPPGISLEKAGQMADNLRAATM.EFPEVEHVVTQVGRNDE
CusA_E.coli_ TT TT β20 η5 α24 TT β21 α25
620 630 640 650 660 670
CusA_E.coli_ ATDSAPLEMVETTIQLKPQEQWR.PGMTMDKIIEELDNTVR.LPGLANLWVPPIRNRIDM
SilA_Salmonella_ ATDSAPLEMVETTIQLKPEDQWR.PGMTIDKIIDELDRTVR.LPGLANLWVPPIRNRIDM
CzcA_A.baumannii_ ATDVMPPNISDAVILLKPHDQWPNPKETLGELRSRMEAFLATLPGNNSEFSQPIELRFNE
CzrA_P.aeruginosa_Q9RLI8 ASDLMPPNASDSYVMLKPQSQWPDPKKSREALLEELQAAALEVPGSVYEFSQPIQLRFNE
NccA_Achromobacter_Q44586 AADPMPPNASDNYIILKPKSEWPEGITTKEQVIERIREKTAPMVSNNYDVTQPIQMRFNE
CznA_H.pylori_ GLDLGGLNQTDTFISFIPKKEW..SVKTKDELLEKIMDSLKDFKGINFSFTQPIEMRISE
ZneA_C.metallidurans_Q1LCD8 GTDPFSPSHIETAVTLHPYSTWT.SGRDKQQLIEAMATRFRDLPGTQVGFSQPMIDGVLD
CusA_E.coli_ β22 α26 β23 TT β24 α27
680 690 700 710 720 730
CusA_E.coli_ LSTGIKSPIGIKVSGTVLADIDAMAEQIEEVARTVPGVASALAERLEGGRYINVEINREK
SilA_Salmonella_ LSTGIKSPIGIKVSGTVLSDIDATAQSIEAVAKTVPGVVSVLAERLEGGRYIDIDINREK
CzcA_A.baumannii_ LISGIRSDIGVKIFGDDMQVLNEQAQALAQKVQKISGATAVKVEQTSGLPVLSVEINRPL
CzrA_P.aeruginosa_Q9RLI8 LISGVRSDVAVKVFGDDMQVLNDTAEKISKVLQGIDGASEVKVEQTTGLPVLTVDIDRDK
NccA_Achromobacter_Q44586 LIGGVRSDVAVKIYGENLDDLASTAQKIAAVLRKTPGATDTRVPLTGGFPTFDIVFDRAA
CznA_H.pylori_ MLTGVRGDLAVKIFGDGISELNELSFQIAQALKGIKGSSEVLTTLNEGVNYLYVTPNKES
ZneA_C.metallidurans_Q1LCD8 KLAGAHSDLVVKVYGNDFAETRQVATAITRLLKTVPGAQDVIIDQEPPLPQVRIDVDRAA
CusA_E.coli_ α28 β25 TT β26 η6 α29 β27 TT
740 750 760 770 780 790
CusA_E.coli_ AARYGMTVADVQLFVTSAVGGAMVGETVEGIARYPINLRYPQSWRDSPQALRQLPILTPM
SilA_Salmonella_ ASRYGMTVGDVQLFVSSAIGGAMVGETVEGVARYPINIRYPQDYRNSPQALKQMPILTPM
CzcA_A.baumannii_ AAQYGLSAKAIQDIVAASVGGQNVGQILQGDRRFDFEIRLEDQQRT.IQNLAQLPIQLPN
CzrA_P.aeruginosa_Q9RLI8 AARFGLNVGDIQDTVATALGGRNAGTLFEGDRRFDIVIRLPETLRADLPALSNLLIPLPP
NccA_Achromobacter_Q44586 IARYGLTVKEVADTVAAAMAGRPSGQIFDGDRRYDIVIRLPGQQRENLDVLGALPVMLPA
CznA_H.pylori_ MADVGITSDEFSKFLKSALEGLVVDVIPTGISRTPVMIRQESDFASSITKIKSLALTSKY
ZneA_C.metallidurans_Q1LCD8 AARLGINVADVMALIQTGIGGSPVTQVFVEDRSYNVVARFIGSSRNDPEAIGNLTLTAAN

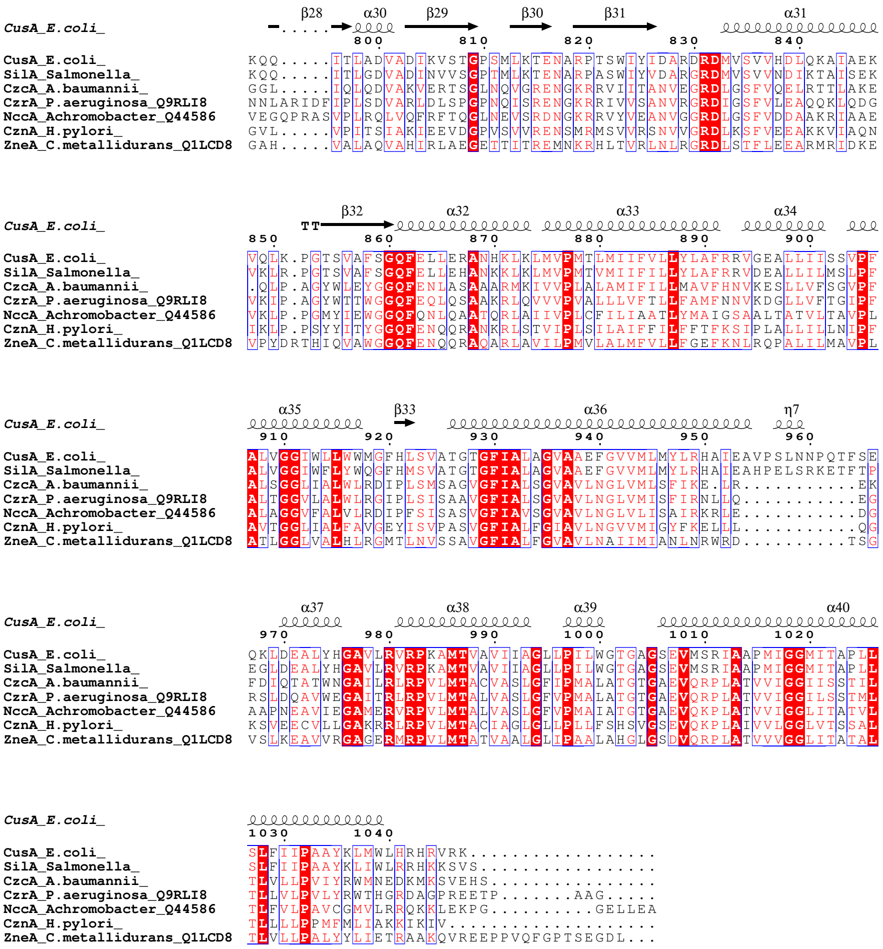

RND HME proteins from different Gram-negative bacteria were aligned against *E. coli* CusA reference sequence (P38054). The sequencws aligned were *Salmonella* SilA (Q9ZHC9), *A. baumannii* CzcA (A0A204EX98), *P. aeruginosa* CzrA (Q9RLI8), *Achromobacter* NccA (Q44586), *H. pylori* CznA (O25622) and *A. metallidurans* ZneA (Q1LCD8). The alignment is shown using ESPript 3.

**Supplementary S1, figure 3:** Phylogenetic tree of HME pumps with ArpB

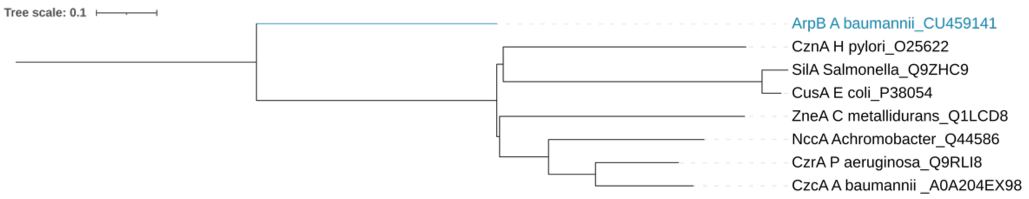


Phylogeny was created using a MAFFT (v.7) alignment of amino acid sequences of characterised heavy metal pumps and ArpB. The tree was constructed using a neighbour joining by phylo.io in MAFFT with a JTT substitution model and 100 bootstraps. The tree was coloured and midpoint rooted in iTol.

**Supplementary S1, figure 4:** Phylogenetic tree of HAE pumps with ArpB

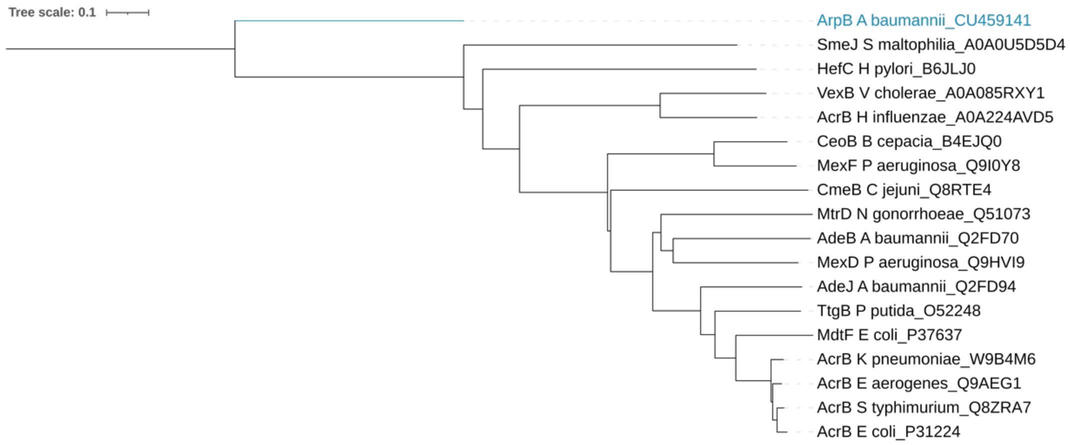


The phylogeny was created using a MAFFT (v.7) alignment of amino acid sequences of characterised hydrophilic and amphiphilic pumps and ArpB. The tree was constructed using a neighbour joining by phylo.io in MAFFT with a JTT substitution model and 100 bootstraps. The tree was coloured and midpoint rooted in iTol.
