## Supplemental 2 - tables for "RND pumps across the *Acinetobacter* genus; AdeIJK is the ancestral efflux system"

**Supplementary S2**

**Supplementary S2, table 1:** List of *Acinetobacter* genus sequences searched with accession codes

| Species | Accession code |
|---|---|
| *A. albensis* | GCA_015209685.1 |
| | GCA_900095025.1 |
| *A. apis* | GCA_900197575.1 |
| *A. baumannii* | CU459141.1 |
| | GCA_001433045.1 |
| | GCA_900444815.1 |
| | GCA_900493605.1 |
| *A. baylyi* | GCA_000046845.1 |
| | GCA_010577805.1 |
| | GCA_010577925.1 |
| | GCA_010577955.1 |
| *A. beijerinckii* | GCA_000368985.1 |
| | GCA_000369005.1 |
| | GCA_000931715.1 |
| *A. bereziniae* | GCA_001055215.1 |
| | GCA_001500155.3 |
| | GCA_006334885.1 |
| | GCA_016502565.1 |
| *A. bohemicus* | GCA_000367925.1 |
| | GCA_008802275.1 |
| | GCA_900116265.1 |
| | GCA_905183085.1 |
| *A. boisseri* | GCA_900096955.1 |
| *A. bouvettii* | GCA_000368865.1 |
| | GCA_001485025.1 |
| | GCA_004209145.1 |
| | GCA_902753875.1 |
| *A. brisouii* | GCA_000368645.1 |
| | GCA_000488275.1 |
| | GCA_000931655.1 |
| | GCA_000964015.1 |
| *A. calcoaceticus* | CP020000.1 |
| | GCA_000818215.1 |
| | GCA_000931735.1 |
| | GCA_001510805.1 |
| *A. celticus* | GCA_001707755.1 |
| *A. chengduensis* | GCA_003664645.1 |
| *A. chinesis* | GCA_002165375.2 |
| *A. colistiniresistens* | GCA_000413935.1 |

| | |
|---|---|
| | GCA_003227755.1 |
| | GCA_007713425.1 |
| | GCA_900406805.1 |
| *A. courvalinii* | GCA_008802255.1 |
| | GCA_014635545.1 |
| | GCA_016502305.1 |
| | GCA_016508605.1 |
| *A. cumulans* | GCA_003024525.3 |
| | GCA_003664635.1 |
| | GCA_003664655.1 |
| | GCA_012371365.1 |
| *A. defluvii* | GCA_001704615.3 |
| | GCA_013072655.1 |
| *A. dispersus* | GCA_001753605.1 |
| | GCA_009884975.1 |
| *A. equi* | GCA_001307195.1 |
| *A. gandensis* | GCA_001678755.1 |
| | GCA_008802205.1 |
| *A. gerneri* | GCA_000368565.1 |
| | GCA_000430245.1 |
| | GCA_000747725.1 |
| *A. guerrae* | GCA_003611455.1 |
| | GCA_009014115.1 |
| | GCA_009372255.1 |
| *A. guillouiae* | GCA_000829655.1 |
| | GCA_002370525.2 |
| | GCA_008996235.1 |
| | GCA_009011835.1 |
| *A. gyllenbergii* | GCA_000413855.1 |
| | GCA_000414075.1 |
| | GCA_000931695.1 |
| | GCA_001682515.1 |
| *A. haemolyticus* | CP018871.1 |
| | GCA_009899805.1 |
| | GCA_009899945.1 |
| | GCA_009899955.1 |
| *A. halotolerans* | GCA_004208515.1 |
| *A. harbinesis* | GCA_000816495.1 |
| *A. indicus* | GCA_002938515.1 |
| | GCA_002938785.1 |
| | GCA_002938915.1 |
| | GCA_013116605.1 |
| *A. johnsonii* | CP010350.1 |

| | |
|---|---|
| | GCA_007989645.1 |
| | GCA_009789205.1 |
| | GCA_012271875.1 |
| *A. junii* | CP019041.1 |
| | GCA_004123275.1 |
| | GCA_012271855.1 |
| | GCA_901873405.1 |
| *A. kookii* | GCA_900096895.1 |
| *A. lactucae* | CP020015.1 |
| | GCA_000399705.1 |
| | GCA_000516615.2 |
| | GCA_001415555.1 |
| *A. lanii* | GCA_011191955.1 |
| | GCA_011578285.1 |
| *A. larvae* | GCA_001704115.1 |
| *A. lwoffii* | GCA_000836095.1 |
| | GCA_006538585.1 |
| | GCA_014769185.1 |
| | GCA_900444925.1 |
| *A. modestus* | GCA_014636095.1 |
| *A. nectaris* | GCA_000488215.1 |
| *A. nosocomialis* | CP040105.1 |
| | GCA_000694975.1 |
| | GCA_001056825.1 |
| | GCA_002144915.1 |
| *A. parvus* | GCA_000248155.2 |
| | GCA_000368025.1 |
| | GCA_000962795.1 |
| | GCA_001485085.1 |
| *A. piscicola* | GCA_002233755.1 |
| | GCA_004152775.1 |
| *A. pittii* | CP002177.1 |
| | GCA_900110525.1 |
| | GCA_900495015.1 |
| | GCA_900496515.1 |
| *A. populi* | GCA_002174125.1 |
| *A. portensis* | GCA_009372215.1 |
| | GCA_010646905.1 |
| *A. pragensis* | GCA_001605895.1 |
| *A. proteolyticus* | GCA_000367945.1 |
| | GCA_001753605.1 |
| | GCA_002835245.1 |
| | GCA_902505965.1 |

| | |
|---|---|
| *A. pseudolwoffii* | GCA_002803605.1 |
| | GCA_900088145.1 |
| *A. puyangensis* | GCA_900096995.1 |
| *A. qingfengensis* | GCA_001753595.1 |
| | GCA_008693185.1 |
| *A. radioresistens* | CP030031.1 |
| | GCA_003006715.1 |
| | GCA_003594775.1 |
| | GCA_006370545.1 |
| *A. rudis* | GCA_000413895.1 |
| | GCA_000829675.1 |
| *A. schindleri* | GCA_001485065.1 |
| | GCA_003069395.1 |
| | GCA_010918895.1 |
| | GCA_014204595.1 |
| *A. seifertii* | GCA_001054375.1 |
| | GCA_002795375.1 |
| | GCA_004378605.1 |
| | GCA_016064815.1 |
| *A. shaoyimingii* | GCA_011174715.1 |
| | GCA_011578045.1 |
| *A. sichuanensis* | GCA_003024515.2 |
| *A. soli* | CP016896.1 |
| | GCA_000760595.1 |
| | GCA_002204165.1 |
| | GCA_018863215.1 |
| *A. tandoii* | GCA_000400735.1 |
| | GCA_000760555.1 |
| | GCA_002795165.1 |
| | GCA_008867985.1 |
| *A. tjernbergiae* | GCA_000488175.1 |
| | GCA_000759995.1 |
| *A. towneri* | CP071766.1 |
| | GCA_000760575.1 |
| | GCA_004786115.1 |
| | GCA_018501125.1 |
| *A. ursingii* | GCA_000949815.1 |
| | GCA_000949835.1 |
| | GCA_001056675.1 |
| | GCA_901873655.1 |
| *A. variabilis* | GCA_000369625.1 |
| | GCA_003938405.1 |
| | GCA_009822135.1 |

| | |
|---|---|
| | GCA_016607565.1 |
| *A. venetianus* | GCA_000368585.1 |
| | GCA_001484985.1 |
| | GCA_001575095.1 |
| | GCA_001577485.1 |
| *A. vivanii* | GCA_016502725.1 |
| | GCA_014635885.1 |
| *A. wanghuae* | GCA_009557235.1 |
| | GCA_009601085.1 |
| *A. wuhouensis* | GCA_001696605.3 |
| | GCA_002165345.2 |
| | GCA_004209115.1 |
| | GCA_004209325.1 |

**Supplementary S2, table 2:** Strains used in this study

| Strain | Reference |
|---|---|
| *A. baumannii* AYE | (Fournier et al., 2006) |
| *A. baylyi* ADP1 | (Vaneechoutte et al., 2006) |
| *A. baumannii* ATCC 17978 | Gift from Laura Piddock |
| *A. baumannii* ATCC 17978 *ΔadeAB ΔadeFGH ΔadeIJK* | Gift from Ayush Kumar |
| *A. baumannii* ATCC 17978 *ΔadeAB ΔadeFGH ΔadeIJK* + empty pVRL2 | This study |
| *A. baumannii* ATCC 17978 *ΔadeAB ΔadeFGH ΔadeIJK* + pVLR2 with *A. baumannii* AYE *adeIJK* | This study |
| *A. baumannii* ATCC 17978 *ΔadeAB ΔadeFGH ΔadeIJK* + pVRL2 with *A. lwoffii* 5867 *adeIJK* | This study |
| *A. baumannii* ATCC 17978 *ΔadeAB ΔadeFGH ΔadeIJK* + pVRL2 with *A. baylyi* ADP1 *adeXYZ* | This study |

**Supplementary S2, table 3:** Primers used in this study

| Primer name | Sequence (5' - 3') |
|---|---|
| Forward cloning primer for *A. baumannii* AYE *adeIJK* | CTAGTACCCGGGATGATGTCGGCTAAGCTTTG |
| Reverse cloning primer for *A. baumannii* AYE *adeIJK* | TACTGTGCGGCCGCTTATTGCTTTTTAAGTTCAG |
| Forward cloning primer for *A. baylyi* ADP1 *adeXYZ* | TGATATCGAATTCCTGCAGCCCGGGATGACGTCGGCTAAGCTTTGG |
| Reverse cloning primer for *A. baylyi* ADP1 *adeXYZ* | AATTGGAGCTCCACCGCGGTGGCGGCCGCTTATTTCTTTTTCAGATCTGCACTTGAAGGCTGATG |

**Supplementary S2, table 4:** HAE proteins found in 100 *A. baumannii* sequences

| Number of HAE proteins | Percentage number of sequences (%) |
|---|---|
| 4 | 3 |
| 5 | 95 |
| 6 | 1 |
| 7 | 1 |

HAE proteins were highlighted in a BLASTp search for conserved HAE RND residues

**Supplementary S2, table 5:** HME proteins found in 100 *A. baumannii* sequences

| Number of HME proteins | Percentage number of sequences (%) |
|---|---|
| 1 | 67 |
| 2 | 27 |
| 3 | 5 |
| 4 | 1 |

HME proteins were highlighted in a BLASTp search for conserved HME RND residues
