## Supplemental 3 - text for "RND pumps across the *Acinetobacter* genus; AdeIJK is the ancestral efflux system"

### Supplementary S3

#### Supplementary S3, text 1: HAE RND conserved residue sequence for BLASTp

```
>HAE_RND_CONSERVED_RESIDUES
--------
XXXXFIXRPXXAXXXXXXIXXXGXXXXXXLPXXXYPXXXXPXXXXXXXYPGAXAXXXXXXVXXXXEXXXXGXXXXXYXXSXXXXXGXXXXXXX
FXXXTXXXXAXXXVXNXXXXXXXXLPXXXXXXXXXXXXXXXXXXXXXXXXXXXX--XXXXXXXXXYXXXXXXXXXXXXXGVGXXXXXGX-
XXAXRXWXXPXXXXXXXLXXXDVXXAXXXXNXQXXXGXXXXXXXXXXXXXXXXXXXXXXLXXXXXXXXXXXXXXXXGXXXXLXDXAXXXXGXX
XXXXXXXXNGXXXXXXXIXXXXGANXXXXXXXXXXXXXXXXXXXPXXXXXXXXYDXTXXXXXSIXXVXXTLXEAXXLVXXVXXXFLXXXRXXX
IPXXXXPXXXXGTFXXXXXXGFSXNXLTXXXXXLAIGXXVDDAIVVXENXXRXXXX-
XGXXXXXAXXXXXXXXXXXXXXXXXXLXAVFXPXAXXXGXXGXXXXXFXXTXXXXXXXSXXXXLXXXPXXXAXXLKXX---XXXXXX-
XXXXX------------
XXXXXXXFXXXXXXYXXXXXXXXXXXXXXXXXXXXXXXXXXXXXXXXPXXXXPXEDXGXXXXXXXXPXXXXXXXXXXXXXXXXXXXX--
XXXXVXXXXXXXGXXXXX--XXXNXXXXFXXLKXXXXRXXX-XXXXXXXXXXXX-XXXXXXXXX-XXXXXXXPXXX---
XLGXXXGXXXXXXXXXXXXXXXXXXXXXXXXXXXXXXXXX----XXXXXXXXXXXXXXXXXXDXXXAXXXGVXXXXIXXTXXXXX-
GXXXXXXFXXXGRXXXVXXXXXXXXRXXXXXXXXXXVXXX-------------------------------------
XGXXXPXXXXXXXXXXXXXXXXXRXNXXXXXXXXXXX--AXGXSXGAXXXXXXXXXX--LPX----
GXXXXXXGXXXXXXXXXXXXXXXXXXXXXXXFLXLAAXYESXXXPXXXXXXVPXXXXGAXXXXXX-----------
XXXXXXXXXXXGLXXXXGLXXKNAILIXEFAXXXX-
XXGXXXXXAXXXAXXXRXRPIXMTXXAXXXGXXPLXXXXGAGXXXXXXXGXXXXXGXXXXTXXXXXXXPXXXXXXXXXXXXXXXXXXXX----
---------XXXXX
```

**Supplementary S3, text 2: HME RND conserved residue sequence for BLASTp**

```
> HME_RND_CONSERVED_RESIDUES
MXXXXXXXXXXXRXXVXXXXXXXXXXGXXXXXXXPXDAXPDXXXXQVXIXTXXXGXXXXXVEXXXTXPXXXXXXXXXGXXXXRXXSXXGXSXX
XXXFXXXXXXYXXRXXXXEXLXXXXXXLPXXXXXXXXPXXTXXGXIXXXXXXXX----XXXXXX---XXXXXX--------------------
LRXXQDWXXXXXLXXXXXVAXXXXXGGXXXXXXXXXDXXXXXXXXXSXXXXXXAXXXXNXXXGXXXXXXXXXXXXVRXXXXXXXXXDXXXIXX
-XXXXGXPXXXXDXAXVXXGXXXRXGXXXXXGXXEXXXGXXXXXXXXNXXXVXXXVXXXXXXXXX-
XLPXGVXXXXXXDRXXLXXXXXXXXXXXLXEXXXXVXXXXXXFLXXXRAXXXXXXXPLXXXXXXXXMXXXXXXXNXMSLGXX--
XXGXXXDXAXVXXENXXXRLXXXXXXX-
XXXXXXXXXXXXXXXXXXXXXXXXXXXXXXIXXXXXPXXXXXGXEXXXFXPAXXXXXAXXGAXXLXXXXXPXXXXXXXXXXXX-
XXXXXXXXXXXXXYXXXLXXXXXXXXXXXXXAXXXXXXXXXXXXXXGXXFXPXXXEGXXXXXXXXXPXXSXXXXXXXXXXXXXXXX-
XXPEVXXXXXXXGXXXXXXDXXXXXXXXXXXXLXPXXXWX-
XXXXXXXXXXXXXXXXXXXGXXXXXXXPIXXRXXXXXXGXXSXXXXKXXGXXXXXXXXXAXXIXXXXXXXXGXXXXXXXXXXGXXXXXXXXXX
RXXXAXXGXXXXXXXXXXXXXXXGXXXXXXXXGXXRXXXXXRXXXXXRXXXXXXXXLXXXXXXXXX-----
XXLXXXAXXXXXXGXXXXXXENXXXXXXXXXXXXGRDXXSXVXXXXXXXXXXXXXX-
XXXXXXXXGQFEXXXXAXXXLXXXXPXXXXXXFXXLXXXFXXXXXXXLXXXXXPXAXXGGXXXLXXXXXXXSXXXXXGFIALXGVAXXXGXVM
XXXXXXXXX-----------
XXXXXXXAXXXGAXXRXRPXXMTXXVXXXGXXPXXXXXGXGXEVXXXXAXXXXGGXXXXXXLXLXXXPXXXXXXXXXXXXXXXXX--------
--------
```

**Supplementary S3, text 3:** 100 *A. baumannii* sequence accession identifiers

GCA_000015425.1_ASM1542v1
GCA_000018445.1_ASM1844v1
GCA_000021245.2_ASM2124v2
GCA_000069205.1_ASM6920v1
GCA_000069245.1_ASM6924v1
GCA_000163355.2_ASM16335v2
GCA_000163375.2_ASM16337v2
GCA_000163395.2_ASM16339v2
GCA_000173395.1_ASM17339v1
GCA_000184475.2_ASM18447v2
GCA_000184495.2_ASM18449v2
GCA_000184515.2_ASM18451v2
GCA_000186645.2_ASM18664v2
GCA_000186665.4_ASM18666v4
GCA_000187205.4_ASM18720v4
GCA_000188215.1_ASM18821v1
GCA_000189655.2_ASM18965v2
GCA_000189675.2_ASM18967v2
GCA_000189695.2_ASM18969v2
GCA_000189735.2_ASM18973v2
GCA_000214965.2_gacin09v1.0
GCA_000214985.2_gacin10v1.0
GCA_000215005.2_AcbauOIFC032v1.0
GCA_000222225.2_ASM22222v2
GCA_000222245.2_ASM22224v2
GCA_000222265.2_ASM22226v2
GCA_000222285.2_ASM22228v2
GCA_000226275.2_ASM22627v2
GCA_000241665.2_ASM24166v2
GCA_000241685.2_ASM24168v2
GCA_000241705.2_ASM24170v2
GCA_000241725.2_ASM24172v2
GCA_000248275.2_ASM24827v2
GCA_000278605.1_gacin08v1.0
GCA_000278625.1_AcbauOIFC137v1.0
GCA_000278645.1_AcbauOIFC109v1.0
GCA_000278665.1_AcbauOIFC143v1.0
GCA_000278685.1_AcbauOIFC189v1.0
GCA_000282795.3_ASM28279v3
GCA_000286535.1_gacin11v1.0
GCA_000286615.1_gacin07v1.0
GCA_000292545.1_ASM29254v1
GCA_000297515.1_gacin24v1.0
GCA_000297535.1_gacin24v1.0
GCA_000297575.1_gacin25v1.0
GCA_000297595.1_gacin26v1.0
GCA_000297935.1_Acin_baum_Ab33333_V1
GCA_000299655.1_ASM29965v1
GCA_000299675.1_ASM29967v1
GCA_000301175.1_ASM30117v1
GCA_000301195.1_ASM30119v1
GCA_000301215.1_ASM30121v1
GCA_000301235.1_ASM30123v1
GCA_000301255.1_ASM30125v1
GCA_000301275.1_ASM30127v1
GCA_000301295.1_ASM30129v1
GCA_000301315.1_ASM30131v1
GCA_000301335.1_ASM30133v1
GCA_000301355.1_ASM30135v1
GCA_000301375.1_ASM30137v1
GCA_000301395.1_ASM30139v1
GCA_000301415.1_ASM30141v1
GCA_000301435.1_ASM30143v1
GCA_000301455.1_ASM30145v1
GCA_000301475.1_ASM30147v1
GCA_000301495.1_ASM30149v1
GCA_000301515.1_ASM30151v1
GCA_000301535.1_ASM30153v1
GCA_000301555.1_ASM30155v1
GCA_000301575.1_ASM30157v1
GCA_000301595.1_ASM30159v1
GCA_000301615.1_ASM30161v1
GCA_000301835.1_ASM30183v1
GCA_000301875.1_ASM30187v1
GCA_000301895.1_ASM30189v1
GCA_000301935.1_ASM30193v1
GCA_000301955.1_ASM30195v1
GCA_000301975.1_ASM30197v1
GCA_000301995.1_ASM30199v1
GCA_000302015.1_ASM30201v1
GCA_000302035.1_ASM30203v1
GCA_000302055.1_ASM30205v1
GCA_000302075.1_ASM30207v1
GCA_000302135.1_ASM30213v1
GCA_000302155.1_ASM30215v1
GCA_000302215.1_ASM30221v1
GCA_000302235.1_ASM30223v1
GCA_000302255.1_ASM30225v1
GCA_000302575.1_ASM30257v1
GCA_000304655.1_gacin18v1.0
GCA_000304675.1_GACIN20v1.0
GCA_000304695.1_gacin17v1.0
GCA_000305215.1_gacin13v1.0
GCA_000305235.1_gacin14v1.0
GCA_000305255.1_gacin15v1.0
GCA_000305275.1_gacin22v1.0
GCA_000305295.1_gacin16v1.0
GCA_000305315.1_gacin21v1.0
GCA_000307895.1_gacin19v1.0
GCA_000307975.2_ASM30797v2

**Supplementary S3, text 4:** ABRicate custom database sequences for finding *Acinetobacter* PAP, RND and OMF genes “AciRND”

```
>AciRND~~~adeA_baumannii~~~CU459141.1~~~AdeA periplasmic adaptor protein for AdeABC RND
efflux pump in Acinetobacter baumannii
TTATGCATTTGAAGCTGCTTTCTGTTCTGCTTGAGGTTTACCTGATTGAGCCGGTTTCGCAGCATTTGGT
GCTGCACCTTGAGAGTTTGCTGGTTGAGCTTGATAAGGTTTTGCTGATACTTCTTGCCCTTCTTTAACTT
TGGCAACACCATCAACAATGACTTTATCGCCGGCTTTTAAGCCGTTAGTCACAATCCAGTTTTGTCCTTG
AACACCAGAGGTTTCAACAGGACGGCTCTCAACAACCCCTTTAGCATTAACAAGCATCGCTACAGCTTGT
CCTGTAGGTAAACGAGTAATGGCAGCTTGAGGAATCAGGTAAGCATTTGGAACAACGCCCTGAACAATTT
GCGCAGTGGTATACATACCCGGAAGCAATAAATGATTCGGGTTAGAGAATACGGCACGTAATGTAATTGT
TCCTGTATCTTGGTTTACAGAAGCGTCAGAGAAAGCAAGTTGCCCTTCGATTGGATAGGTAGAACCATCT
TCAAGCTTTAATTTTACTTTCGTGTTGTTACTGTTATTTAAACTGCCTTTGCTTAGTTGTTGACGTAAAC
GCAATAACTCAGCACTAGACTGATTAATATCAACATAGATAGGATCTAATTGTTGAATCGTTACCAACGG
GTCAGTCTGGTTTGCAGTAACCAAAGCACCAGCCGTTACTGAAGAACGACCAGATTGGCCAGAAATAGGA
GAGCGAATTGTAGAATAACCAAGATCTACATTTGCATTTGTTACTTGAGCCTTAGCTGCTGCAACTTGTG
CTTCTGCAACATTGACTTGACCAAGTAAGTCATCATATTCCTGTTTAGACACAGCATTACTAGAAACAAG
TTGTTTATAACGATTTAACTTGGTACGTAGTGAAGCTAGATTTGCCTGTTGTTGTAGGAGTGATGCTTTT
GCATTTTCTAACGTTGCACGGTTCGTTCTAGAGTCGAGCTCATAAAGCGCCTGACCTTCACGAACATAGC
TTCCTTCAGCAAATAAACGTTTTAAAATCACGCCACTTGTTTGAGGACGAACTTCAGAAATTTGATATGC
TGAAGTACGGCCTGAAAGCTCAACGCTTTGTTCAACACTTTGTGGTTGAGCAACAATAACACCTACTTCT
GCAGGCGGCATTTTCTGAGCAGCAGCAGCTTGCTGTTTCTCATCGGAGCCTTTGCTACAACCAACAAGCG
CGATACTTGTTGCTAATGCGCAAGCAGTAAGGGCTGGTGCCCAAAGCTTAGCCGACATCAT
>AciRND~~~adeB_baumannii~~~CU459141.1~~~AdeB RND pump gene for AdeABC efflux pump in
Acinetobacter baumannii
ATGATGTCACAATTTTTTATTCGTCGTCCCGTTTTTGCTTGGGTTATCGCGATCTTCATTATTATATTTG
GATTGCTGAGTATTCCTAAACTGCCAATTGCACGTTTTCCAAGTGTAGCCCCGCCACAGGTGAATATTAG
TGCGACTTATCCTGGTGCTACAGCTAAAACCATTAACGATAGCGTTGTAACCTTAATTGAGCGCGAATTA
TCGGGTGTAAAAAATCTACTCTACTATAGTGCGACAACAGATACCTCCGGTACAGCAGAGATTACCGCTA
CGTTTAAACCGGGCACAGATGTGGAAATGGCTCAGGTCGACGTTCAAAATAAAATCAAGGCTGTAGAAGC
TCGCTTGCCGCAAGTCGTACGTCAGCAAGGCTTACAGGTTGAAGCTTCATCGTCCGGATTTTTAATGCTG
GTCGGGATTAACTCTCCAAATAATCAATATTCCGAAGTTGATTTGAGTGATTATTTGGTTCGAAATGTTG
TAGAAGAGCTAAAACGTGTCGAAGGTGTAGGGAAGGTTCAATCTTTCGGTGCAGAGAAAGCTATGCGTAT
TTGGGTCGACCCGAATAAGCTTGTTTCTTACGGTTTATCGATTAGTGATGTGAATAATGCCATTCGTGAA
AATAATGTCGAAATTGCACCCGGCCGACTTGGTGATTTACCAGCTGAAAAAGGCCAGCTCATTACTATTC
CATTGTCTGCTCAAGGGCAATTGTCTAGTCTTGAGCAATTTAAAAATATCAGCTTAAAAAGTAAAACTAA
CGGTAGCGTAATTAAATTATCTGATGTTGCCAATGTAGAAATAGGTTCACAAGCATATAACTTTGCCATT
TTGGAAAATGGTAAGCCTGCTACCGCGGCTGCAATTCAATTAAGCCCGGGAGCTAACGCCGTGAAAACTG
CCGAAGGTGTTCGAGCAAAAATTGAAGAATTGAAGCTAAATTTACCGGAAGGCATGGAATTTAGTATTCC
TTACGACACCGCGCCGTTTGTCAAAATTTCAATTGAAAAGGTAATTCATACATTACTTGAAGCCATGGTT
CTGGTTTTCATTGTGATGTATCTATTTTTACACAATGTCCGCTATACGCTTATTCCAGCAATTGTGGCGC
CTATTGCCTTACTCGGTACTTTTACCGTGATGTTGCTTGCCGGCTTTTCAATTAACGTACTCACCATGTT
CGGTATGGTGCTTGCCATCGGGATTATTGTCGACGATGCCATTGTTGTCGTTGAAAACGTAGAAAGGATT
ATGGCGACAGAAGGATTATCGCCTAAAGATGCAACCTCTAAAGCAATGAAAGAAATTACCAGCCCGATTA
TTGGTATTACGCTGGTATTGGCGGCAGTATTTTTACCTATGGCATTTGCGAGTGGTTCTGTAGGGGTAAT
CTATAAACAGTTTACCTTGACCATGTCGGTATCTATTTTATTTTCAGCGCTATTGGCACTCATTTTAACA
CCGGCACTTTGTGCCACGATTTTAAAGCCAATCGATGGGCATCATCAGAAGAAGGGGTTCTTTGCATGGT
TTGACCGTAGTTTCGATAAAGTCACTAAAAAGTATGAATTGATGCTGCTTAAAATCATCAAACATACAGT
TCCAATGATGGTGATCTTTTTAGTAATTACCGGTATTACCTTTGCCGGAATGAAATATTGGCCAACAGCA
TTTATGCCAGAGGAAGATCAAGGTTGGTTCATGACTTCGTTCCAGCTACCTTCAGATGCAACCGCAGAGC
GTACTCGGAATGTAGTCAATCAATTTGAAAATAATTTGAAAGACAATCCCGATGTAAAAAGTAATACCGC
CATTTTGGGATGGGGTTTTAGTGGTGCAGGACAAAATGTAGCTGTGGCTTTTACGACACTTAAAGACTTC
AAAGAGCGGACTAGCTCTGCATCCAAGATGACAAGCGACGTTAATTCTTCTATGGCGAACAGTACGGAAG
GCGAGACCATGGCCGTTTTACCACCCGCTATTGATGAGTTAGGTACTTTTTCAGGTTTCAGCCTACGTTT
ACAAGACCGTGCTAACTTAGGTATGCCTGCTTTATTGGCTGCTCAAGATGAACTTATGGCAATGGCAGCC
AAGAATAAAAAGTTCTATATGGTTTGGAATGAAGGGTTGCCACAAGGTGACAATATTTCTTTAAAAATTG
ACCGTGAAAAGCTTAGTGCACTTGGTGTTAAGTTTTCTGATGTTTCAGACATCATCTCTACATCAATGGG
TTCAATGTATATCAATGACTTCCCTAATCAAGGACGTATGCAACAAGTCATTGTACAAGTTGAGGCTAAA
TCACGTATGCAATTGAAAGATATCTTGAATCTGAAAGTCATGGGTTCAAGCGGTCAATTAGTCTCGTTAT
CAGAAGTTGTAACGCCACAATGGAATAAGGCACCACAACAATATAATCGTTATAACGGACGACCATCTTT
GAGTATTGCTGGTATTCCTAACTTCGATACGTCATCGGGTGAAGCAATGCGTGAAATGGAACAACTGATT
GCGAAATTACCGAAAGGTATTGGCTACGAGTGGACAGGTATTTCCTTACAGGAAAAGCAGTCTGAATCAC
AAATGGCCTTTTTACTTGGTTTATCAATGCTCGTTGTTTTCCTCGTATTGGCTGCACTCTATGAAAGCTG
GGCAATTCCACTTTCTGTGATGCTGGTTGTGCCACTCGGTATTTTTGGAGCAATCATTGCCATTATGTCT
AGAGGGTTAATGAATGATGTGTTCTTCAAAATCGGGCTAATTACCATTATTGGTCTATCGGCAAAGAATG
CAATTTTGATTGTTGAATTTGCGAAAATGCTAAAAGAGGAAGGCATGAGTTTGATTGAAGCCACTGTTGC
CGCAGCCAAACTTCGCTTACGACCAATCCTAATGACTTCTCTTGCATTTACGTGTGGTGTAATTCCTTTG
GTTATTGCAACAGGTGCAAGTTCAGAAACTCAACATGCTTTAGGCACAGGGGTTTTTGGTGGCATGATTT
CAGCAACCATTCTGGCTATTTTCTTTGTTCCCGTGTTTTTTATCTTCATTTTGGGTGCAGTAGAAAAGCT
ATTTTCCTCTAAGAAAAAAATCTCATCCTAA
>AciRND~~~adeC_baumannii~~~CU459141.1~~~AdeC outer membrane channel for AdeABC efflux pump in
Acinetobacter baumannii
TTGGAGAATACTATGTCTAAATCGGCAATCGTATCTCGTGGACTCATTCTTTCTACACTCTCAGTCACTT
TAGTTGCATGTGTCAATATGCAAGCGCCACAGCCTGCAATCACATCTCATATTCCTCAAAATTTTAGTCA
AAATCATTCTGGAAAAACGATTGCAGAAAAAAGTTATAAAGAATTTATTTCTGATCCGAAATTAGTACAG
```

GTCATTGAAATCAGTTTAAATAACAACCGTGATTTACGGACTGCTACGCTTAATATTGAACGTGTACAGC
AACAATACCAAATCACAAAAAATAGCCAGCTCCCAACCATTGGCGTAACGGGAAATGCAGTGCGGCAGGT
TAGCCCATCGATTAACCCCAATAACCCAGTTTCTACATTTCAGGTGGGTTTAGGAATGACTGCATATGAA
CTAGACTTCTGGGGCCGTGTTCAAAATTTAAAAGATGCTGCATTAAATAACTACCTTGCAACTCAAAGCG
CGAAAGAGGCTGTGCAAATTAGTTTAATCAGTAACATCACACAGGTTTGGTTAAATTATGCTTTTGCACA
AGCCAACTTAAACCTTGCCGAGCAAACCTTAAAAGCACAAGTCGATGCTTATAATCTAAATAAGAAGCGC
TTTGATGTTGGTATTGATAGTGAAGTTCCATTAAAACAAGCGCAAATTTCGGTAGAGACTGCTCGAAATG
ATGTTGCAACTTATAAAACTCAAATTCAACAGGCAAAAAATTTACTGGATTTATTAGCAGGTCATCCTGT
TCCGCAAAATTTACTTCCGAATCATGCTATTCAAAATATTACCTTTGAGAAAAACTTTGCAGCCGGTTTA
CCAAGTGATTTATTAAATCATCGTCCAGACCTTAAAGCTGCCGAATATGAGTTACGTGCTGCAGGAGCAA
ATATTGGTGCTGCTAAAGCACGGATGTTCCCAACCATAAGCCTGACAGGCTCGACGGGTTATGCATCATC
TGAACTGAAAGATTTATTTAAAACAGGCAATTTTGCATGGTCGATTGGACCTAATATCGATCTACCAATT
TTTGATTGGGGAACAAGAAAAACTAATATTAAAATTGCGGAAACTGACCAGAAAATTGCTTTAGCTAAAT
ATGAAAAAGCCATTCAATCAGCTTTTCGTGAAGTTAATGATGCGCTTGCTACACATGCGCATATTGGTGA
ACGATTAGACGCTCAGCGTCGTTTAGTCTCTGCGACTGCTGCAACCTATAAACTATCTATGGCGCGCTAC
AGAGCGGGTGTTGATAGTTATTTTACGGTTTTAGATGCGCAACGTTCCGCTTACGCAGCACAACAAGGAT
TACTTGCGCTTGAGCAAATGGAATTAAATAACCAGATTGAACTCTATAAAGTTTTAGGAGGAGGAATATC
AAAAGTCTAA
>AciRND~~~adeD_pittii~~~NC_016603.1~~~AdeD periplasmic adaptor protein of AdeDE efflux pump in Acinetobacter pittii
TCAACTCATTGGGGTAGGTTCAATCGTCATGCCGGCAGTTAAGCCTTGCAGCCCCTCAACAATGACTTGA
TCTCCCGCTTTTAGGCCAGAGGTCACTAACCAACGATTATCAATTGCATCGCCTAGAGTAATTTTTCGTT
GCTCGGCTTTATTATCTTTTCCAACAATCCAAACAAGTGGATTTCCTTCTGCGTCTCGGAACAGCGCACC
TTGAGGAATTAAGACGCTGTTAAGACGTGTTCCCGTTCCAAGTTCTGCCCGCACATACAGGCCGGGTAAT
AACGCATTATTAGGATTGGGAAATGCTGCACGTAAGGTGACACTTCCAGTCGCCTCATCTACAGCAACAT
CACTAAACTTAAAAGTCCCCTGTTCTGCATAGGCAGTGCCATTCTCAAGCTTCAAACGCACTGAAAGTTC
AGTCGGTTTTATTCCATTTTCAGTAAGCTGTTTACGCAACGCCATGTATTCATCACTCGACTGAGTCAAA
TCGACATACATGGGGTCAAGTTTTTGAATCGTTGCTAAAGGTTCTGTTTGAGCAGAGGTGACTAAAGCCC
CACGTGTAATAGATGAACGCCCTATTCGTCCTGAAATTGGCGCACTAACTTGTGTGTAACGTAAATTGGT
TCGTGCTGTTTTCAGTACAGCTTCATTTACGGCAACAGTCGCTTTCGCTTGCTCATATGCACTTTGAGCA
TTATCAAGCTCTTGTTGACTCACCCCATTTACTTTAATTAATTCTTTATAACGCTCGGCTTGAAGGCGAG
TCGAATTTACAGTGGCTTTTGCTAATGCAAGATTACCAGCGGCTTCATCCACACTATCCCGATAGAGTGA
TGAATCAATTTTATAAAGTGGCTGCCCTTTGTTTACTTGCGTGCCTTCCTTAAATAATTGATCTATTACT
ACTCCATTGACTTGTGGACGGACCTGAGAAATTTCAGAGGCCACCGTTCTACCAGCCAATTGATAGGTAG
ATTCAATCGTCTCTGGTTTTATAGTGATCGTACCCACCTGCTGAATATTTTTAGGGGCAGGCTGGTCAGC
ATCACAGCCATTCAAGAAAAGCAACAAAGGCAATAAACACAAAGTATTACGATGAAAAAAA
>AciRND~~~adeE_pittii~~~NC_016603.1~~~AdeE RND pump gene n of AdeDE efflux pump in Acinetobacter pittii
TCATTTTGCCTTTTTGGTGTTTAACTTATTGGTTAAACGCCGAACTAAAAGAAAAAATAAAGGCACAAAG
AAAATAGAAAGTAGAGTTCCCGAAATCATTCCACCAGTTACGGCGATACCAATTTCTTTTCGGCTAACAG
CTCCTGCTCCTGTAGATACAGCTAATGGAAGAACACCTGCAACAAAAGCCAGAGATGTCATAATAATTGG
CCGTAATCGCTGACCAGCACCTTCAATAATGGCATCCATTAATGCCTGACCAGCCTCTAACTTGGCTGCC
GCAAATTCAACAATCAAAATAGCATTTTTGGCAGAAAGTCCAATGGTGGTAAGCATAGCGACCTGAAAGT
AAATATCGTTAACAAAGCCTGCCAGACTTGCAGCCACTACAGCGCCAATAAGACCTAAAGGAATTACTAG
CATGACAGAGACAGGAATAGACCAACTTTCATAAAGTGCTGCCAAACATAAAAAGATAAAGATGATTGAG
GCTAAATATAACCAGATTGTTTGGCCACCCGCTAGTTTCTCTTGATAAGACAACCCACTCCACTGAAGAT
TAAAACCTTGTTGCTGGTCTACTAACTCCTGCATCTTATCCATAGCCCCACCAGAACTCTCACCAGCTGC
CGCAGAGCCTTGTATTTGAACGGCTGAAAGACCGTTAAAGCGTTGCAGCATTTGAGGGCCCATCTGCCAT
TTCACAGAAGAGAAGCTAGAAAATGGCGTCATCTCACCAGTAGCACCGCGAACATACCATTGCCCAATAT
CTTGGGGTAAGGAACGATAGATTGCCTCTCCTTGCAAATAAACTCGCTTAACACGTCCCTGATCTATGAA
ATCGTTAATATAAGATCCGCCCCAGGCCGATGACAGCGTATTACTAATATCTGCTTGGGCTAATCCAAGA
GCACTGGCTTTTCTTTGATCGATATTGACTTGTAACTGGGCTTTATCTTCTAAGCTATTCAAACGTACTG
CAGCTAGGCCAGAATCAGCATTTGCTGCTTTTAACACATTATTTTGTGCTGCGAGTAAAGCATCTCGTCC
TTTATTTTCAGCATCTTGCAACCAAAATTCAAAGCCACTAGATTGCCCAAGGCCTCTTACAGCAGCGGGT
GAAAGTACCTGAACACGTGCGTTACGTAGTGACTTAAAATATGCATTTGCACGAGCAATCACAGCTTCGG
CTGTATTTTCACTACCTTCTCGGTCATCCCAATGTTTGAGTCCGGCAAATGCCATTCCCACATTTTGCCC
ACTGCCTGCGTTATTTCTACCCATCACCATGAAAATCACATTGAGATTATTCTTTTCTTTCGTCAAAAAG
TAATCTGCAATCTGATTACCAACCCGCTCTGTTTCAGCTAGTGTCGTGCCTACAGGCGTACTAAATTGAA
CCATAACCGAGCCTTGGTCTTCTTGAGGTAAAAACCCCGTATTCATACGCGTATATTGCCAACCTAACAA
AGCTGTAATACCTACGAATATGACCATAAAGACTTTGGGTTTACCTAAAACAGCAACTAGCTTTGTGCGG
TAGCCTGTTTGACCTTGCTCTACTTTATAGTTAAACCAAGTAAAAAATTTTCTCTGCTTATGTTTTTCAT
TTGCAGGTTTGAGTAAGGTTGCACATAGCGCAGGTGAAAGTGTTAAAGCAACAATTGCAGAAAGTACCAT
AGCCGATACGAGCGTTACCGAGAACTGGCGATAGATAATCCCGACAGAACCACCAAAAAATGCCATAGGT
AAAAATACTGCTGCGAGTACCATGGCAATACCCACGAGCGCGCCAGATATTTCTTGCATTGAAATTAGGG
TGGCTTGCCGAGCATCAAGATTTTGCTCATGCATAACCCGCTCAACATTTTCTACGACGACAATCGCGTC
ATCTACAAGCAAACCAATTGCGAGTACCATTGCAAACAAAGTCAGTGTATTAATGCTATAGCCAAGTACG
CTCAACACACCAAATGTACCTAACAATACGACCGGAACTGCAATTGCCGGAATTAAAGTCGCACGCCAGC
TCTGCAAAAAATAAGAACATGACAATAATGACGAGAACAATTGCTTCAGCTAAAGTTTTAATCACCCCATT
TACCGAAGCTTCTACAAATGGCGTACTGTCGCGTGGGTATGCAACTTTTAAACCCGCTGGCATAGAGGCA
GTTAACCGTGTGACTTCAGCTTTAACTCGTTCAGCCGTTTCTAATGCATTGGCACCAGAAGCAAGTTGAA
TAGACATACCCGAAGCTGGCTTTCCATTCAAACGTGTAGAAGTCTGATAACTCTCTGCACCACGCTCGAC
TCGGGCAACATCTTTAAGTAATACAACCGCACCATTGGTTTGTGTCCGCAAAATAATATTTTCAAATTGG
CTCACGGTTTGTAACCGAGATAATGCTGTTACTGTTGCATTTAGAGCCTGCCCATCACGCGTGGGTAATG
CGCCCAATTCTCCTGCTGTAATTTGAGTGTTTTGTGCTTCAATTGCAGTTCTAACGTCTGATGGCATTAA

TCCGTAACTGTTTAATTTATGAGGATCAAGCCAGATGCGCATTGCATATTGAGCACCAAATACGGTGATT
TCACCCACTCCATCGACCCGACTTAATGGATCTTGCAGTGTACTCACCATATAGTCAGATATGTCGACAG
AAGATCTAGTCCCACTTTCATCGTAAAGCGCAAAGACCAGTAAACTATCGCCTTGTGATTTTGTCACAGT
GATCCCTTGCTGCTGGACCTCTTGAGGCAATCGACTCAGCGCCTGATTCACAGCATTTTGCACCTGCACC
TGAGCAGTGTCAGGATTGGTATTCTGATCAAAACTGAGACTTATCCGTGCTTGACCCGCCGAACTACTAC
TCGATGAAAAATAAAGTAGACCATCAATCCCTTTAATTTGTTGCTCCAAAATCTGTGTAACACTGTCTTC
AACAGTTTTGGCCGATGCGCCCGGATAATTTGCAGTAACATTTACACCCGGTGGCGCAATGTCAGGATAT
TGCTCAATGGGTAACGTTAAAATCGAAATCGTCCCCAACGCCATAATACAAATAGAGAGGACCCAAGCAA
AAATAGGTCGTGCAATAAAAAAACTCGACAACAT
>AciRND~~~adeF_baumannii~~~CU459141.1~~~AdeF periplasmic adaptor protein of adeFGH efflux pump in Acinetobacter baumannii
TTAACCTTTTGCCGGAGTTGATGTTTTATCTGTTGGCTGAGGTTGAGGAGGAGTAGCGCTAGCAGTGATT
TGTGAATTTGGCATAGGGACGAGATGCGGTGTAACAGGGTCACCCGGACGAATCCGTTGTAAACCATTCA
CTACAATACGATCACCCGCTTGTAATCCGCTATTTACGATTTGCAAGCCATCTTGTTGGGCACCGAGTTT
TACTTCGCGATAAGCAGTTTGATTTTTCGCATCTACTACTACGACAAAACGTTTATCTTGGTCGACACCA
ACCGCGGTTGGACTAATCAGAATCGCTGGGCGAGGTTGGCCTCCACCTAAACGAATTCGTGCATATAGGC
CTGGTAATAAAACACCGTTTGGATTGTCAAAAGTTGCGCGAACACGGATCGTACCTGAGGTTGTATTCAG
ATTGTTATCGATTGAGTTGATTGTACCTTCACGAGTAAAGCCTGTTTCATTGGCAAGTCCCATATAGACA
GGTACTTGTGCTGAATTACGCTGATTACTGATATATTTCAGGTAAGTTTGTTCATCAACATCGAAAGATG
CATAAAGGCGGGATACAGACACTAAACTTGTTAAAACCTGTGCGCCGTTACCTGCAGAAACTACATTACC
AACGGTGACTTCAGCTCGTGAAATCCGGCCGCTGACAGGTGCTGTAATACGGGTGTATTCTAGATTTAAA
CGTGCAGATTGGACAGCAGCTCTAGCGGCTTGTAGGTTAGCATTCGCTGAACGTGCATCATTTTCGGCTA
AATCCAGTTCTTGGCGAGAAACAGCATTACTCTGAATGAGACGTTGAATACGCGAAAGATTGCTTGCGGT
ATATGTTACCTGTGCTTCAGCTGAAGCAAGTTGGGCTTTTGCACGGTTCAGTTCTGCTTCAAAAGGACGA
GGGTCGATTGTGAAAAGTAAATCACCTTTTTTAACGAGGCTTCCATCTTTGAAATGTACGGCAATAAGTT
TTCCTGAAACTTGAGGCCGAATATCAACTTGATCAATTGCTTCTAAACGACCGGAATATTCTTGCCAATC
GGTAATGGTTTTGCTTACTACTGGGGCTACATCAACAGTAGCAGCTTGTTGGGCAGCGGTTGGTGCAGCT
TTTGCATCGGCATTTTCATGTAACAACATAAAACTGCCACCGGTTGCTAAAATAGCGACAAAGATGGCAG
ACAGTGCAAACTGTTTGCGGGAAAATGACAT
>AciRND~~~adeG_baumannii~~~CU459141.1~~~AdeG RND pump of AdeFGH efflux pump in Acinetobacter baumannii
TTAATGATCATGTGGGCTAGCTAACGGCGCTTCATGAACTGCCGCAGAATGCAGTTTATGTTTGCTGTTG
AGGGTACGAATCAGAACGTAAAAGGCCGGGGTGAGGAATAAACCAAAGAATGTTACACCGATCATACCGA
AGAATACGGCAACACCCATCGCATGTCGCATTTCAGAACCTGCGCCAGTTGAAGTAACCAGTGGCACTAC
ACCCATAATAAATGCAATAGAGGTCATTAAAATTGGGCGTAAACGTAGACGACTTGCTTCAACGGCTGCT
TTAAAGGCAGTCGCACCTTGCATTTCAAGTTCCCTCGCAAATTCGACAATTAAGATGGCATTTTTACAGG
CTAGCCCGACCAGTACCATTAGACCGATTTGAGTAAAGATGTTGTTATCTCCAGCTGTCAACCAGACACC
TGTCAGAGCCGCTAAGATTCCCATTGGTACAATTAAGATAACTGCTAATGGTAGGGTTAAGCTTTCATAC
TGAGCAGCTAACACTAAGAACACGAGTAATACGCTAATAGGGAATACCCAAAGTCCAGCATTACCAGCCA
AGATTTTTTGATAAGTTAAATCTGTCCATTCAAACTTGATACCACGCGGTAGAGTTTGTGCAGCAATACG
TTCAACCGCAGCTTCTGCTTGGCTAGATGAATAACCTGGGGCAGGGCCACCGTTAATATCTGCTGATGTG
TAACCGTTATAACGAACGACCATTTCAGGACCATAGGTTTGAGTTACATTCACCAATGAAGATAATGGCA
CCATTTGTCCGGCACTATTACGGGTTTTAAGCTGCAAAATATCTTCAGGGTTAGCACGGAAAGGCGCATC
GGCTTGTGCACGAACCTGATAAACACGTCCAAAGCGGTTAAAGTCGTTAACGTACTGAGAACCTAAATAA
ATCTGCATAGTATTGAAAACATCTGTCACAGCAACGCCTTGCTGTTTAGCTTTTACACGGTCCAGATCTA
CGTTGAGTTGAGGTACGTTAATTTGATAACTTGAGAACATTGGACCCAGTTCAGGGGCTGATTGTGCTGC
CTTCATAAAGTTTTGTGCAGCATCGTTCAAGGCTGAATAGCCTAAGGCACCTCGGTCTTCAAGTTGTAGT
TTAAAGCCGCCCATAGTACCTAAGCCCATCACTGGTGGCGGTGGGAAAACCGCGATATAGGCATCTTGAA
TAGCTGAATATTTCTGGTTGAGCGCACCTGCAATTGCATTTGCAGATAAGTCTTTTGCCTTACGTTCATC
AAATGGCTTTAAAGTCACAAAGACAATACCGGCACTTGAGCTATTGGTGAAACCGTTAATTGATAGGCCA
GGAAAGGCAACTGCACTTTCTACACCAGGTTGTTTAAGTGCAGTGTCACTCATTTTACGAATGACCGCTT
CGGTACGATCTAATGATGCGCCGTTTGGTAGCTGCGCAAAGCTAATTAAATATTGTTTGTCCTGCGCAGG
AACGAAACCACCTGGAACAATATAGGAAATACCAACGGTTAAACCTAAGAGTGCTGCATAGACACCCATT
GCCGAAGCTTTATGGGAAATGACACGGCTGACGCCTTGACTATAACGGTCTGAAGCACGTGAAAACACAC
GGTTAAACAGTGCAAAGAAACGACCGAATACACGATTCATAATACGTGTTAAGGCATCCGGTTTAGCATC
ATGTCCTTTCAGTAACAGCGCTGCCAAAGCAGGAGATAGGGTAAGCGAGTTAAATGCCGAAATAACCGTT
GAAATGGCAATGGTCATGGCAAATTGTTTATAGAATTGCCCTGTTAAGCCTGTCATAAAGGCAAGAGGTA
CGAATACTGCAACAAGTGTTAAAGCAATGGCAATAATCGGTCCACTGACTTCTCGCATGGCACGGTAAGT
CGCCTCCCTTGGGTTTAAGCCTGCTTCAATATTCCTCTCGACATTTTCGACGACCACAATCGCGTCATCG
ACGACAATCCCGATGGCAAGTACCATTCCGAACAGTGATAGCGCATTGATTGAGTAACCAAAAGCGAGCA
TGAGCGCGAATGTACCAATAATTGAAACCGGTACGGCAAGCAATGGAATGATTGAGGCACGCCATGTTTG
CAAGAATAAAATAACGACCACAACAACCAGTGTAATTGCTTCAAGTAAGGTATGAACGACCGCTTTAATA
CTTGCACGTACGAATTGAGTCGGGTCATAAACAATGTCGTATTTAATTGAAGATGGGAAATCTTTTGAAA
GCTCCTTCATTGTGCTACGCACTTGATCGGAAACTTGTAAAGCATTCGCACCCGGTGCTTGGAAAATTGG
AATCGCGACCGCTTGTTTATTATCAAGCAATGAACGTAAGCCATATTGAGAGGCTGCAAGTTCGACACGA
GCAACATCACCCAATCGGGTAACCGCGCCATCTGGTGCAGTTTTTAAAATGATATCTGCGAATTCTTGTT
CAGTAGTTAAACGACCTTGAGCATTGACTGAAAGCTGTAAAGGTGAATTACTTGGTGATGCACCGATTGT
ACCCGCTGCAACCTGAATATTTTGTTCACGGATTGCATTCACAATTTCGGTCGCGGTGAGGTTACGCTGC
GCTACTTTTTGCGGGTCAAGCCATACACGCATCGCGTAGTCACCAGAACCAAATAATCCGACTTCACCAA
CCCCTTGTAAACGCGCTAAACGGTCTTTCACGTTGAGCACCGCATAGTTACGTAAGTAGGTCATGTCATA
GCGATTATCTGGTGAGGTCAGATGCACTACCATAGTTAAAGTAGGTGAGCTTTTTAGTGTGGTTACACCT
AAGCGCTGTACATCTTCAGGTAAACGGGGCATGGCCTGAGACACACGGTTTTGAACCAATTGTTGGGCTT
TGTCTGGGTCGATACCGAGCTTAAAGTTCACCGTAATGGTTAGGTTACCGTCGCTGTTTGCTTGAGATTG
CATATACAGCATGTCTTCGACGCCGTTGATTGACTCTTCGAGCGGAGATGCAACCGTTTCAGCAATCACT

TTTGGGTTTGCACCCGGATATTGGGCGCGTACCACCACAGATGGTGGAACAACCTCGGGATATTCAGAAA
TCGGTAACTGAAATACCGAAAGGAGACCGGCGAGTAAAATCAAGACTGATAGCACACCAGCAAAGATCGG
CCGATCAATAAAAAATTTAGAAATATTCAT
>AciRND~~~adeH_baumannii~~~CU459141.1~~~AdeH outer membrane channel of AdeFGH RND efflux pump in Acinetobacter baumannii
TTAACTACTCCAACCGCCCCCTAAAGCACGGATTAAATTGATGCTTGCAATGATTTGGCTGCCTTTCAGC
TGAGCTGCTAATTGTTCTTGTTGCAAAATAGTGCGGTCAGAATCAATGACATCAAGATAGCTAATAGCAC
CTTCTCGATAACGTAAATGAGAAAGTTGATTGGCATGACGAGAAGAGGAGAGTGCTTGGTTTTGAGCCTG
AATTTGCTGATCGAGAATTCTTTGATCAGATAAACCATTTTCAACTTCGCGAAATGCATTCAGTACAGTT
TGTCTATAGTTGGCGACGCTTTCCTCATAAGCCGCTCTTGCTTGAGCAACGCCTGCTTTACGTTGTCCAC
CATCAAATAAAGGTAACGACAAAATAGTACCAGCGACAGGTCCTAGTAAAAAAGTCCGACTCGACCATTT
ACCCAACTCGCTTAAACTTGAAGATTCATAACCTAAAGCTCCTGTAAGACTGAGTTTTGGGAAAAATGCT
GCACGAGCAATTCCAATACGCGCATTATCTGCTGCCATTGCACGCTCTGCAGCCGCAATATCGGGTCGTC
TTTCAAGTAAAGTTGACGGCAAACCGGCAGGGAGACGGATACTATTTGCAGTTAAAGGTTGAACTGCCAA
GTTAAAGTCTGCTGGTGGTTTTCCTAATAAGACTGCAAGCGCATGTTCTGCACTGGCTCTGTTACGAGCA
ATATTTAGGGCAGTGGTTTGTGCGGTAGCAAGTTCGGTTTGTGCACGAGAAACATCTAATTCACTGACCA
GTCCGTTTTTAAAACGAAGCTGCATTAAATCTCGTGTTTCACCTAATAATTTGATTGTACGGTTATAAAT
TGCCTGTTCGGTATCAAGTTGACGTATCAGAAAATAACCTTGAGCTACATCCGCTTGTAGAGCTAAAAGT
GCCGACTGATATAGTGCCTCTTGTTGCTGTAGATCCGCTGTTGCTGCGTTGACACTACTTGCTACACGAC
CAAATAAATCGAGCTCATATGAAACATTGGCTTGAGCTCGCCATAAGGTTTGAGCCGAAGTATGTGCATT
GTCATCTAAACCGAGTGAAGCCGGAGACGGCTTTTGGCGGGTTGGCCCAAATCCGGCATCAATACTTGGT
AAGCGTTCAGCTTGAGCTGCCGAACGTAATGCACGTGAAGCCTGAATATTTGCTGCCACCGCTTTTAGGT
TCTGGTTGCCCGCGATAGCTTGCTGTTCAAGTTCATTCAGTTGAGCATCATTGTAAATGCGCCACCATTC
ACCACGAGTTTGCTGATCAGCAGGTTGGGCAATCTTCCAGTTATTATCTTCAAGTTTGGGGTCAGATTCT
TTGAATTTGACTGGCACTATAACTTTTGCAGGTTGATATTCTGGGGCCAATGAGCAGCCTGCAAGGAGCA
GGCTTCCCATGAGTGAGGACAACAACCAGTTTTGTTTTGATGTAATCACCAA
>AciRND~~~adeI_baumannii~~~CU459141.1~~~AdeI periplasmic adaptor protein of AdeIJK efflux pump in Acinetobacter baumannii
TTATGCATTTGAAGCTGCTTTCTGTTCTGCTTGAGGTTTACCTGATTGAGCCGGTTTCGCAGCATTTGGT
GCTGCACCTTGAGAGTTTGCTGGTTGAGCTTGATAAGGTTTTGCTGATACTTCTTGCCCTTCTTTAACTT
TGGCAACACCATCAACAATGACTTTATCGCCGGCTTTTAAGCCGTTAGTCACAATCCAGTTTTGTCCTTG
AACACCAGAGGTTTCAACAGGACGGCTCTCAACAACCCCTTTAGCATTAACAAGCATCGCTACAGCTTGT
CCTGTAGGTAAACGAGTAATGGCAGCTTGAGGAATCAGGTAAGCATTTGGAACAACGCCCTGAACAATTT
GCGCAGTGGTATACATACCCGGAAGCAATAAATGATTCGGGTTAGAGAATACGGCACGTAATGTAATTGT
TCCTGTATCTTGGTTTACAGAAGCGTCAGAGAAAGCAAGTTGCCCTTCGATTGGATAGGTAGAACCATCT
TCAAGCTTTAATTTTACTTTCGTGTTGTTACTGTTATTTAAACTGCCTTTGCTTAGTTGTTGACGTAAAC
GCAATAACTCAGCACTAGACTGATTAATATCAACATAGATAGGATCTAATTGTTGAATCGTTACCAACGG
GTCAGTCTGGTTTGCAGTAACCAAAGCACCAGCCGTTACTGAAGAACGACCAGATTGGCCAGAAATAGGA
GAGCGAATTGTAGAATAACCAAGATCTACATTTGCATTTGTTACTTGAGCCTTAGCTGCTGCAACTTGTG
CTTCTGCAACATTGACTTGACCAAGTAAGTCATCATATTCCTGTTTAGACACAGCATTACTAGAAACAAG
TTGTTTATAACGATTTAACTTGGTACGTAGTGAAGCTAGATTTGCCTGTTGTTGTAGGAGTGATGCTTTT
GCATTTTCTAACGTTGCACGGTTCGTTCTAGAGTCGAGCTCATAAAGCGCCTGACCTTCACGAACATAGC
TTCCTTCAGCAAATAAACGTTTTAAAATCACGCCACTTGTTTGAGGACGAACTTCAGAAATTTGATATGC
TGAAGTACGGCCTGAAAGCTCAACGCTTTGTTCAACACTTTGTGGTTGAGCAACAATAACACCTACTTCT
GCAGGCGGCATTTTCTGAGCAGCAGCAGCTTGCTGTTTCTCATCGGAGCCTTTGCTACAACCAACAAGCG
CGATACTTGTTGCTAATGCGCAAGCAGTAAGGGCTGGTGCCCAAAGCTTAGCCGACATCAT
>AciRND~~~adeJ_baumannii~~~CU459141.1~~~AdeJ RND pump protein of AdeIJK efflux pump in Acinetobacter baumannii
TCACGATTTATGCTCCTGAGTGTTTATGGTTTTTGGTTTGTACTTAAAGATACTACGAATCCACACATAG
AATACAGGGATAAAGAAGATACCTAAGAACGTCGCGCTGAGTACGCCACCAAGTACACCAAAGCCTACAG
AGTGCTGACTTCCTGCACCGGCACCTGTTGAAAGTGCAAGTGGAAGTACACCGAAACCGAAGGCTAGGGT
GGTCATGATAATTGGACGTAAACGCATTTTTGCAGCATGTAAGGTTGCATCAAGTAGATCTTCACCTTTT
TCCTGCAATTCTTTTGCGAATTCAACAATCAAGATCGCATTTTTTGCAGAAAGACCGATAACCGCAATAA
TCGCTACCTGGAAGTAAATGTTATTTGAGAGATTTGGATCTCCTTTAATAATCATGCCCAAGTAGGTCAA
TACGATTGCACCAATGACACCAAGTGGTACCACAAGTAAAACCGAGAACGGAATTGACCAGCTTTCATAT
AGTGCAGCCAAGCAAAGGAATACGATTAACAATGAAAGTGCGTATAAGAACGGCGCTTGAGCACCAGACT
CACGTTCTTCAAGTGATAAGCCTGTCCATTCATAGTCGAAACCTTGTAAGCCCATAGATGGTAACTTACC
AATAATTTCTTCCATTGCTTTCATGGCATCACCAGAGCTAACGCCAGGTGCAGGTGTACCTTGAATGTTA
ACCGATGACACGCCGTTATAACGTTCGAGACGTGGAGAACCATACGTCCATTCACCTGTAGCAAATGCCG
AGAATGGAACCATCTCACCTTTGTTATTACGTACATACCATTTGTTTAAGTCTTCAGGCATCATACGGCT
GCCCGCATCACCTTGAACATAAACTTTTTTCACACGACCACGGTCAACGAAATCGTTAATGTATGAGCCA
CCCCATGCAATACGCATTGTATTGTTGATTTCGGCAATACTAACGCCCATAGCACCAGCTTGAGCCTGAT
CTACATTAATTTGATACTGAGGAGTATCTTCCTGACCATTTGGACGCACACCTACAAGACGTTTATCTTG
TGATGCCAAACCTAAAATCGTGTTACGAGCTGCAATCAGTTTCTCATGGCCTTGACCACTTGAATCTTTA
AGCTGCAAGTTAAATCCGGCAGTTACACCAAGTTCAGGCATTGCTGGAAGCTGTAACGGCATAACGTATG
ATGCATCTTTAATGATCATATTTAATGCCATACCACGCTGAATCAATGAACCAATTTGAGTTTCTGGTGT
CGTACGTTTGCTCCAGTCTTTCAACTTAACGAAGCCAATACCCGCGTTTTGACCAACACCTGTGAATGAG
AAACCAGAAACAGTGAAAATAGATTCCACGGTATCTTTTTCATTCATAAAGAAGTTAGTCATGGTATCAA
TCACTTTACCGGTACGGTCAAGTGTTGCATTTGGTGGTAATTGTACAAGTGTCATGACCACACCCTGATC
TTCTTCTGGTAAGAATGAAGACGGGAGTTTTTGGAACAAGAAGACTAAAAGGGCAACTACAACAGCATAG
AGCACGCCAGAGAAGATTTTGCCTTTAAGCATGCGGCTAACACCATTTTGGTAGCTATGCGACATGCGGT
CAAAACCATTGTTAAAGCTTCTAAAGAAACGCGCAAAGATATTATTGCTTGGTTCTTTATTAGGATCATG
CTGTTTCAAGATAGTTGCACAAAGTGCCGGTGTGAACGTCAACGCTACAATTAACGACAGAACCATTGCA
GTTACAAGGGTAATCGAGAACTGGCGGTAAATTACACCTGTTGTACCACCAAAGAAAGCCATTGGTACGA

ATACCGCTGTCAATACGCTGGTAATACCTACTAACGCACCAGAAATCTGCTGCATTGAGCGAGAAGTGGC
GGTAACCGGATCGGTATGGTCTTCACTCATCACACGTTCAACGTTTTCGACTACAACAATGGCGTCGTCG
ACCAGAAGACCGATTGCCAATACCATAGCGAACATGGTTAAGGTGTTAATTGAGAAGCCAAAGATATTAA
TGACCGCAAATGTACCTAATACAACTACTGGAACTGCAAGCGTTGGAATAATCGTTGCACGCCAGTTTTG
TAAGAATAGGAACATGACAATGAATACCAAAATCACGGCTTCAATTAATGTGTGTACTACACTTTCAATT
GAAAGACGGATAAATGGAGTCGTGTCATAAGCCAGTTTATCTGCAAGACCTGTTGGATAGTTCTTACGTA
GTTCAGATAAACGTTGTTCAACTGCTTCGGCTGTGTCGAGTGCGTTAGCACCTGTTGCAATTTTAATTGC
AAGACCAGCTGCCGGTTTACCGTTAAACTTCGAGTCGAATTGATAGTTATCCGAACCTAATTCTACGCGA
GCAACATCTTTTAATCGAACCTCAGCACCTGATGCTGTGTTCTTTAAGAAGATATTTTTAAACTGTTCAG
GAGTCTGCAATAAGCTTTGTGCATTTACTGTTGCGTTAAGAACTTGACCTTGTACAGCCGGAGCACCACC
TAACTGACCTACGGCAACTTGCGAGTTCTGCGCTTGTAAGGCAGTTGCAATATCACTAGGCGTAAGTTGA
TAGCTTGTTAATTTAGCTGGATCAAGCCAGATACGCATTGCGTATGAGCCACCGAAGACCTGTACCTCAC
CAACACCGGCAACACGGCTAAGCGGTTCTTTAATTGACGAGTTTACATAGTCTTTAATATCAGAGTCTGA
CAGGTTGTTATCTGGTGAATAGAATGCAATAACTTGCAAGAAGCTCGCACCAGATTTAGTAACTGTTACA
CCTTGACGTTGTACATCTTCAGGTAAAAGCGCAGTTGCAGATTGCAATTTGTTTTGAACTTGAACCTGTG
CAATATCAGGGTCAACACCTTGTTCAAAGTTTAATTGAATAGATGCCTGACCATTACCAGCACTGTTAGA
TGAAATATAACGTAAGCCATCAAGACCATTCATTTGTTGTTCAATGATCTGGGTTACAGTATTTTCAACT
GTTTCAGCCGATGCACCAGGATAAGTCGCAGCAATCGTTACGGTTGGTGGTGCAATCGTTGGATATTGTG
CAATAGGCATTTTTGTTAGCGTAAGAATACCCGCCAACATAATGACTAATGCAATCACCCACGCAAATAT
GGGGCGATGAATAAAAAATTGTGCCAT
>AciRND~~~adeK_baumannii~~~CU459141.1~~~~AdeK outer membrane channel protein of AdeIJK efflux pump in Acinetobacter baumannii
TTATTGCTTTTTAAGTTCAGCACTAGATGGTTGATGTACCACTGTATCTGAAGTATTTGCTTTTAAACCG
CCACCTAGAGTTTTGTATAACTCGATTTGGTTGTTTAAGTTTGCCTGTTGAAGCAATAACAAACCTTGTT
CAGCCGCATATGAAGAACGCTGTGCATCAAGAACTGTCAAGTAACTATCAATACCAGCACGGAAGCGGGC
ATTTGAAAGTGTGTAGTTGCGGTTAGTTGCTTCTACTAGACGTTGCTGAGCGGTTAAACGCTCGCCAATA
TTGGCACGAGTTGCAAGTGCATCATTAACTTCACGGAAAGCCGACTGAACTGATTTTTCATAATCAGACA
ATGCAATTTTCTGATCGGTTTCAGAAATTTTTACATTGGCACGGCGTGTACCCCAGTCAAAGATTGGTAA
ATCTAAGCTTGGACCAACTGACCAGACAAAACCACCAGACTTAAATAGATCACTTAAGTCAGTTGATGCA
TAACCAGCCGAACCCGTTAAGCTAATGGTTGGGAATAAACGTGCTTTTGCTGCACCGATATTCGCACCCG
CAGCGCTTAAGTTGTATTCAGCAGCTTTTACATCTGGACGGTTATTTAACAAGTCACTTGGTAAACCGGC
AGTAAACACATTTTGTTGAGCAATGCGTTTTACAGGTTGTGTAGGTAACAAGTTTTGTGGAACAGGTTGG
CCTACAAGCAAGTTCAACAAGTTTTGTGCTTGAGCAATTTGAGTTTTGTAGTTCGCTACATCATTACGCG
CAGTTTCTACAGAAATCTGTGCTTGACGTAATGGAACTTCACTGTCGATACCTACATCAAAACGTTTTTT
GTTGAGATTGTAAGAATCTAACTGTGCTTTAAGCGTTTGCTCTGCCAGTCTTAAGTTTGCTGTTGCAAAC
GAATAATTTAACCATGCTTGAGCAACTTGGCTAATCAGACTGATTTGAGTCGAATCACGAGCACTTTGTG
TTGCAAGATAACTATCTAAAGCAGCATCTTTGAGGCTACGAACACGACCCCAGAAATCTAGCTCATAAGC
AGTTACACCCAAACCTACTTGATAAGTAGAGTAGGGGTTATTCGGATCACGGCTTTGAGAAACCTGACGA
ATTGCACTACCACTTGCACCGATTGTTGGAAGCTGGTTGTTCTGTGTAATCTGATATTGCTGTTGAGCAC
GTTCAATATTGAGCGTTGCTGTACGTAAGTCACGGTTATTGGCAAGAGCCAAATCAATCACTTCAAGCAA
ACGCGGGTCAGCAAAGAACTGTTTATAACCCTGTTCAGCAATAGACGTACCAGAAGCGCTGTTATATGCA
TAGCTTTGTGGTATATCGGTTTTCACGACTGGTTCTGGGCCGCGCATGCTTTGACAAGCTGCCAAAGCAA
GCGCAAGTGCAGATACCGCAATGCTACGACCTGAAATAGACCATACTTTTTGCATCAC
>AciRND~~~adeI_lwoffii~~~GCA_900444925.1~~~AdeI periplasmic adaptor protein for AdeIJK efflux pump in Acinetobacter lwoffii
ttatgcatttgcagcaggtttttgttcaggtttctctttagcatcatctgcctttttctcgctcgttgcaggctgttgcggtgcagcaccttg
tggcgcagcagcttgaggttgataaggttttggtacaacttgctgttcaggttttactttcgcaataccatctacaatcactttatcgcctgg
attcagaccttcagtcacaatccagttgctgccttgggaaccaacggtagtcactggacgtggcgtcactttattgtcagcaccgaccagcat
cgccatcgcctgacctgttggggtacgggtcaaagctgcttgcggaatcaggaacgcattcggtacttcaccctgtacgatttgtgcagtggc
aaacatacctggcaaaagcaggtgatttttgttcgggaataccgcacgtaaagtcacggtacctgtttccgggttcacgctggcatcagagaa
tgcaagacggccttcaaccgggtagatactgccatcttcaagcttcaacttgactttggtgttgttcgagctgttcaagctgccttggttaag
ctgttgacgtaaacgtaaaagctcggcacttgactggttgatatctacatagattggatctagacgctgaattgtcaccaatggctctggctg
attcgctgttaccaaagcacccgccgtcaccgatgaacggttggtctggcctgaaatcggagcgcgtacagtcgaatagcccaggttgatttc
ggcattacgcaaagttgcgcgattggcattcaaatcagcttcagccagtttgacctgtgccgcgatgtcatcatattcctgtttggaaatggc
attggtaccgaccagttgacggtaacggccttccttcacacgcaatacgcccagattggcttcagcacgggcaatcgctgcacgagcattgtc
cactgtagtgcggttggtgcttgaatcgatttcatacaaggcctgaccttcacgcacatagctgccttccgtgaacaagcgtttaagcaccac
gccactggcttgtggacgtacatctgaaatttcaaatgctgttgtacgacctgagagctctacagactgctctacactttgcggttgtgcaac
aagaacaccgacttcagtcggtggcatttgttgttgagcaccagcttgctggccctctttaggatctttgctacaaccaacaagtgcaagact
tgttgccaatgcgcaagcagtcagggcaggtgcccaaagctttgcagacatcat
>AciRND~~~adeJ_lwoffii~~~GCA_900444925.1~~~AdeJ RND pump protein for AdeIJK efflux pump in Acinetobacter lwoffii
ttacgacttatactcctgatttgttgttttggtttgtatttaaagacacttcgaatccacacgtagaataccgggataaagaagatacctaa
caaggtcgagctgatcacgccgccgagtacgccataaccgactgagtgctggctacctgcacccgcccctgaagccagagcaagcggtaatac
accaaaaccaaaggccaatgtggtcatgataattggacgtaaacgcatttttgccgcatgcaaggtcgcttcaaataactcttcaccttgttc
ctgcaattctttcgcaaactcgacaatcaagatcgcgtttttcgcagaaagaccaatcacggcaataatcgctacctggaagtaaatgttatt
cgacaggtttggattttgcagcagcaccattccaccaaaggtgagtaataatgcaccgacaatacccaatgggaccaccagtagaaccgagaa
cggaatcgaccagctttcatacagggctgccagacacaggaagacaatcagcagtgacaaggcatacaggaacggagcttgtgcaccagattc
acgctcttctaaagacaggcctgtccattcgaagtcgaaaccttgcaagcccatttctggtaacttagccacaatttcttccatggccaccat
tgcgtcgcccgagctgatacccggagcaggtgtaccttgaatgttcatggacgatacgccgttataacgctcaagacgtggtgaaccgtaggt
ccattcgcctgtggcaaatgccgagaatggcaccatttcaccacgattattacgcacataccattggtccagatcttccggcatcatacgtga
gccagcttcaccctgaacatagactttcttgacacgaccacgatcaatgaaatcattgatatatgaaccgccccaagcaatacccatggtgct
gttaatttcggcaacgctgacacccagtgcacctgcttgcgcatgatccacaatgacacggtactgaggagtatcttcctgaccgtttggacg
tacgcctgccagacgcgaatcttgtgcggccatacccaggattgcattacgtgctgccaggagctgttcatgactttgaccacttgcagcctt
gagctgcaagttgaagccagccgacacacccagttcaggcatcgcaggaagctgcaatggcatgatataagacgcatctttgataatcatgtt

cagtgccataccacgttgaataatggcaccgacttgagattccggactgtgacgctctgaccagtctttcaacttaatgaaagccagacctgc
attttgtcccacacccgtgaaggagaaacctgccacactaaagacggattctacatgctctttctcgttttccaagaaatagttggtcatggt
gctgaccactttgtcggtacgttccaaggttgcatttggcggtaactgaaccagcgtcatgaccacgccctggtcttcatccggtaggaacga
agaaggcagtttctggaacaacagcacaataatacccagtaccacggcatacaccacacctgagaaaatcttgctgtgggtcatgcggttgac
gccgccctgatatttcacagagaccttgtcaaaactgctattaaaccagcggaagaaacgcgcaaaaataccattactttccggcttgttcgg
atcatgctgtttcagcagggtcgcacacagggcaggagtaaaggttaaggcaaccaccagtgacaagaccatcgctgtgaccagcgtaatcga
gaactggcgataaatgaccccggtggttccaccgaagaaagccattggaacgaatactgccgacagcaccgaggtaataccaatcaatgcgcc
agaaatctgtttcatcgatttttcagttgcagcaaccggatccagatgttcttccaccatgacccgttcgacgttctcgaccacaacaatcgc
gtcatcgaccagaagaccaatcgccaataccatggcaaacatggtcagggtattaatcgagaagccaaagatgttaatgacggcaaatgtacc
caatactacgactggaactgccatggttggaatgatggtggcacgccagttctgcaagaacaggaacatcaccaggaatactagaataatcgc
ttcgatcagagttttaactacactcttaatcgacagttcaataaatggtgttgtatcaaacgccagttggtcgaccatgccttgcggatagtt
cggacgtaactgttttaagcgttcttcaaccgcttgtgcggtatccagggcattggcgccagtcgcaagcttgattgctacaccaccagccgg
tttgccattaaatttagaatcgaactgatagttatctgcaccgagttcaactcgtgcaacatcacctaagcgaacctgcgcaccagaagtggc
atttttcaggaagatattgctgaactgctccggcgtttgtaacagactttgagccgttaccgtcgcatttaacacttgtccttgaactgcagg
cgcaccacccaactgacctacggcaacctgtgcattctgggtacggattgcattagcaatatcacttggcgttacttgcaggctggccatctt
ggctggatccagccagatacgcatcgcataagaaccaccaaagacctgaacttcaccgaccccggccacacggcttaaaggttcagcaatatt
cgagttaacatagtctttgatgtcatctgcggtcaggctgctatcaggtgaatagaatgccaaaacttgcaagaagcttgcacccgatttggt
cacccgcacaccctgacgctgtacatcttcaggcagcagggcagtcgcagactgtaatttgttttgaacctgaacctgtgcgatatcaggatc
gacaccctggtcaaaattcaaggaaatagacgaagagccgttacccgcactgttagacgagatatagcgcaagccatccataccgttcatctg
ctgctcgatgatctgggtcactgtattttcaacagtttcagcagatgcacccggataagtcgcagaaatcgtcacggtcggtggcgcgatggt
tggatattgtgcaatcggcattttgctgatggttaaaatacccgccagcataatcaccaatgcaatcacccatgcgaaaatcgggcgatgaat
aaagaattgagccat
>AciRND~~~adeK_lwoffii~~~GCA_900444925.1~~~AdeK outer membrane channel protein for AdeIJK
efflux pump in Acinetobacter lwoffii
ttattgcgctggttgttctgccgcacggcgctgttcagatacagacggtgtattcggtaaggtatccgtcgtattgactttcaagccgccccc
cagggttttgtagacttcaacctgattgttcagattggcctgttcaagcaataacaaaccttgttctgctgcataagacgcacgctgtgcgtc
cagtaccgtcagatagctgtcaattccggcgcggaaacgtgcctgtgacaagcgataggtggtgttggttgcttcgaccaggcggcgctgagc
tgcaagacgatcaccaatattctgacgtaccgccaaagcatcattcacttcacggaaagcagtctgaatcgacttttcgtaatctgacaaggc
aatttgctgatcggtttcagaaatacggatattggcctgacgggtaccccaatcaaaaattggaatatccagactcggaccaaatgaccaggc
aaaactgcccgacttaaacaggtcgcctaaatccgtagaggcataacctgctgaaccggtcaaactaatcgtcgggaacaggcgtgcacgtgc
tgccgcaatattggcacctgccgcagacagacggtattctgctgcacgaacatctggacggttaatcagtagatcacttggcaagcctgaacc
caatgcagtatttgaggtgatgcgggttacgcgttgtgctggtaataaattcgcaggcacttgctgaccgaccagcaggtttaacaggttttg
tgcctgtgctacctgagtgcggtaattcgcaacatcattacgtgcagtttcaaccgagatctgtgcctgacgtactggaagctcgctgtcgat
cccgacatcaaaacgtttcttgttcagattgaaagattcaagctgggctttcagcgtttgttcagccaaagccagattggcattggcaaacga
gtagctgagccaagcctgagacacctgaccaatcaaggcaatttgcgtcgcatctcgcgcactttgggtcgccaggaaactatccagcgcatt
gtcacgcaagctgcggacacgcccccagaaatccagttcatacgcagttacacccagtccgacattataagtcgtcactgcaccacgattctg
ggaattcatggtgtcctggcgcaatacgctaccactggcaccaatcgtcggtaactggttgttttcagtaatctgatactgttgctgggctcg
ttgaatattgagggcagcaatacgcaaatcacggttattgtttaatgccagatccagcacctgtaacaggcgctgatcggcaaaaaagttttt
ataaccttgttctgctacagaagtaccagacgcattggtatagctgctcgggatatctgcctgcgcaaccggctctggactgcgcatgctttg
acaagccgtcaaagcaagcgcaagggcagataccgcaatgctacgacctgtaatagaccatacattttgcat
>AciRND~~~adeX_baylyi~~~CR543861.1~~~AdeX periplasmic adaptor protein of AdeXYZ efflux pump
in Acinetobacter baylyi
ATGACGTCGGCTAAGCTTTGGGTACCTGCCCTTACTGCTTGCGCATTGGCAACAAGTATTGCACTTGTTG
GTTGTAGTAAAGATCCTAAAAATGCTGAACAGGCTGGTGCTGCTCAGCAAATGCCACCACCACAGGTGGG
TGTGATCGTTGCTCAACCGCAAAGCGTAGAACAATCAGTAGAACTTTCTGGACGAACATCTGCTTACGAG
ATTTCTGAAGTTCGACCACAAACAAGCGGAATCATTTTAAAACGCTTGTTTGCAGAAGGTAGCTACGTTC
GTGAAGGACAGGCGCTTTACGAACTAGATTCACGTACCAATCGTGCGACGCTCGACAATGCAAAGGCAGC
ACTTTTGCAACAACGGGCAAATCTAAACTCGCTTAAAACCAAGATGAATCGTTATAAACAATTGGTTTCA
AGTAATGCGGTTTCCAAACAGGAATACGATGATCTGGTCGGTCAGGTCAATGTAGCTGAGGCACAAGTTG
CAGCGGCACAGGCACAAGTGCAAAATGCGCAGGTTGATTTAGGTTACTCTACCATTCGCTCACCAATCTC
GGGTCAATCTGGTAAATCATCCGTAACTGCTGGTGCATTGGTTACAGCAAATCAGACCACACCACTGGTG
ACCATTCAACGAATGGATCCGATTTATGTAGATATCAATCAATCAAGTACAGAATTACTTCGCTTGCGTC
AGCAATTGAGTAAAGGTAGTCTGGACAACAGTAACAATACTAAGGTTAAACTTCGTCTGGAAGATGGTTC
TATCTATCCAGTTGAAGGGCGCTTGGCATTTTCTGATGCAAGTGTAAGCACTGATACAGGTACTGTTACC
CTACGTGCTGTTTTCCCGAATCCTAATCATTTATTGCTTCCTGGTATGTATGCAACTGCACAAATTGCTC
AAGCTGTCATTCCAAATGCATATCTGATTCCTCAAGCTGCGGTTACACGTACACCAACGGGTCAAGCAAC
TGCAATGGTAGTCAATGCTAAAAATACCGCAGAAACACGTCAAATTCAGACTGCTGGTGTACAGGGTAAA
GACTGGATTGTCACTGAAGGGCTTAAATCAGGTGATAAAGTGATTGTGGATGGTATTGCCAAGGTTAAAG
AAGGTCAGCAAGTTACTGCTCAACCTTACCAAGCACAACCA
>AciRND~~~adeY_baylyi~~~CR543861.1~~~AdeY RND pump protein of AdeXYZ efflux pump in
Acinetobacter baylyi
ATGGCTAAATTTTTTATTCATCGCCCCATCTTTGCGTGGGTGATCGCACTGGTCATTATGCTGGCGGGTA
TTCTTACTCTCACCAAAATGCCAATTGCGCAATATCCGACGATTGCTCCACCTAAGGTTACAATTTCTGC
AACCTATCCTGGTGCATCTGCGCAAACAGTTGAAAATACAGTCACACAGATCATTGAGCAACAAATGAAT
GGTCTGGATGGCTTACGCTATATCTCGTCGAATAGTGCGGGAAATGGTCAGGCCTCTATTGAATTAAACT
TTGAACAAGGTATCGATCCTGATATCGCACAAGTTCAGGTACAAAATAAACTTCAATCTGCAACGGCATT
GCTTCCAGAAGACGTGCAACGTCAAGGCGTGAAGGTCACTAAGTCTGGTGCCAGCTTCATGCAAGTTATT
GCATTTTACTCACCTGATGGAAGTTTATCGGGTTCAGACATTAAAGATTACGTAAATTCAAATATTTCTG
AACCGCTTAGCCGTGTAGCGGGTGTGGGTGAAGTTCAGGTATTTGGTGGCTCTTATGCCATGCGTATTTG
GCTTGATCCTGCAAAACTTACCAGCTATCAACTCACACCAAGTGATATTTCTACTGCCTTGAATGCACAA
AATGCTCAGGTTGCAGTGGGTCAACTCGGTGGTGCACCAGCTGTTGATGGTCAGGTATTAAATGCCACGG
TGAATGCTCAAAGTCTTTTACAAACACCAGAACAGTTTCGTAATATTTTCCTGAAAAATACCAGTACGGG
TGCAAAAGTTCTGTTAAAAGATGTCGCACGCGTTGAGTTAGGATCAGACAATTATCAGTTTGATTCCAAG
TTTGATGGAAAACCAGCAGGTGGTGTGGCGATCAAACTGGCAACAGGAGCAAATGCGCTCGATACTGCTA

AAGCAGTAGAAGCCCGTTTGGTAGAGCTACGCAAAAACTATCCAGCTGGTTTAAAAGATAAGCTTGCTTA
CGACACCACACCTTTCATCAGACTTTCAATTGAAAGCGTAGTACATACGCTCATTGAAGCTATCGTATTG
GTATTTATCGTGATGTTCCTGTTCTTGCAGAACTGGCGTGCCACGATTATTCCGACACTTGCAGTTCCAG
TGGTGGTACTCGGAACCTTTGCCGTTATTAACCTTTTTGGCTTTACGATTAATACTTTAACCATGTTTGC
TATGGTACTGGCGATTGGTCTTCTAGTCGATGACGCAATTGTGGTGGTTGAAAACGTCGAACGAGTCATG
TCTGAAGAGCATACCGATCCAGTTACAGCCACTGAGCAATCAATGCAACAGATTTCAGGTGCATTGATTG
GTATTACCAGCGTATTGTCTGCGGTATTCATTCCAATGGCATTCTTTAGTGGTACTACAGGCGTTATTTA
TCGTCAGTTTTCAATCACACTGGTGACCGCAATGGTTCTTTCCTTAGTGATTGCCCTGACCTTTACGCCA
GCACTTTGTGCTACTTTGCTCAAACAGCATGATCCGAACAAGAAAGAAAGTAACCATATTTTTGCGCGCT
TCTTTAGATGGTTTAACAGAACCTTTGATCGTCTGTCACACGGTTATCAAAATGGCGTAAACCGCATGCT
GCATCATAAAATTCTATCTGGAATTCTTTATGCTGCCATTTTCGGAATTTTGGTGTTGTTCTTTGTCAAA
CTCCCTTCTTCATTCTTGCCAGAAGAAGATCAGGGCGTGGTTTTGACACTGGTTCAATTACCACCGAATG
CCTCACTTGACCGTACTGGTAAAGTCATTGATACCATGACCAATTATTTCATGAATAAGGAAAAAGATTC
GGTCGAATCAATCTTTACTGTTGCTGGCTTCTCTTTTACTGGGGTCGGTCAAAACGCAGGTCTAGCCTTT
GTTAAATTGAAAGACTGGAGTGCACGTACCACACCTGAAACCCAGATTGGCAATATCATTCAGCGTGGTA
TGGCACTCAATATGATCGTGAAAGATGCATCATATATTATGCCATTACAGCTTCCTGCTATGCCTGAACT
GGGTGTAACTGCAGGCTTCAACTTGCAACTTAAGGATTCGAGTGGTCAAGGCCATGACAAATTGATTGCT
GCACGTAATGCTGTACTGGGTATGGCATCACAGGATAAACGCCTCATGGGTGTACGTCCAAATGGTCAGG
AAGATACTCCTCAGTATCAAATCAATATTGATCAGGCACAAGCGGGTGCACTGGGTGTCAGTATCGCTGA
TATCAACAATACCATGAGTATTGCATGGGGTGGTTCATACGTGAATGACTTTGTTGATCGCGGCCGCGTG
AAAAAAGTGTATGTTCAAGGCGACTCTCAAAGCCGAATGATGCCTGAAGACCTCAACAAATGGTATGTGC
GCAATAGCGCAGGCCAAATGGTTCCATTCTCTGCTTTTGCGACAGGTGAATGGACATATGGTTCACCACG
TCTTGAGCGTTATAATGGGGTATCTTCAGTGAACATTCAGGGTTCACCTGCTACAGGTGTAAGTTCTGGT
GATTCAATGAAAGCCATGGAAGAGATTGTCAATAAACTTCCATCCATGGGACTTCAGGGTTTTGATTACG
AATGGACAGGTCTATCGCTTGAAGAACGTGAGTCTGGAGCGCAAGCACCATTCTTATATGCCTTGTCATT
ATTGATTGTATTCTTATGTCTGGCAGCCTTGTATGAGAGCTGGTCTATTCCATTCTCAGTTCTGCTGGTG
GTTCCGTTAGGTGTAGTTGGGGCAGTATTACTGACCTATTTAGGTATGGTCATTAAAGGCGATCCAAACT
TATCCAATAATATTTATTTCCAAGTTGCTATGATTGCGGTGATTGGTCTTTCGGCAAAAAATGCGATTTT
GATTGTTGAGTTTGCAAAAGAGCTTCAAGAAAAAGGTGAAGACCTCCTTGAAGCAACCTTGCATGCCGCA
AAAATGCGTTTACGTCCAATTATCATGACGACTCTAGCCTTCGGATTTGGTGTACTTCCTCTCGCACTGG
CCTCTGGTGCAGGTGCAGGTAGCCAGCATTCAGTCGGTTATGGTGTACTTGGTGGCGTAATCACTTCAAC
GCTTTTAGGTATCTTCTTTATCCCAATTTTCTTTGTGTGGATTCGGGGTGTCTTTAAGTACAAACCAAAA
ACCGTTAATAATCAGGAGCACAGATCGTGA
>AciRND~~~adeZ_baylyi~~~R543861.1~~~AdeZ outer membrane channel of AdeXYZ efflux pump in
Acinetobacter baylyi
GTGATGCAAAAAGTATGGTCTATTTCAGGTCGTAGCATTGCGGTATCTGCACTTGCGCTTGCTTTGGCTG
CATGCCAAAGCATGCGTGGCCCAGAACCAGTTGTCAAAGCCGACATTCCGACAAGTTATACCTATAACGC
GTCAGGTCAATCTATTGCAGAGCAAGGCTATAAAGACTTTTTCTCTGACCCTCGCTTATTGCAAGTGATT
GAGTTGGCTCTCAACAATAACCGTGATCTACGTACCGCTGCATTGAATATTCAGCGTGCACAACAACAGT
ATCAAATTACTGAAAATAACCAGCTTCCAACAATTGGAGCAAGTGGTAGTGCAATCCGTCAGGTAAGCAT
CACACGTGATCCCAATAATCCATACTCCACGTTTCAAGTTGGACTAGGCGTGACTGCCTATGAGCTTGAT
TTCTGGGGGCGAGTACGCAGTCTGAAGGATGCAGCACTCGATAATTATCTGGCGACACAAAGCACACGTG
ACACCACTCAGATCAGTCTAGTGAGTCAGGTGGCTCAGGCGTGGTTAAGCTACTCTTTCGCCGTTGCAAA
TTTGAAGCTGGCTGATCAAACACTGAAAGCTCAACTCGATTCTTATAATCTGAACAAAAAACGTTTTGAT
GTAGGTATTGATAGTGAAGTTCCTTTACGTCAAGCTCAGATTTCGGTCGAAACTGCACGTAACGATGTAG
CCAACTATAAAACGCAAATCGTTCAGGCTCAGAACTTACTCAATCTTTTAGTAGGTCAGTCTGTTCCTCA
GAATTTGCTACCGAATCAACCTGTCAAATCTATTACACGTCAAAATGTATTTAGTACAGGCTTACCAAGT
GACTTGTTAAACAATCGTCCAGATGTAAAATCTGCGGAATATCAACTGAGTGCGGCAGGTGCCAATATTG
GTGCTGCCAAAGCTGCGTTATTTCCAACAATTAGCTTAACAGGTTCAGCAGGCTATGCATCTACACAATT
AAGCGACCTGTTTAAATCTGGCGCTGGTGTCTGGTCAATTGGCCCAAGTCTTGAATTACCAATTTTTGAT
TGGGGTACGCGTCGTGCCAATATCAAAATTTCTGAAACTGACCAAAAAATTGCTCTTTCTAATTATGAAA
AGTCAATTCAGTCAGCTTTCCGTGAAGTGAATGATGCTTTAGCAACGCGTGCCAATATTGGTGACCGTAT
CTCTGCCCAGCGCCGTCTGGTAGATGCAACTAATACCACATATAGCTTATCAAATGCCCGCTTCCGTGCA
GGTATCGATAACTATTTGACTGTATTAGATGCACAACGTACCTCCTACTCGGCTGAACAAGGTCTTTTAC
TGCTTGAACAAGCGAATCTGAATAACCAGATTGAACTCTACAAAACCTTAGGTGGTGGTCTAAAAGCCAA
CACCACGGATACTGTGGTTCATCAGCCTTCAAGTGCAGATCTGAAAAAGAAATAA
>AciRND~~~czcA_baumannii~~~CU459141.1~~~CzcA RND pump protein of CzcABCD efflux pump in
Acinetobacter baumannii
ATGGACATTAATAAGCCTGAGCTGCCCAAGCCAGAAGGACTGTTTGACAGAGTTATTCAGTTTGCTATTC
AAAATGCCATTTGGGTCATGCTTTTTGTTCTGGCGTGGATTGGTGTTGGCATTTGGAGTTATCAAAAACT
CCCTATTGATGCCGTTCCAGATATTACCAATACCCAAGTCCAGATTAATACTCAAGCCAATGGGTATACC
GCTTTAGAGGTCGAACAGCGGATTACCTATCCTATTGAAAATGCGATGGCAGGAATTCCAAATCTGGAAC
AAACTCGCTCCATTTCTCGTTATGGACTTTCCCAAGTCACCATCATTTTTAAAGATGGGACAGATATTTA
TTGGGCAAGACAACTGATTAACCAGCGTTTACAAGAAGCGGATGGCCAACTACCCGAATCAGTTGATCCT
ATCATGTCTCCTGTTTCAACAGGCTTGGGTGAAATTTACCAATGGGTGGTAAAAGCAAAATCAGGTGCTA
AGAAAGCAGATGGCACAGCCTATACAGCCATGGATTTACGTGAAATTCAGGACTGGATTGTACGTCCTCA
ATTGCAGCGTGTGAAAGGCGTGGCGGAAATCAACAGTATCGGTGGCTACAATAAAACCTATATTGTGTCA
CCTGATTTAAAACGTTTACAGCAGCTTCAGATTTCAATCAATGAATTTCAAACTGCTCTGCAAGAAAATA
ATGAAAATCGCGGTGCAGGTTTTATCGAAGAAAACGGAGAGCAACTCACCGTTCGTGTGCCGGGCATGTT
AAGTAGTGTTGAAGATATTCAAAACATTACTGTAAGTACCAAAAATGGTTTGCCGATCCGCGTGGCTGAT
GTTGCGAATGTCTCTATTGGGCATGATTTAAGAACAGGGGCTGCAACTTACAACGGTGAGGAAACGGTTC
TTGGCATTGCCATGATGATGATGGGAGAAAACAGCCGTACAGTTGCTCAGGCGGTTGATACCAAAATTAA
GGAAATACAATCTACCTTACCTAAAGGGGTGGAAATCGAGACGGTTTATGACCGGACGAGCTTAGTAAAC
AAAGCAATTGCGACGGTGCAGAAGAACCTTGTAGAAGGCGCGATTCTGGTTATTGTGATTTTGTTTATTT

TTCTAGGAAATTTTCGAGCTGCTTTAATCACAGCCTGCGTTATTCCACTTTCGATGTTATTTACTCTGAC
AGGTATGGCTGAACAAAACATTAGTGCCAACCTCATGAGCTTAGGAGCGCTCGATTTCGGGATTATTGTG
GATGGCGCGGTAGTTATTGTAGAGAACTGTATTCGACGTCTGGCAGAAGCACAGCATGCTCTACATCGGC
CTCTTACACGGTCTGAACGATTCAAAGAAGTTTTCCTTGCAGCAAAACAGGCCCGTCGCCCACTTATTTT
TGGGCAAATGATTATTTTGGTGGTCTATTTACCTATTTTTGCCCTATCTGGTGTTGAAGCCAAAATGTTC
CATCCAATGGCGATGACAGTGGTGATGGCATTATTGGGTGCCATGATTCTTTCTGTGACATTTGTACCTG
CGGCAGTTGCTCTTTTTGTTACGGGTGAAGTGAAAGAAAAAGAAACACGTTGGATGCAGCTTTTAAAGCA
GAAATATCAAAATATTCTCGATCAGGCTTATCAACTTAAAATTGTGGTCGTTAGTTTTGCATTAAGTATT
TTAGTTCTCACAGGTGTATTAGCTACCCAAATGGGTAGTGAATTTGCTCCGCAGCTCAGTGAAGGTGACT
TTGCCCTGCAGCAAATGCGTTCACCAAGTACTGGTCTCGAGCAATCACTGCGAATGCAGGAAAATACTGA
AAAGCTCATTTTGAAAAACTTTCCAGAAGTGAAAGCTGTTTTTGCTCGAACAGGGACAGCTGAGGTTGCA
ACTGATGTGATGCCGCCGAATATCTCTGATGCGGTCATTTTGCTTAAACCGCATGATCAATGGCCAAACC
CGAAAGAGACGCTAAGTGAGCTACGTTCACGTATGGAAGCTTTCTTGGCAACCTTACCGGGTAATAACAG
TGAGTTCTCTCAACCGATTGAGCTACGCTTTAATGAGCTAATTTCAGGGATTCGTAGTGATATTGGCGTC
AAGATTTTTGGCGACGATATGCAAGTACTCAATGAGCAGGCACAGGCACTCGCTCAGAAAGTACAAAAAA
TTTCAGGTGCTACCGCGGTTAAGGTAGAACAAACGAGCGGTTTACCGGTGCTGAGTGTTGAAATTAACCG
ACCTCTGGCTGCGCAATACGGATTATCTGCAAAAGCTATTCAAGATATTGTGGCGGCAAGTGTTGGTGGA
CAAAATGTTGGGCAGATTTTACAAGGCGATAGACGATTTGATTTTGAAATTCGCCTAGAAGACCAGCAGC
GTACCATCCAAAACTTAGCTCAACTTCCGATTCAATTACCAAATGGTGGACTCATCCAACTTCAAGATGT
GGCTAAAGTTGAACGTACCTCGGGCCTAAATCAGGTGGACGTGAAAATGGTAAGCGCCGTGTCATTATT
ACTGCAAACGTAGAAGGCCGTGACTTAGGATCATTTGTTCAAGAGTTAAGAGCAACACTCGCAAAAGAAC
AACTGCCAGCGGGCTATTGGTTGGAATATGGCGGTCAGTTTGAAAATCTCGCTTCAGCTGCTGCGCGAAT
GAAAATTGTTGTTCCATTGGCGCTTGCTATGATTTTTATTCTGCTCATGGCTGTATTCCATAATGTTAAA
GAAAGTCTGTTGGTCTTTAGCGGCGTGCCGTTTGCTTTGTCAGGTGGTCTGATTGCGCTTTGGCTAAGAG
ATATTCCACTGTCCATGTCGGCTGGCGTTGGGTTTATTGCATTATCTGGTGTCGCTGTTTTGAATGGTTT
GGTGATGCTGAGCTTTATTAAAGAGCTTAGAGAAAAATTTGATATTCAAACAGCCACGTGGAATGGGGCG
ATCTTACGTTTAAGACCGGTACTCATGACGGCTTGTGTTGCTTCACTTGGTTTTATTCCGATGGCTTTGG
CTACGGGAACTGGTGCAGAAGTTCAGCGGCCTTTGGCAACAGTGGTTATTGGTGGCATTATTTCATCTAC
GATATTAACTTTGGTTTTATTACCGGTCATTTATCGATGGATGAATGAAGACAAGACGAAAAGTGTTGAG
CATTCATAA
>AciRND~~~czcB_baumannii~~~CU459141.1~~~CzcB periplasmic adaptor protein of CzcABCD efflux pump in Acinetobacter baumannii
TTGGCAACAAGCTCAAACTTTTCCAACTCAGGCAGGAGAATAATAATGTCGGCAAATTTAAAGAAAAATT
CTCAATGGCTTTGGGTTGGAGTAATTGCTGCCATTACTGCAATATTAATTGGTTTACTCGTTTTAAATTC
AAAAAATAAATCTAATTCTTCGGAACCATCGGAAGGCCACGGGCATGCTGAAGAAGAGGGTGAAGAACAC
CATGATGAGGGAGAAAAACCACTACTACTCACTGCTCAGCAAATGCAGGAACAAAATTTAAAAATTGAAC
AAGCTGAACTAGGTGAAGTTCCTCAACTTCAGACTTATCCAGCCAAACTAGTAGTTAATACTGACCGCCA
AGCCCATGTTTCGCCAAGTTTTAGTGGTCGTGTCGAAGCGGTATATGTTGAACTAGGACAACAAGTTAAA
AAAGGCCAGGCACTTGCAAGCTTATTAGTGCCAGATTTAGTCGATCAGCAAGCCAATTTGCAAATAGCCC
AGTCTAATCTTGAGTTAGCACGTCAGGACTATGAGCGTGAACGTAGCTTATGGTCTCAAGGAATTTCCGC
GAAACAAGATTATCAACGAGCTTATAACGCTTATCAGCAAGCACAAATTCAGGTTAAAGCATCTCGCTCG
CGCTTAAGTGCATTCGGGGCAGGTTCGGGTTCAGCAGGGCGTTATACATTAACAGCGCCGATTGCGGGTA
TCGTGAGTAATAAAGATATTGTAGTGGGTGAAAACGTACAGTTAGCAGATCAGCTTTTTATTATTAATCA
GCTTGATCAGTTATGGCTGGAATTCATTTTACCAAGCAATGCAAATATCAATGTACAGCCAAATCAACAG
ATTGAATTTAAATCTTTACAAACTGGGAATACATTTTCTGCTCAGGTTCAAAGTTTAATAACAGAGGCAG
ATGCTCAGACTGGACGCTTACAAGTGCGTGCCAAGGTTTTGGCAAATAGCAGTGAGCTGCGTCCAAACTT
GATGGTTAACGTTGAGCTAAATGCAGGATCAACACAAAAAGTGACTCGTGTAAAAGCGCAAGCAGTTCAA
CAAGTTGAAGGTAAAGACGTTATCTTTACACCAAAAATAGTCAAAACAGGTTTTGAGTTTGAACCTGTGA
CTGTACAGCTAGGCCAACGTTCTAAAGATGGTCAATGGGTTGAAGTTGTAAAAGGAATTAATCCAAGCCA
ACGCTATATTGCAGAAGGCAGCTTCTTACTGAAGTCCGAATTGGAAAAAGGAGAGGCTGAGCATGGACAT
TAA
>AciRND~~~czcC_baumannii~~~CU459141.1~~~CzcC associating protein of CzcABCD efflux pump in Acinetobacter baumannii
ATGTCTTCTATTTTTTTAAATCGGGTGAGCCATCCCGACTTATTGAGCTGTAACGTTAAGATAACGTTGA
AAAAGGTTGTAACTATCTCAAGCCTGTCTATTGTCATGTTATTGGCGGGACAAATGGCAAATGCTGCGAC
TGATTTTCAACAGGGCTCTTATACCCAAAAAGCAGCTTTTAGTTTTGAACAGGCATTAGCGCGAGCACAA
AGCTATCAAATTCAACAAGGTGTTTGGCAGGCACAGCAGCAAATGGCTGAGGCGCAATTAAAACAAAGTC
GCTTATGGGCAAATCCAAGTCTTTCTATTGAGCAAACAGGTTTGCAGAGTGACCAAGAAAAAGAACTCGC
GATTGGCATCTCTCAACCTCTAGATATATTTTTGGGCAGCGTAAGGCCGCACAGCATTTAGCTAAAGTAGAA
ATGTCAAAGGTTGACTTGGCCGAACAACGTTATAAGGCTGAGCTTGAGTTAATAGTTAAATATTTCTGGT
CACAAGTGGCCTTGCTTGAACTTGAAAAGTCTCTCATTGGAGAACAGTTAGCAGTTAGCCAAGAGAACTT
GTCTGCATCGGAAAAGCGTTATCAGGCAGGGAGTATTGCTCAGGTTGATGTAGATCGAGTACGTATGTCT
CACTTGGAAAACCAGCGTTTATATCAGCAGGTCGATTTGAAACTACAAGTTGCTAAACAACAACTGGCCA
ACTTATGGGGCGGTGATTCAAATCAGTTCCAACTATCTCAAAGCTCTAATCAGCTGTGGGTATTGGCAGC
GGATGTAGAGTCTGGTCAAGATCGGCAAAACAATTTGCTAGAGCGCTCTTTTCAACTAGATGCTCTTGCG
CAGCAAGCCACTATTCAACAGCTTAAGGCAAAAGCAAGGCCACAACCTACGGTAACTTTGGGGGTGAATA
ACACTCGTTCTCCAGAGCAGCGTACGGAAAATCAAATCCGTTTAGGGGTGGAAATTCCTTTAAATCTTTT
CAATCGTCAACAATACGGAATCAAGATTGCTCAGGCCAAGCAAGAGCTGTCTCAACGCCAACAGAGCTTT
TATCGACAGCAGAATCAAGCGGATATTGAAACTCTGATGTCAGAGTTAAAAGGCTTACATATTCAGTTCA
AGCAATTAAATGATCAGCAAGTTCCTCTTGCCGTTCAGGTTCAACAAAAAACATTGCAAGGTTTCCGTTT
GGGTAAGTTTGCCGTTACCGATGTGCAGCAAGCCACAATGCAGTTGCAAGATGTGCGCTTACGTAAAGTC
GAGCTATTAAAACAGGCTTGGCAAAATTCAATTGAAATTCAAAGTTTACGTCTAGGGCTAGAGCCAGAAC
AGATCATGGCGAAAGATGCCTTAATGCAACTTAATCAACGTGCTTGGCAACAAGCTCAAACTTTTCCAAC
TCAGGCAGGAGAATAA

>AciRND~~~czcD_baumannii~~~CU459141.1~~~CzcD outer membrane channel protein of CzcABCD efflux pump in Acinetobacter baumannii
ATGGGTGGACATCATGGTCATGATCATAGTCATGCTGTAGTGACTGAGGGTAATGCTAAAAAATTAACGA
TTGCCTTGGCGCTTACCACGACATTCTTAATTGTTGAGGTGATTGCAGGTTTAATCACACAAAGTTTGGC
ATTGCTCTCTGACGCTGCACATATGTTTACAGATGCAGCTGCTTTAGCAATTGCTTTGGTTGCCATACAG
ATTTCTAAACGTCCTGCCGATAATAAACGTACTTTCGGTTATCAGCGCTTTGAAATTCTGGCCGCTTTAT
TTAATGCACTTATGCTTTTTGTGGTGGCAATTTATATTTTATATGAAGCCTATATCCGCTTTTCGCAGCC
ACCTGAAATTCAAAGTGTAGGTATGCTCATTGTGGCGACCATTGGTTTGGTAATAAACCTCATCTCAATG
AAAATTCTTATGTCGAGTGCTAATAACAGCTTAAATGTGAAAGGTGCTTATCTAGAAGTATTGAGTGATG
CACTAGGCTCAGTTGGCGTTATTATTGGTGCAATTATTATTTACTTCACTAATTGGTATTGGATTGACAC
GCTTATTGCGGTACTGATTGGATTTTGGGTATTGCCAAGAACATGGGTTTTACTTAAACAAAGTATTAAT
ATTTTGCTCGAAGGTGTACCCGAAGAAGTCGATATTGAAAAGCTACGTGCAGATTTACTTTCATTAAATG
GTGTGGAGAGTATTCACCAACTCAAAGTATGGGCAATTACCTCTAAAAATATCCATTTAACTGTACACTT
ATTTGCGCCTGAAGCTGACCGTAACAAGCTCTATCAAGATGCAGTTGAAATGCTTTCTCATGAGCATGGT
ATTGGTGAAGTGACATTGCAAATTGAAGATGATGCTGAGATTAACTGTCAGCATATTGCTCAACATGCTT
CACACGAGCATAACGATAATGACAAGACGCATTCACATCAGCATTAA
>AciRND~~~abeD_baumannii~~~CU459141.1~~~AbeD RND pump protein characterised in Acinetobacter baumannii
ATGCTATCTAAATTTTTTATTCAACGCCCCATTTTTGCCAATGTATTGGCGATCATTGTTATGGCTTTCG
GTATTTTTTCGGTTATGAATTTGCCTGTAGAACGGTATCCAGACATTGCTCCACCTAAAATTACTGTGTC
AGCCAACTATAGTGGTGCAGATGCACAAACGGTTGAGCAAAGTGTTACTCAAATTTTAGAACAGCAAATA
CAAGGGATTGATCACTTACTCTATTTTAGTTCATCGAGTGACTCATCTGGACGTAGCCGAATTACTATAA
GTTTTGATAACGGAACAAATCCGGATACTGCTCAGGTCCAAGTACAAAATAGTATTAGTGGTGTCATACG
TCGCTTACCTGATGAAGTTCAACGCCAAGGTGTTACGGTAAGTAAGTCACTGGGTGACACTTTTATGGTA
ATTGGCTTATATGACTCGACTGGTAAAACAGGAAACATTGAGCTATCGGACTATTTAACTACGCATGTGG
TAGATAACCTGAACCGTATTGAAGGGGTGGGTGAAACTGATATATTTGGTTCACAATATGCCATGCGTAT
CTGGTTAAATCCCGATAAATTAAAACAATATAATTTAATGCCAAGTGATGTAGCGAATGCAATCACCGCA
CAAAATACTCAGGTCGCCGCAGGGGCAATTGGTGACTTACCCGTAATTGACGGTCAATATTTAAATACAA
AAGTCACAGCAGGTTCTCGCTTAAAAACAGTTGAGGATTTTAAAAATATTGTCGTGAAGTCGAATAAAAC
AGCGAGTTATGTGTATTTAAAAGATATTGCCAGAGTTGAGCTAGGTGCAGAAAACTATCAGTCTTTTAAC
ACTATTAATGGCTATCCTGCCGCAGGTTTGGGTATTTCTTTATCTTCGGGTGCAAATGCAATTCAGACCT
CTAAGCTCATCCACCAAACTCTAGATCAGCTTACAACGAAACTACCAGCGGGTTATAAAATCGTTTATCC
ACGAGATAATACGCCCTTTGTTCAAGAATCAATTAAGGAAGTAGTAAAGACTCTGGTAGAGGCGATCATT
CTGGTTATTTTGGTCATGTTCCTGTTCTTACAAAGCTGGCGTGCTACGCTCATTCCGAGTATTACCGTTC
CAGTTGTGATTTTAGGAACTTTCGCTGTCTTATATGTGCTTGGTTTTAGTATTAACACCTTAACGTTATT
TGCGCTGGTACTCGCGATTGGTTTGCTGGTCGATGACGCGATTGTGGTCGTAGAAAACGTTGAGCGGCTC
ATGCATGAACAGCACTTATCTCCTAAAGAAGCTGCTATTGAGTCGATGGGGGAAATTAGTGGTGCCTTAG
TCGGGATTACGTTGGTTTTAACTGCTGTTTTTATTCCAATGTCCTTTTTAGGCGGTTCAATTGGGGTGAT
TTACCGTCAGTTTTCTATTACTTTAGTTGCCGCTATGGCGTTGTCGCTTATTGTTGCGCTCATTTTAACA
CCGGCTTTATGTGCATTAATTTTAAAACCAAATCCTCAACCTCAGCGTTGGGCAGTATGGTTTAACCAAA
AGATTGAGCAACTTAAAAATCAATATATCAAGCTTGTTCAGACGAGTATTCATTACAGTAAATCAGTTAT
TGTGATTTTTGTGGCTTTAATTGCCGTTTTTACGCTGTTCTATAACGGTTTAAAAAGCGGTTTTATTCCT
AAAGAAGACCAAGGAATTTTAAGTGTTCAAATTAAGCTCGTAGACAGCGCACCAATTTCTCAAAGCCAGA
AAATTGGTGAGCAAGTCCGCCAATATTTCTTAACTCAAGAAGATAAAAATGTAGATTTGGTTTTAATCCG
CTATGGACGAAATTATTCGGGCACAGGACAAAACTTGGCACAAGGGTTTATTGCTCTAAAACCGTGGGAT
GTCCGAACAGGAAAAGAAAACTCGGCTGAGGCTATACAAAAGCGTGCCATGAAATACTTTAGTCATTTTA
ATAATGCACAGATTAATGTGACTTTACCTGCCTCAGTTAATGGCTTAGGTCAAACAGATGGTCTGGATTT
ATGGATTCAGGATTTGAATGGGCAAGGGCAAGATTTTCTAGATAGTGCCTTCCGCCAATTGCAGGCTCAA
AGTAAAAATTATTCAACTTTCGAAAACTTTGATAAGCAGTCAACCAATAGCAAGGCAAATCTTAATATTA
AGATTGACCAGAAACAGGCACTAGCAAATGGATTACAGCTATCGGCAATTAATAACACTTTGTCGAGCGC
ATGGGGCGGAACTTATGTAAATGACTTTATTGATCGGGGCCGTATTAAACGTGTCATGATTCAAGGTGAT
GCCGAGTTTAGATCTAAACCGGAAGATTTATATAACTGGTCTGTACGTAATGACCAAAATGAAATGGTTC
CCTTTAGTTCATTTGCCAACTTTAGCTGGGGCGGGGCGCCAGAAATTGTAAAACGCTATATGGGATATAG
TGCTTTACAACTACAAGCAGATGTCGCGAGTGGCAGCAGTTCTGGCCAAGCGATGAAAGATGTAGAACAA
CTTGTTAACCAACAAAAAGATATTGGTTTAGCGTGGACAGGTTTATCTTTTGAAGAACAGAAGTCGACTA
ATCAGGCAGTGTGGTTATATTTAATTTCGGCTGGATTTATTTTCTTATGTTTGGCTGCTTTATATGAAAG
CTTAAGTATTCCCGCCGCCGTAATGACATCTATTCCGCTTGGTGTAGGAGGAAGTGTGATTTTCTCTTAT
ATTTTTGGCTTGCCAAATGACGTGTATTTCCAAATTGCGCTATTAACGACTATTGGTTTGTCATGTAAAA
ATGCCATTTTAATTGTTGAGTTCGCGGCCTTAGCGCAAGAAAAAGGTAAGAATGCCATTCAGGCAGCCTT
AGAAGGTGCGAGCTTACGATTAAGACCGATTCTAATGACCTCTTTAGCCTTTGGGGCAGGCGTAATTCCG
CTTGTGTTTGCTCAAGGCGCTGGTGCGGTTAGTCGTCAAGAGATTGGTATTAGTATTTTAGGTGGCGTGA
TGTTTGGTACTGTGCTGGTTCTGTTTTTTATTCCGGTCATGTACGTGTTATTACGTTCACTGTTTAAATC
GAAAGCTTCAACCTAA
>AciRND~~~arpA_baumannii~~~CU459141.1~~~ArpA periplasmic adaptor protein of ArpAB efflux pump in Acinetobacter baumannii
ATGAATCTGTTAAAACCTCTCATCATGAGTATGATGATTGTTTGCATGGTCACATTAATGGGGTGTAGTA
AAGAGGCTCCCAAAACAGAAGAAATACCCTATGTGATGGTGACCCAGCCTTCAACCACACTTCACGAACA
AAAAAGCTATGCTGGAGATGTACAGGCTCGACAACAAACTGCCTTGGCATTTCGGGTGGGTGGACAAGTT
ACGGCTCGCTATGTGGATGTAGGTGACCGGGTTAGAGTTGGGCAAGTATTAGCAAAACTCGATGTGGCGG
ATGCACAGCTACAATTAAATGCTGCCAAAGCTCAATTAGAAAATGCACAGGCAGCAGCAAAAACAGCCTC
AGATGAGCTTAAGCGGTTTCAACAATTATTACCCATAAATGCCGTGAGCCGTTCGCAATACGATACGGTA
AAAAATCAATATGATGCGGCCCAGGCAGCATTACAACAAGCTCGTTCTAATTATGAAGTTTCTGCCAACC
AGACTGGTTATAACCAACTTGTTTCTAATAAAAACGGGGTGATTACAGCGCGTAATATTGAAATTGGACA
GGTGGTTGCAGCAGGGCAAGCGGCTTATCAACTGGCAATTGATGGTGAACGTGAAGTGGTCATCGGCGTA

CCGGAACAAGCGGTTAGCGAGATTAAAGTTGGCCAAGCGGCATGGATAACTTTGTGGTCTAAACCGAACG
AACGATTTGCCGGATATGTACGCGAAGTTTCTCCAGCTGCTGACCAGTCCCGTACATTCACAGTTAAGGT
GGCACTCAAAGAAGGTCAGTCTGCTATTCAGCTGGGACAAAGTGCACGCGTATTTTTTAGTTCGACTCAA
ACTAATGTCATGAGTGTGCCACTTTCGAGTGTATCTGCAACAGATAACCAACCTTATGTATGGGTGGTGA
ATGCGAATCAGACCTTACGTAAAGTGCCTGTAACGATTGGTGCTTATGCCAGAGATAGTGTGCCGGTATT
ATCGGGTTTAACACCAAATGATTGGGTAGTGATTGGTGGTGTGCACTTGCTGCGCGATAAACAGAAGATT
CACCCGATTGATCGTGAAAATCGTGCAGTGAAAATTCAGGGAGCCAAATAA
>AciRND~~~arpB_baumannii~~~CU459141.1~~~ArpB RND pump protein of ArpAB efflux pump in Acinetobacter baumannii
ATGAAATTTAATCTCTCTGAATGGGCACTGAACAATAAGGGTATTGTCCTTTATTTCATGCTCTTGCTCG
GCATTATTGGTGCAATTTCTTATTCAAAACTCTCACAAAGTGAAGATCCGCCATTTACCTTTAAAGTCAT
GGTCGTACAAACCTACTGGCCAGGCGCGACAGCCAAAGAAGTTTCTACTTTAGTTACGGACCGTATCGAA
AAGGAACTGATGACCACAGGTCAGTATGACAAGATTATGGCGTATTCCCGTCCAGGCGAGTCGATGGTGA
CTTTTGTTGCTAAAGATTCTCTCACTTCTGCGCAAATTCCTGATGTTTGGTACAACGTTCGGAAGAAGGT
CAATGACATTCGCCATGAACTCCCAAGTGGTGTGCAAGGTCCATTTTTTAATGATGAATTTGGCGATACT
TTCGGTAATATTTATGTACTGACAGGCAAAGACTTTGACTACGCGCTTTTGAAAGAATATGCCGATCGTT
TGCAATTACAACTACAGAGAGTCAAAGATGTAGGCAAAGTTGAGCTGATCGGGCTACAAGATCAGAAAAT
CTGGATTGAAATTTCAAACACTAAAGCGGTTCAGCTCGGTATTCCTGTTTCTGCCATACAAGAAGCCCTG
CAAAAGCAAAATAGCATGGCAAGTGCAGGCTTTTTTGAAACTGGAACGGATCGTATTCAAATTCGAGTAA
GTGGCCAATTACAAAGCGTAGAGGACATTAAAAAAATGCCTTTACTGGTAGGCGATAAAACCATTCAGCT
TGGTGACGTTGCTGATGTCTACCGTGGTTTTAGCCAACCTGCTCAGCCGCGTATGCGTTTTATGGGTGAC
AATGGTATTGGTATTGCCGTATCTATGCGTAAAGGCGGCGATATTATTGCCTTAGGTAAAAATCTGGAAA
CTGAATTTGCCCAACTGCAAAAAACATTACCTCTAGGTATGAAACTACAAAAAGTATCTGACCAACCGGT
AGCGGTACAACGTAGTATCCATGAGTTTGTCAAAGTACTTGCTGAAGCGGTCATTATTGTCTTGTTAGTG
AGCTTTTTCTCATTAGGTTTCCGGACGGGTTTAGTTGTCGCTTTTTCCATTCCTTTGGTTTTAGCAATGA
CTTTTGCTGGCATGAATTTATTTGATGTCGGGCTGCACAAGATCTCGCTTGGTGCCCTAATTCTAGCTTT
GGGTTTGCTTGTAGATGATGCCATTATTGCTGTCGAGATGATGGCCATTAAGATGGAGCAGGGGTATAGC
CGAATTAAGGCCGCCGGATTTGCATGGAAAACAACAGCATTTCCGATGTTGACGGGGACATTAATTACCG
CCGCAGGCTTTTTACCTATTGCTACGGCTCAGTCCAGTACAGGTGAATACACACGCTCTATCTTTCAGGT
CGTGACGATTGCTTTATTGGTGTCGTGGGTTGCCGCAGTTTTATTTGTACCTTATTTGGGTGAAAAACTA
CTACCTGATTTTACCAAGACCGGTCATCAAGCACCTTGGTATGTCCGTTTATGGGCAAGAGTAACTAAAA
AACCGCAACCACAAACTGTGGCCATTTCACAGGACCACCATTACGATCCTTATCAATCTAGTTTCTATTT
ACGTTTTAGGAAAATGGTCGAGTTTTGTGTGACCTACCGTAAAACCGTGATTGCAACAACGGTGGGGATT
TTTGTGCTGTCTGTACTCATGTTTAAGATGGTGCCACAGCAGTTTTTCCCGCCTTCAAACCGCGCTGAAA
TTTTAGTCGATTTAAAACTCGAAGAAGGCGCATCGTTAACCGCTACAGAGCAAGCGGTGAAAAAGGTTGA
ACAATTCCTGTCTAAACAAAAAGGCATTGATAATTATGTAGCCTATGTCGGTACAGGTTCACCACGTTTT
TATTTACCTCTAGACCAGCAATTACCGCAAGCCAGCTTTGCGCAGTTTGTTGTTTTGGCATCATCGCTTG
ATGATCGTGATGAAATTCGCCGTTCTTTAGAAACCCAAATTAAGCAGTTGCTCCCACAAGTCCGTACTCG
CGTGTCATTACTCGAAAATGGCCCACCTGTTGGTTATCCATTGCAGTATCGTGTGTCAGGTGAAGATTTA
AATCTGGTACGTAAAGAAGCACAGCAGGTTGCTAGGGTAATTAGTGAAAACCCGAATACCACCAATGTGC
ATTTGGATTGGGGTGAGCCAAGCAAGATTATTTCAATTCAAATTGATCAAGACCGTGCTCGACAAATGGG
TGTGTCCAGCCTCGATTTAGCCAACTTCTTAAACGCCTCAATTACAGGTAGTGCGATTGAGCAATACCGT
GAAAAGCGTGAGCTGATTGAAATCCGATTACGTGGTGATAAAGCTGAGCGTGTTGAAGTAGCTTCACTGG
CGAGCCTTGCAGTACCAACTGCGAATGGAACAACTGTACCTTTAGCTCAAATTGCGAAGATTGAGTATAA
GTTTGAAGATGGTCTGATTTGGCACCGTAATCGGTTACCGACAATTACTGTTCGTGCAGATATTCGTACC
AATTTACAGCCAGCTACCGTTGTTGGTGAGTTAGCTGAATCAATGGACAAGTTACGCGCTGAGCTGCCAA
GTGGCTACCTCATTGAAGTGGGGGGAACAGTGGAAGAGTCGGCACGCGGACAAAGTTCGGTCAATGCCGG
TATGCCACTCTTTTTGGCAGTGGTCATGACATTACTCATGATTCAGCTGAAGAGTTTATCTCGGGCAACA
ATTGTATTTTTGACTGCACCATTAGGCTTAATTGGCGTTGTTTTATTCTTACTTTTGTTTAATAAACCAT
TTGGTTTTGTTGCCATGCTAGGAACCATTGCCTTATCCGGCATGATTATGCGTAACTCACTCATTCTGAT
TGATCAGATTGAACAAGACAGACAGGCAGGGCATCCAACGTGGGAAGCAATTATTGATGCAACAGTACGC
CGTTTCCGTCCGATCATTCTCACGGCATTGGCAGCAGTACTCGCCATGATCCCTCTTTCGCGGAGTATTT
TCTTCGGTCCAATGGCTGTTGCGATTATGGGCGGACTCATTGTTGCTACCTTGCTGACATTATTTTTCTT
ACCTGCATTGTATGCAGCGTGGTTTAAGGTGAAAAAAACAGCATAA
>AciRND~~~acrA_nosocomialis~~~CP029351.1~~~AcrA periplasmic adaptor protein of AcrAB efflux pump in Acinetobacter nosocomialis
ATGAATCTGTTAAAACCTCTCATCATGAGTATGGTGATTGTTTGCACGGTCACATTAATGGGGTGTAGTA
AAGAGGCACCCAAAACAGAAGAAATACCCTACGTGATGGTAACGCAGCCTTCGACCACACTAAACGAATT
AAAAAGCTATGCAGGAGATGTACAGGCTCGACAACAAACTGCCTTGGCATTTCGAGTAGGTGGACAAGTT
ACGGCTCGCTATGTCGATGTAGGTGACCGAGTTAAAGTTGGACAAGTATTAGCGAAACTCGATGTAGCAG
ATGCACAGTTACAATTAAATGCGGCCAAAGCTCAATTACAAAATGCACAGGCAGCAGCGAAAACAGCATC
AGATGAACTTAAACGTTTCCAACAATTATTACCTGTAAATGCTGTTAGCCGTTCACAATTTGATACGGTA
AAAAATCAATACGACGCAGCGCAAGCAGCGTTACAACAAGCCCGCTCTAACTATGAAGTTTCTGCAAACC
AAACTGGTTATAACCAACTCATTTCTAATAAAAACGGGGTGATTACCGCGCGTAATATTGAAATTGGACA
AGTCGTTTCAGCAGGACAAGCGGCTTATCAATTGGCGATTGATGGTGAACGTGAAGTAGTTATAGGCGTA
CCAGAACAAGCAGTTACCGAGATCAAGGTTGGTCAAGCAGCTTGGATTACCTTGTGGTCTAAACCGAATG
AAAAATTTGCTGGATATGTACGTGAAGTTTCTCCAGCGGCTGACCAGTCTCGTACTTTTACAGTCAAAGT
AGCACTCAAAGAAGGCCAATCTGCTATTCAGCTTGGACAAAGTGCACGGGTATTTTTTAGCTCGACTCAA
ACCAATGTCATGAGTGTGCCACTTTCGAGTGTTTCTGCCACAGATAATCAGCCCTATGTATGGGTCGTAA
ATGCAAATCAAACCTTACGCAAAGTACCTGTAACCGTGGGAGCTTATGGCCGAGACAGTGTGCCTGTATT
AACGGGGTTGACACCAAATGATTGGGTTGTGATTGGTGGTGTCCATTTACTACGTGATAAGCAGAAGATC
CATCCGATTGACCGTGAAAACCGCGCGGTGAAAATTCAGGGGGCAACAAAGCCATGA
>AciRND~~~acrB_nosocomialis~~~CP029351.1~~~AcrB RND pump protein of AcrAB efflux pump in Acinetobacter nosocomialis

ATGAAATTCAATCTCTCGGAATGGGCACTGAACAATAAAGGTATCGTCCTTTATTTTATGTTGTTGCTTG
CCATTATCGGTGCAATTTCCTATTCCAAATTATCTCAAAGTGAAGATCCGCCATTTACCTTTAAAGTGAT
GGTTGTCAAAACATTCTGGCCTGGAGCGACTGCCAAAGAAGTTTCAACGTTAGTGACTGATCGTATTGAA
AAGGAACTGATGACAACTGGGCAGTATGAAAAGATTATGGCCTACTCCCGTCCAGGTGAGTCGATGGTTA
CATTTGTTGCTAAGGACTCGCTCACTTCTGCACAAATTCCTGATGTTTGGTACAACGTTCGTAAAAAGGT
TAATGACATTCGTACTGAACTGCCAAGTGGTGTACAAGGTCCTTTTTTCAATGATGAGTTCGGGGATACT
TTCGGTAATATTTATGTACTGACAGGCAAAGACTTTGACTATGCACTTCTCAAAGAATATGCAGATCGTT
TGCAATTACAACTACAAAGAGTCAAGGATGTTAGTAAAGTTGAATTGATTGGCTTACAAGATCAAAAAAT
CTGGATTGAGATTTCAAATACAAAAGCAGTCCAACTCGGTATTCCTCTTTCTGCAATTCAAGAAGCCTTA
CAAAAGCAAAATAGTATGGCAAGCGCAGGTTTCTTTGAAACCGGTACAGACCGTATTCAAATACGTGTAA
GTGGGCATTTAAACAGCGTTGATGAAATTAAAAAAATTCCTTTATTAGTCGGCAATAAAACCATTCAGCT
TGGTGATGTTGCTGATGTCTACCGTGGTTTTAGTCAACCGGCTCAGCCACGTATGCGTTTTATGGGGGAA
AATGGTATTGGTCTCGCCGTGTCTATGCGTAAAGGCGGGGATATTATTGCCTTGGGCAAAAATCTAGAGT
CCGAATTTGCCCAGCTCCAAAAAACCTTACCTTTAGGCATGAAACTACAGAAGGTATCTGATCAACCTGT
AGCAGTACAACGTAGTATTCATGAATTTGTAAAAGTACTCGCTGAAGCAATCATTATTGTCTTGTTAGTA
AGCTTTTTCTCTTTAGGTTTCCGTACCGGATTGGTAGTCGCTTTTTCCATTCCTTTAGTTTTAGCAATGA
CTTTTGCCGGCATGAATGTATTCGATGTCGGGTTACATAAAATTTCTCTCGGTGCTCTGATTTTAGCTTT
AGGTCTACTGGTTGATGACGCCATTATTGCAGTCGAGATGATGGCAATTAAGATGGAACAGGGCTATAGC
CGAATTAAGGCTGCTGGATTTGCATGGAAAACAACGGCCTTTCCAATGCTGACAGGAACATTGATTACCG
CGGCAGGTTTTTTACCTATCGCTACGGCACAGTCCAGTACGGGTGAATATACACGTTCAATCTTTCAGGT
GGTGACGATTGCCTTATTGGTGTCTTGGGTTGCCGCAGTTTTATTTGTACCTTATTTGGGTGAAAAACTA
CTGCCTGATTTTACCAAGACTGGGCATCAAGCACCTTGGTATGTTCGTTTATGGGCAAGATTAACGAAAA
AACCACAACCACAAACGATCGAGATTTCACAGGACCATCATTACGATCCTTATCAATCTAGTTTCTATTT
ACGTTTTAGGAAAATGGTCGAGTTTTGTGTGACCTATCGTAAGACTGTAATTGCGACAACGGTTGGGATA
TTCGTGCTCTCTGTACTGATGTTTAAAATGGTTCCTCAGCAGTTCTTCCCACCATCAAACCGTACCGAGA
TTTTAGTCGATCTAAAACTTGAAGAAGGCGCTTCTTTAACTGCTACAGAACAAGCGGTGAAAAAGGTTGA
ACAGTTCCTGTCTAAACAAAAAGGTATTGATAACTATGTGGCCTATGTGGGTACAGGCTCACCACGTTTT
TATTTACCGTTAGACCAGCAATTACCACAAGCCAGCTTTGCACAGTTTGTTGTTTTAGCGACCTCACTTG
ATGACCGTGATGAAATTCGTCGCTCGCTAGAAACTCAAATTAAACAGTTGCTCCCACAAGTTCGTACCCG
TGTGTCATTACTGGAAAATGGTCCGCCTGTAGGTTATCCATTGCAGTATCGTGTGTCAGGTGAAGATCTC
AATCTGGTACGTAAAGAAGCACAGCAGGTTGCGAAGGTGATTGGTGAAAACCCAAATACTACCAATGTAC
ATTTGGACTGGGGTGAGCCGAGCAAAATCATTGCGATTCAGATTGATCAAGATCGTGCTCGACAAATGGG
TGTCTCAAGTGTGGATTTGGCAAACTTCCTGAATGCTTCAATTACAGGTAGTGCGATTGAACAGTATCGT
GAAAAGCGTGAACTCATTGAAATCCGTTTACGTGGAGATCAAGCTGAGCGTGTTGAAGTCGCTTCACTGG
CAAGTTTAGCGGTCCCAACAGCCAATGGAACGACAGTACCGTTAGCCCAGATTGCGAAAATTGAATATAA
GTTTGAGGATGGCCTGATCTGGCATCGTAACCGCTTACCAACAATTACCGTACGTGCAGATATTCGTACC
AAATTGCAGCCAGCGACTGTTGTTGGCGAGTTAGCCGAATCAATGGATAAGTTACGTGCTGAGTTACCAA
GTGGTTACCTCATTGAAGTGGGGGGAACTGTCGAAGAGTCAGCACGAGGACAAAGTTCGGTCAATGCTGG
TATGCCATTATTTTTGGCAGTGGTCATGACATTACTCATGATTCAGCTTAAGAGTCTATCGAGAGCAACT
ATTGTATTTTTGACTGCACCATTAGGCTTAATAGGAGTAGTTCTGTTCTTACTTTTATTTAATAAGCCAT
TTGGTTTTGTGGCTATGCTGGGAACCATTGCTTTATCTGGAATGATTATGCGTAACTCGCTCATTTTGAT
TGATCAGATTGAACAAGACCGGCAAGCAGGGCATCCAACATGGGAAGCGATTATTGACGCAACCGTACGC
CGTTTCCGTCCAATTATTCTGACAGCCTTGGCAGCGGTATTAGCAATGATCCCACTTTCACGAAGTATTT
TCTTCGGTCCAATGGCCGTTGCAATTATGGGTGGACTGATTGTAGCCACCTTGCTGACATTATTTTTCCT
GCCTGCATTGTATGCTGCGTGGTTTAAGGTGAAAAAAACAGCATAA
