## Supplemental 4 - structure for "RND pumps across the *Acinetobacter* genus; AdeIJK is the ancestral efflux system"

**Supplementary S4:**

**Supplementary S4 text 4:** Detailed text regarding the structure of the binding pockets of AdeJ/X

The principal ligand-binding pockets of the RND pumps are the PBP and the DBP, and the structural analysis of their geometry, combined with positional conservation analysis of the residues lining these pockets can be used to inform predictions on the drug-binding specificities of the respective pumps.

As mentioned in the main text of the manuscript, the AdeJ and AdeY reveal very close overall homology (with identity approaching 80%) and positional conservation, allowing to isolate them as a distinct subgroup of RND pumps. On the other hand, the sequence alignment of *Ab*AdeJ with the closest paralogue *Ab*AdeB (figure 6), reveals several deletions in the latter (corresponding to *Ab*AdeJ residue numbers 509-513; 594-597; 854-857; 923-929 and 1048-1053 inclusive), as well as insertions (602; 607-608, 1043-1046). Despite that, *A. baumannii* AdeJ and AdeB share close structural organisation and reflecting that, a structural alignment of *Ab*AdeJ ((7M4P:C) (1)), with *Ab*AdeB EtBr-bound form AdeB-ET-III, (7KGI: C (2)) resulted in an R.M.S.D. of just 0.998 Å over the *Ab*AdeJ C-alpha backbone. This is marginally better than the corresponding AdeJ vs AcrB superposition of 1.17Å, although this figure is conformation dependent (supplementary S4, figure 5).

At least 21 residues form the PBP within *E. coli* AcrB (3–5), and there is a high conservation of the pocket between the AdeJ/Y and its RND orthologues with the notable exception in the region covered by residue range 660-688 (*Ab*AdeJ numbering), which forms the bottom section of the PBP and includes the so-called F-loop (668-682) (figure 6; supplementary S4, figure 6, B). Strikingly, while this region (660-688) is nearly fully conserved within the AdeJ/Ys, it shows a major deviation *from both* the *Ab*AdeB (1,2,6) and the orthologous AcrB transporters from *E. coli* and *Salmonella* (7,8), *Neisseria* MtrD and the *Pseudomonas* MexB (9,10).

The F-loop itself shows full positional equivalency between the AdeJ/Y and AcrB. However, in comparison with the F-loop of AcrB, where this loop features the prominent residue F666, preceded by an A665, both of which form part of the hydrophobic binding site at the lower entrance of PBP, in *Ab*AdeJ (and similar to *Ab*AdeB), this pair is replaced by a Leucine (L666) and a Proline (P665) respectively, giving a markedly different character to the loop and its dynamics. At the same time, the AdeJ/Y F-loop is also very different from the one seen in AdeB, where it features a 3-residue deletion immediately after L666, thus clearly differentiating the AdeJ/Y-type transporters from the AdeB-type. This feature suggests a common, but distinct mode of pump action between the AdeJ/Y and the rest of the RND family, once again highlighting their close relationship and uniqueness.

The uniqueness of the PBP of AdeJ/Y is further highlighted by a distinctive V573, conserved across the AdeJ/Y, but which in AcrB/MtrD is M573, is substituted by a Valine within the entire AdeJ/Y clade (as well as in MexB), and by a Trp in AdeB. Once again, these differences might be responsible for the differential efflux profiles of AdeJ/Y and AdeB. Intriguingly, the neighbouring AcrB M575, which also plays an important role in coordination macrolides and rifamycins (and which is also conserved in both MtrD and AdeB), shows limited variation within the AdeJ/Y clade, and, in AdeY, is substituted by a Leucine.

Beyond these regions however, the PBP is surprisingly conserved, with S79, Q579, F618, E675, L676, G681, and G721, conserved not only among the AdeJ/Y-clade members, but also in AdeB, AcrB and even *Neisseria gonorrhoeae's* MtrD (9,11). Another prominently conserved residue within the AdeJ/Y-subfamily is R718 (*Ab*AdeJ numbering), found in the front of the PBP (supplementary S4, figure 6, panel B). It corresponds to R717 of AcrB and R714 of MtrD, where it has been shown to be important for coordination of macrolides and rifamycins (3,4,12,13), contributing to substrate specificity (14–16). Indeed, the described conservation of the PBP appears to be reflected in the rifampicin MICs seen in Table 2. It is therefore rather intriguing that that residue is not conserved in AdeB, where it is substituted by Tryptophane, suggesting a significantly different coordination of potential ligands it the PBP between AdeJ/Y and the former.

Taken together, the PBP of the AdeJ/Y-transporters shows close conservation to the overall more-distantly related AcrB/MtrD/MexB pumps, but a clear departure from the AdeB clade, and it is tempting to speculate that these discrepancies are responsible for the differences in the efflux profile between AdeJ/Y and AdeB (17,18). Indeed, earlier work on *Ab*AdeIJK and *Ab*AdeABC reconstituted in *E. coli* (17), showed AdeIJK being significantly more effective in the efflux of β-lactams, novobiocin, and ethidium bromide than the referent AcrAB-TolC pump, but being notably less effective in erythromycin efflux. At the same time AdeABC was more effective in tetracycline efflux but had lower efficiency of extrusion of lipophilic β-lactams, novobiocin, and ethidium bromide. Recent analysis of the substrate specificity of the latter in *E. coli* (13), suggested that it provides a better protection towards polyaromatic compounds and lower resistance towards antibiotic compounds compared to AcrAB-TolC, and while reproducing of MIC profiles using *Acinetobacter* strains is more challenging, our data, in particular on tetracycline and rifampicin seems to cross-corroborate these earlier reports, which can also be linked to the expectations based on the architecture of the AdeJ/Y-binding pockets. This is particularly pronounced in the description of the PBP, which is considered to be responsible for the recognition of the higher molecular weight drugs, including ansamycins and macrolides, but also may contribute to processing of tetracyclines (19,20).

As mentioned above, the PBP and DBP are separated by the flexible G-loop (covering residue range 613-624 in *Ab*AdeJ), containing a conserved phenylalanine (F618), revealed to be involved in drug-binding in the DBP of both the newly determined *Ab*AdeJ fluorocycline-bound structures (1,21) and in the AdeB (2). Within the AdeJ/Y the whole of the G-loop is strictly conserved, which suggest similar binding properties in the upper part of the PBP and the front part of the DBP. The only exception could be considered the flanking residue S614, which is divergent between the AdeJ/Y, and also AdeB and AcrB, however it is outward facing and so unlikely to contribute directly to substrate specificity.

Of the 23 residues reported to provide the DBP drug-binding determinants in the *E. coli* AcrB (3,5), 6 are conserved among AdeJ/Y, AdeB, AcrB, and MtrD (1,2,6,9,13). Those correspond to *Ab*AdeJ residues F136, F178, Y327, V613, F618, and F629. Intriguingly, F611 (*Ab*AdeJ), the substitution of which in AcrB (F610A) has the been reported to have most significant impact on the substrate MICs (22), and which impacts the DBP dynamics (23), while preserved within the AdeJ/Y subfamily, as well as across AcrB/MtrD/MexB, is substituted by a Threonine in AdeB. While most of the rest of DBP substitutions are fairly conserved, a notable DBP residue that provides further differentiation between AdeJ/Y and AdeB is V573, which is substituted by a bulky aromatic W568 in the latter, and which, due to its unique position, also contributes to PBP. Once again, this pattern of conservation suggests that AdeJ/Y is functionally closer related to AcrB/MrtD than to AdeB.

However, it is the residues in the DPB which show discrepancy between the members of AdeJ/Y-clade that are perhaps of highest interest. Indeed, it is striking that while there are very few substitutions in the DBP, three out of four that are present (A46, Q91 and T128) are clustered together at the back of the pocket (supplementary S4, figure 6, panel C), forming a plausible interaction site, which is hinted by the covariation of the A46/T128 positions. which in the case of *A. lwoffi* and *A. baylyi* is instead represented by a S46 in combination with either R128 or K128 respectively. The last DBP residue to show variation within the AdeJ/Y clade is Y327 (F in *A. lwofii*) and is found at the bottom of the pocket, opposite side across from the A46/T128 pair (Supplementary figure 6, panel C). This is the only significant departure within the AdeJ/Y-clade in general and warrants further future investigation, as it may point to subtle differentiation between the pump substrates. Our analysis shows that the substitution of the small beta-branch hydroxylated residue with a long positively charged side-chain not only introduces a big steric barrier at the back-side of the DBP, but also leads to a significant change of the electrostatics and hydration pattern in the pocket, which is expected to impact some ligands in this region. Indeed, the presence of the R128/K128 side chain in either *Al*AdeJ or *Ab*AdeJ, would provide steric clashes with the crystallised binding modes of doxorubicin, as seen in PDB ID 2DR6 and PDB ID 4DX5 (4,24), where all 3 residues are involved in doxorubicin coordination, and of the pyridopyrimidine derivative EPI D13-9001 (as seen in PDB ID 3W9H-3W9J; (25)). Molecular dynamics (23) have also suggested that these residues are involved in both doxorubicin, D13-9001 and phenylalanine-arginine β-naphthylamide (*PAβN*).

The last DBP residue to show variation within the AdeJ/Y clade is Y327 (F327 in *A. lwoffii*) and is found at the bottom of the pocket, opposite side across from the A46/T128 pair discussed above. The residue occupies a critical position in the pocket and has been implicated in both eravacycline and the TP-6076 fluorocycline binding as evidenced by the recent *Ab*AdeJ structures (PDB IDs 7M4P and 7RY3 respectively (1,21). In departure from the previous pattern, it is conserved between AdeJ/Y and AdeB. Residues in equivalent position to Y327 have been reported to be involved in the binding of pretty much all small planar compounds and inhibitors described above in other related pumps, including AcrB (Y327), MtrD (Y325) and MexB (Y325), but due to the conservative nature of the substitution, we assess that is unlikely to cause a major phenotypic effect. However, it is worth noting, that the variable residues from the PBP (namely M575 and T679), do not just contact each other, but also form an arc with the variable Y327 in DBP, which may suggest some joint selective pressure between these substitutions (supplementary S4, figure 5C).

Taken together, while AdeJ/Y type transporters display close similarity of overall structure and potential drug-recognition determinants, which are partly shared with AcrB/MtrD/MexB type pumps, while showing clear separation from the AdeB. However, they possess a number of structural peculiarities, that land support to differentiating AdeJ/Y into a separate structural subfamily of RND-transporters and hint towards a distinct functional role for them, which is yet-to-be determined.


1. Zhang Z, Morgan CE, Bonomo RA, Yu EW. Cryo-em determination of eravacycline-bound structures of the ribosome and the multidrug efflux pump adej of acinetobacter baumannii. MBio [Internet]. 2021;12(3). Available from: https://journals.asm.org/doi/abs/10.1128/mBio.01031-21
2. Morgan CE, Glaza P, Leus I V., Trinh A, Su CC, Cui M, et al. Cryoelectron microscopy structures of adeb illuminate mechanisms of simultaneous binding and exporting of substrates. MBio [Internet]. 2021;12(1):1–15. Available from: https://journals.asm.org/doi/abs/10.1128/mBio.03690-20
3. Nakashima R, Sakurai K, Yamasaki S, Nishino K, Yamaguchi A. Structures of the multidrug exporter AcrB reveal a proximal multisite drug-binding pocket. Nat 2011 4807378 [Internet]. 2011 Nov 27;480(7378):565–9. Available from: https://www.nature.com/articles/nature10641
4. Eicher T, Cha HJ, Seeger MA, Brandstätter L, El-Delik J, Bohnert JA, et al. Transport of drugs by the multidrug transporter AcrB involves an access and a deep binding pocket that are separated by a switch-loop. Proc Natl Acad Sci U S A [Internet]. 2012 Apr 10;109(15):5687–92. Available from: https://pubmed.ncbi.nlm.nih.gov/22451937/
5. Vargiu A V., Nikaido H. Multidrug binding properties of the AcrB efflux pump characterized by molecular dynamics simulations. Proc Natl Acad Sci U S A [Internet]. 2012 Dec 11;109(50):20637–42. Available from: https://pubmed.ncbi.nlm.nih.gov/23175790/
6. Su CC, Morgan CE, Kambakam S, Rajavel M, Scott H, Huang W, et al. Cryo-Electron Microscopy Structure of an Acinetobacter baumannii Multidrug Efflux Pump. MBio [Internet]. 2019 Aug 27;10(4). Available from: https://mbio.asm.org/content/10/4/e01295-19
7. Murakami S, Nakashima R, Yamashita E, Yamaguchi A. Crystal structure of bacterial multidrug efflux transporter AcrB. Nature [Internet]. 2002;419:587–93. Available from: www.nature.com/nature
8. Johnson RM, Fais C, Parmar M, Cheruvara H, Marshall RL, Hesketh SJ, et al. Cryo-EM Structure and Molecular Dynamics Analysis of the Fluoroquinolone Resistant Mutant of the AcrB Transporter from Salmonella. Microorganisms [Internet]. 2020 Jun 1;8(6):1–21. Available from: https://pubmed.ncbi.nlm.nih.gov/32585951/
9. Lyu M, Moseng MA, Reimche JL, Holley CL, Dhulipala V, Su CC, et al. Cryo-EM structures of a gonococcal multidrug efflux pump illuminate a mechanism of drug recognition and resistance. MBio [Internet]. 2020 May 1;11(3). Available from: https://journals.asm.org/doi/abs/10.1128/mBio.00996-20
10. Sennhauser G, Bukowska MA, Briand C, Grütter MG. Crystal structure of the multidrug exporter MexB from Pseudomonas aeruginosa. J Mol Biol [Internet]. 2009 May 29;389(1):134–45. Available from: https://pubmed.ncbi.nlm.nih.gov/19361527/
11. Chitsaz M, Booth L, Blyth MT, O'mara ML, Brown MH. Multidrug Resistance in Neisseria gonorrhoeae: Identification of Functionally Important Residues in the MtrD Efflux Protein. MBio [Internet]. 2019 Nov 1;10(6). Available from: https://pubmed.ncbi.nlm.nih.gov/31744915/
12. Tam HK, Foong WE, Oswald C, Herrmann A, Zeng H, Pos KM. Allosteric drug transport mechanism of multidrug transporter AcrB. Nat Commun 2021 121 [Internet]. 2021

Jun 29;12(1):1–10. Available from: https://www.nature.com/articles/s41467-021-24151-3
13. Ornik-Cha A, Wilhelm J, Kobylka J, Sjuts H, Vargiu A V., Malloci G, et al. Structural and functional analysis of the promiscuous AcrB and AdeB efflux pumps suggests different drug binding mechanisms. Nat Commun [Internet]. 2021 Dec 1;12(1). Available from: https://pubmed.ncbi.nlm.nih.gov/34824229/
14. Middlemiss JK, Poole K. Differential impact of MexB mutations on substrate selectivity of the MexAB-OprM multidrug efflux pump of Pseudomonas aeruginosa. J Bacteriol [Internet]. 2004 Mar;186(5):1258–69. Available from: https://pubmed.ncbi.nlm.nih.gov/14973037/
15. Yu EW, Aires JR, McDermott G, Nikaido H. A periplasmic drug-binding site of the AcrB multidrug efflux pump: a crystallographic and site-directed mutagenesis study. J Bacteriol [Internet]. 2005 Oct;187(19):6804–15. Available from: https://pubmed.ncbi.nlm.nih.gov/16166543/
16. Trampari E, Holden ER, Wickham GJ, Ravi A, Prischi F, de Oliveira Martins L, et al. Antibiotics select for novel pathways of resistance in biofilms. bioRxiv [Internet]. 2019 Apr 10;605212. Available from: https://www.biorxiv.org/content/10.1101/605212v1
17. Sugawara E, Nikaido H. Properties of AdeABC and AdeIJK efflux systems of Acinetobacter baumannii compared with those of the AcrAB-TolC system of Escherichia coli. Antimicrob Agents Chemother [Internet]. 2014 Dec;58(12):7250–7. Available from: http://www.ncbi.nlm.nih.gov/pubmed/25246403
18. Migliaccio A, Esposito EP, Bagattini M, Berisio R, Triassi M, De Gregorio E, et al. Inhibition of AdeB, AceI, and AmvA Efflux Pumps Restores Chlorhexidine and Benzalkonium Susceptibility in Acinetobacter baumannii ATCC 19606. Front Microbiol [Internet]. 2022 Feb 7;12. Available from: https://pubmed.ncbi.nlm.nih.gov/35197939/
19. Nakashima R, Sakurai K, Yamasaki S, Nishino K, Yamaguchi A. Structures of the multidrug exporter AcrB reveal a proximal multisite drug-binding pocket. Nature [Internet]. 2011 Dec 22;480(7378):565–9. Available from: https://pubmed.ncbi.nlm.nih.gov/22121023/
20. Tam HK, Foong WE, Oswald C, Herrmann A, Zeng H, Pos KM. Allosteric drug transport mechanism of multidrug transporter AcrB. Nat Commun 2021 121 [Internet]. 2021 Jun 29;12(1):1–10. Available from: https://www.nature.com/articles/s41467-021-24151-3
21. Morgan CE, Zhang Z, Bonomo RA, Yu EW. An Analysis of the Novel Fluorocycline TP-6076 Bound to Both the Ribosome and Multidrug Efflux Pump AdeJ from Acinetobacter baumannii. MBio [Internet]. 2022 Feb 1;13(1). Available from: https://journals.asm.org/doi/10.1128/mbio.03732-21
22. Bohnert JA, Schuster S, Seeger MA, Fähnrich E, Pos KM, Kern W V. Site-directed mutagenesis reveals putative substrate binding residues in the Escherichia coli RND efflux pump AcrB. J Bacteriol [Internet]. 2008 Dec;190(24):8225–9. Available from: https://pubmed.ncbi.nlm.nih.gov/18849422/
23. Vargiu A V., Ruggerone P, Opperman TJ, Nguyen ST, Nikaido H. Molecular mechanism of MBX2319 inhibition of Escherichia coli AcrB multidrug efflux pump and comparison with other inhibitors. Antimicrob Agents Chemother [Internet]. 2014 Oct 1;58(10):6224–34. Available from: https://pubmed.ncbi.nlm.nih.gov/25114133/
24. Murakami S, Nakashima R, Yamashita E, Matsumoto T, Yamaguchi A. Crystal

structures of a multidrug transporter reveal a functionally rotating mechanism. Nature [Internet]. 2006 Sep 16;443(7108):173–9. Available from: http://www.ncbi.nlm.nih.gov/pubmed/16915237

25. Nakashima R, Sakurai K, Yamasaki S, Hayashi K, Nagata C, Hoshino K, et al. Structural basis for the inhibition of bacterial multidrug exporters. Nature [Internet]. 2013;500(7460):102–6. Available from: https://pubmed.ncbi.nlm.nih.gov/23812586/

**Supplementary S4, figure 5:** Structural similarity of AdeJ to other RND pumps

**A.**

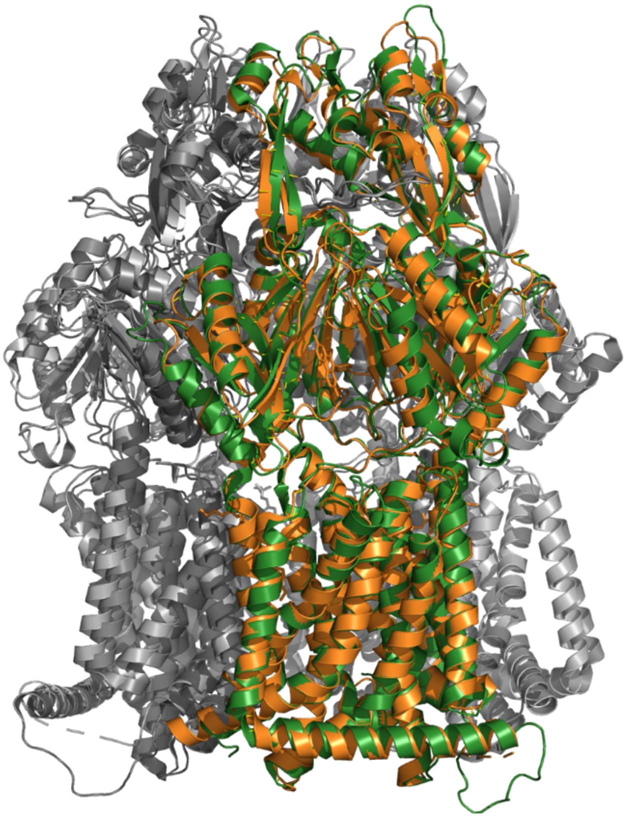

**B.**

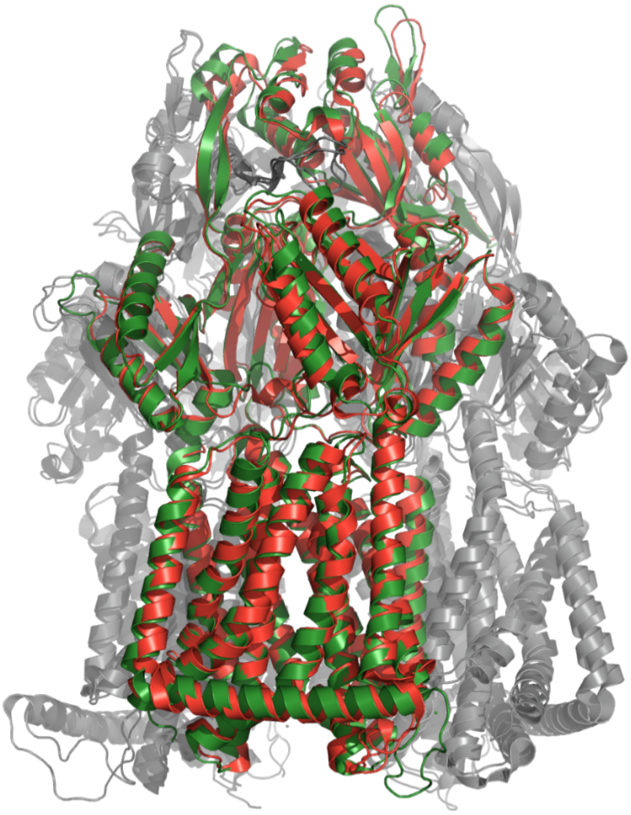

A - *Ab*AdeJ (green) superposed over *Ab*AdeB (orange) showing the close relation between the two pumps. AdeB is based on the experimental structure PDB ID 7KGI resulting in an RMSD of under 1

Angstrom (0.998Å) over the C-alphas. B - *Ab*AdeJ (green) superposed over *E. coli* AcrB (red). The RMSD RMSD of 1.17Å. Based on the experimental structure PDB ID 4DX5.

**Supplementary S4, figure 6:** *Ab*AdeJ trimer model (A), PBP features (B) and DBP features (C)

**A.**

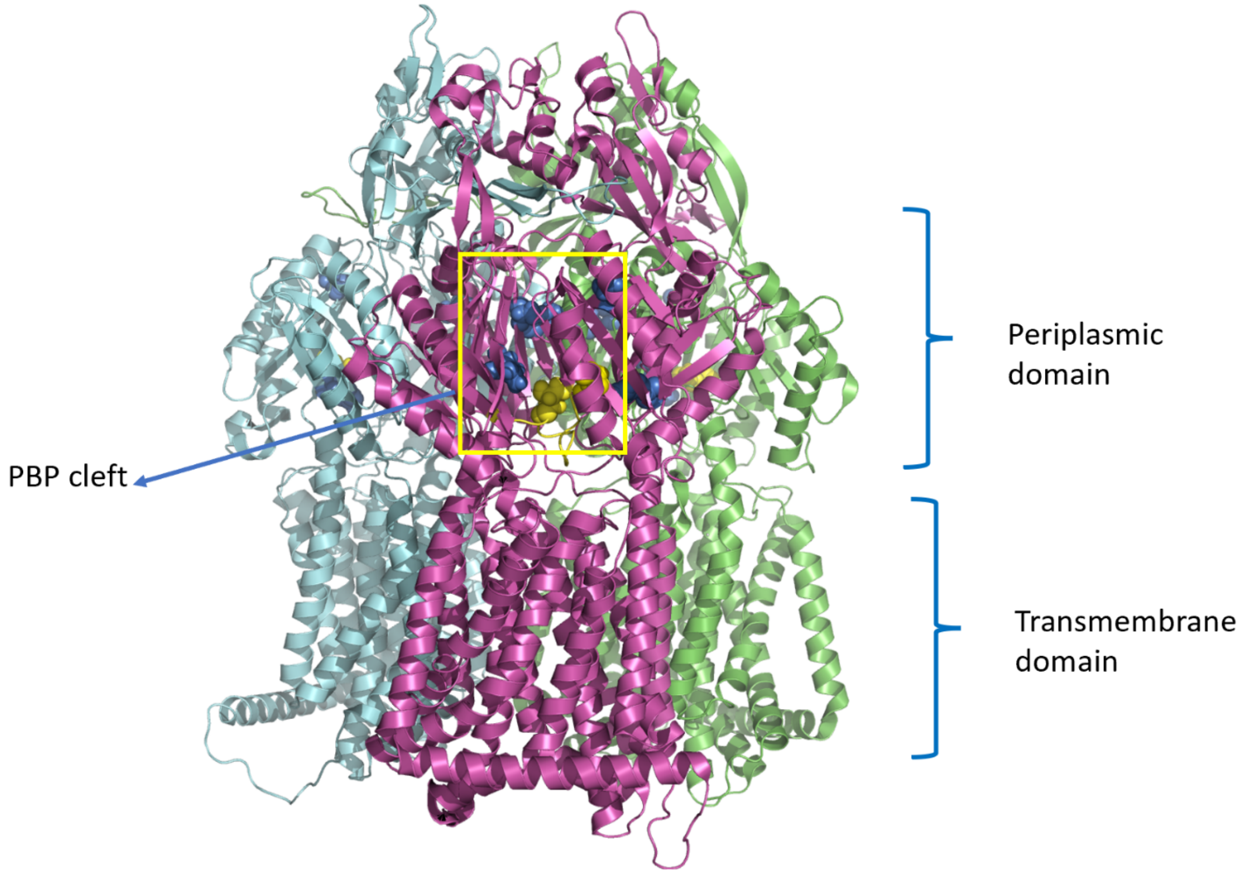


**B.**

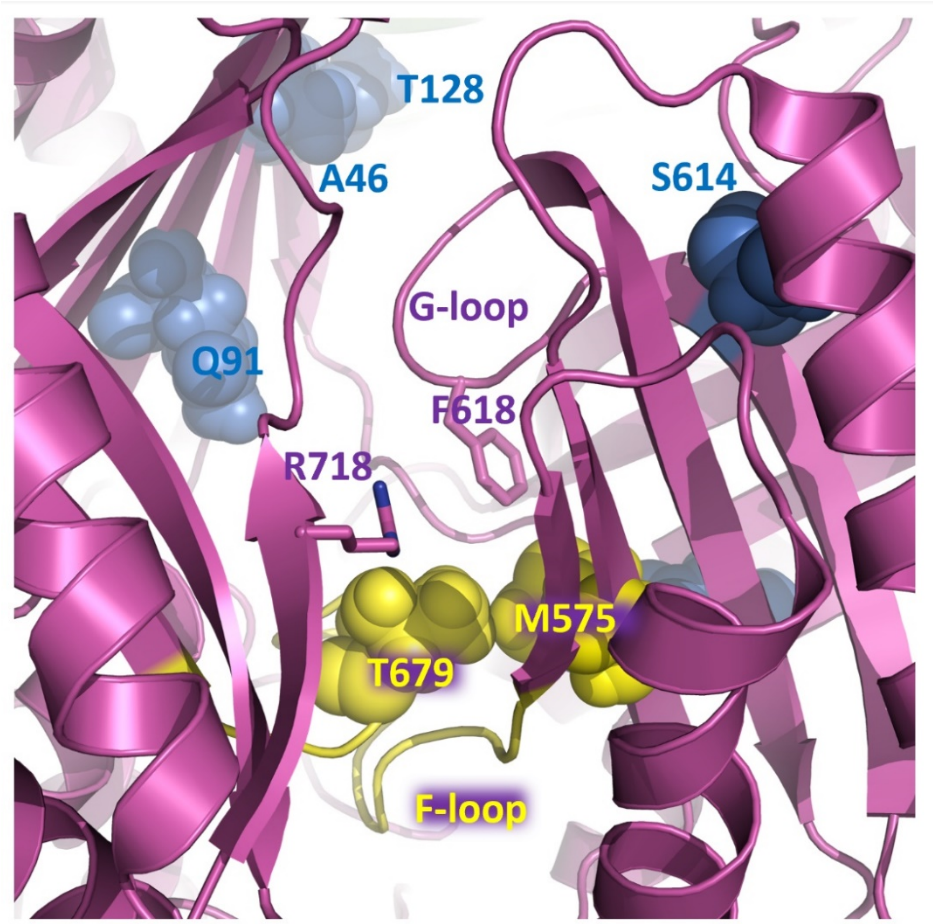

**C.**

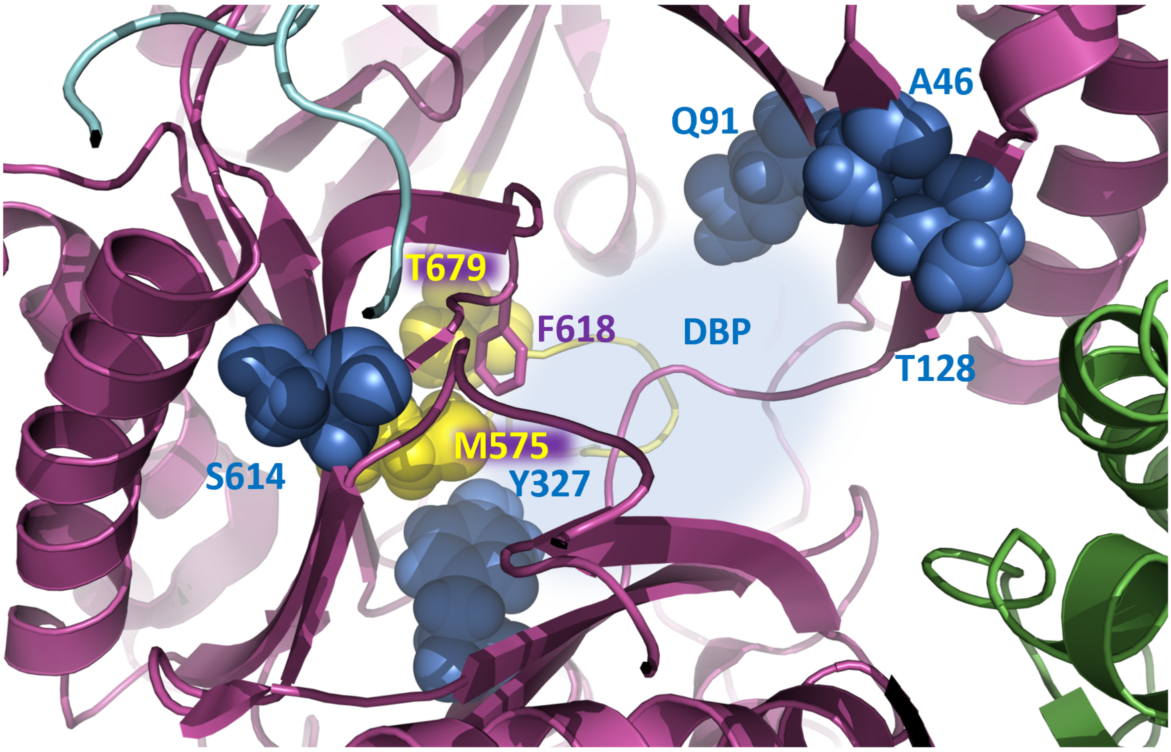


A - General view of the AdeJ trimer (based on the experimental structure of AbAdeJ PDB ID 7M4P), highlighting the PBP (yellow) and DPB (blue) residue positions that show discrepancy between the members of the AdeJ/Y.

B - Main features of PBP, and the prominent/divergent residue positions discussed in the text are shown based on the experimental structure of AbAdeJ (7M4P.pdb; chain C). Divergent residues are shown in space-fill. Those belonging to the PBP are coloured yellow; the ones belonging to the DBP are coloured blue.

C - Main features of the DBP, seen from the top of the protomer. Some elements of the top of the pocket and the funnel domain have been removed.

**Supplementary S4, table 6:** Residue conservation in the principle drug binding pockets across AdeJ/Y and comparison with residues within AcrB.
Conservation is colour-coded as follows. Complete conservation across AdeJ/Y/AcrB – green; Divergence between the AdeJ/Y – purple; Non-conservative substitutions divergent from AcrB – in red. Where additional data or residue context is available it is commented upon. Residue numbers only shown in AdeY and *Al*AdeJ where the number is different from the *Ab*AdeJ.

| AdeJ *A. baumanii* | AdeY *A. baylyi* | AdeJ *A. lwoffii* | AcrB *E. coli* | *Comments* |
|---|---|---|---|---|
| | | *PBP* | | |
| *S79* | *S* | *S* | *S79* | *Strictly conserved* |
| *Q91* | *E* | *S* | *T91* | *Divergent within AdeJ/Y. In AdeB it is also a T.* |
| *A134* | *A* | *A* | *S134* | *Conserved; Semi-conservative substitution relative to AcrB; also an S in AdeB* |
| *S135* | *S* | *S* | *S135* | *Conserved* |
| *K292* | *K* | *K* | *K292* | *Conserved* |
| *V573* | *V* | *V* | *M573* | *Conserved across AdeJ/Y; Significant substitution relative to AcrB, diagnostic for the AdeJ/Y subfamily* |
| *M575* | *L* | *M* | *M575* | *Divergent within the AdeJ/Y, however conserved relative AcrB* |
| *Q579* | *Q* | *Q* | *Q577* | *Conserved* |
| *F618* | *F619* | *F619* | *F617* | *PBP/DBP – G-loop* |
| *G622* | *G623* | *G623* | *G621* | *Conserved* |
| *A625* | *A626* | *A626* | *T624* | *Conserved in AdeJ/Y. A semi-conservative substitution relative to AcrB; and a conservative substitution relative to AdeB (V)* |
| *F629* | *F630* | *F630* | *F626* | *Conserved* |
| *S663* | *S664* | *S664* | *M662* | *Conserved across AdeJ/Y; A non-conservative substitution relative to AcrB and AdeB (M in AdeB). The region shows breakdown of alignment.* |
| *M666* | *M667* | *M667* | *F664* | *Conserved across AdeJ/Y; A non-conservative substitution relative to AcrB. Breakdown of alignment* |
| *L668* | *L669* | *L669* | *F666* | *Conserved across AdeJ/Y; A non-conservative substitution relative to Acr. Breakdown of alignment.* |
| *Q669* | *Q670* | *Q670* | *N667* | *Conserved across AdeJ/Y; Semi conservative substitution* |
| *L670* | *L671* | *L671* | *L668* | *Conserved* |
| *E675* | *E676* | *E676* | *E673* | *Conserved* |
| *L676* | *L677* | *L677* | *L674* | *Conserved* |
| *V678* | *V679* | *V679* | *T676* | *Conserved across AdeJ/Y; Semi-conservative Substitution relative to AcrB and AdeB (T in AdeB)* |
| *G681* | *G682* | *G682* | *G679* | *Conserved* |
| *F682* | *F683* | *Y683* | *F680* | *Conserved* |
| *N683* | *N684* | *N684* | *D681* | *Conserved across AdeJ/Y; Semi-conservative Substitution relative to AcrB* |
| *R718* | *R719* | *R719* | *R717* | *Conserved across AdeJ/Y and AcrB, as well as MtrD and MexB. Not conserved in AdeB (W in AdeB). Implicated in macrolide and rifampin binding* |
| *N720* | *N* | *N* | *N719* | *Conserved* |

| | | | | |
|---|---|---|---|---|
| G721 | G | G | G720 | Conserved |
| N827 | N | N | E826 | Conserved across AdeJ/Y. Non-conservative substitution in AcrB (S in AdeB!) |
| DBP | | | | |
| A46 | S46 | S46 | S46 | Divergent between AdeJ/AdeY, but semi conservative substitution relative to AcrB; Appears to co-vary with T128 |
| S89 | S | S | Q89 | Conserved across AdeJ/Y. Non-conservative substitution relative to AcrB and AdeB (E). |
| T128 | K128 | R128 | S128 | Divergent between AdeJ/Y<br>Appears to co-vary with A46. It is a Q in AdeB |
| T130 | T | T | E130 | Conserved in the AdeJ/Y, bit represents a non-conservative substitution relative to AcrB and AdeB (E130) |
| S132 | S | S | S132 | Conserved |
| A134 | A | A | S134 | Semi-conservative (S in AdeB) |
| F136 | F | F | F136 | Conserved |
| V139 | V | V | V139 | Conserved; Implicated in Eravacyclin binding as revealed by PDBID 7M4P (PMID: 34044590) |
| Q176 | Q | Q | Q176 | Conserved |
| V177 | V | V | L177 | Semi-conservative (S in AdeB) |
| F178 | F | F | F178 | Conserved (Eravacycline binding) |
| G179 | G | G | G179 | Conserved (Eravacycline binding) |
| G180 | G | G | S180 | Conserved within AdeJ/Y; Semi-conservative relative to AcrB (Eravacycline binding) |
| D273 | D | D | E273 | Conserved; Semi-conservative relative to AcrB (Q in AdeB) |
| N274 | N | N | N274 | Conserved |
| Y275 | Y | Y | Y275 | Conserved |
| F277 | F | F | I277 | Conserved across AdeJ/Y; Semi-conservative substitution in AcrB |
| Q276 | Q | Q | D276 | Conserved within the AdeJ/Y; Semi-conservative substitution relative to AcrB and AdeB(N in AdeB); Change of electrostatics of the pocket. |
| F277 | F | F | I277 | Substitution (Eravacycline binding) |
| A290 | A | A | G290 | Substitution |
| A326 | A | A | P326 new | Substitution (Eravacycline binding) |
| Y327 | Y | F | Y327 | Divergent between AdeJ/AdeY (semi-conservative substitution)<br>Implicated in Eravacycline binding |
| V573 | V | V | M573 | Conserved in the AdeJ/Y.<br>Major change of the character of the pocket relative to AcrB and AdeB (W) |
| F611 | F612 | F612 | F610 | Conserved<br>(T in AdeB); implicated in Eravacycline binding |
| V613 | V | V | V612 | Conserved<br>(I in AdeB); implicated in Eravacycline binding |
| S614 | A615 | A615 | N613 | Semi-conservative substitution within AdeJ/Y. Divergent from both AcrB (N) and AdeB(L). flanking the G-loop but outward facing, so unlikely to be impacting ligand binding directly. |

| | | | | |
|---|---|---|---|---|
| *F616* | *F617* | *F617* | *F615* | *DBP conserved (W in AdeB); Residue flanking the gating loop; Implicated in Eravacycine binding;* |
| *F618* | *F619* | *F619* | *F617* | *Conserved throughout RNDs (F612 in AdeB); gating loop residue* |
| *V621* | *V622* | *V622* | *R620* | *Non-conservative substitution, resulting in a major change in the DBP.* |
| *F629* | *F630* | *F630* | *F628* | *Conserved; implicated in Eravacycline binding* |
| | | *Additional features* | | |
| *R124* | *R* | *R* | *Q124* | *Conserved within the AdeJ/Y ; Non-conservative relative to AcrB (Q in AdeB) (exit gate as suggested PMID: 23175790)* |
| *Y759* | *Y* | *Y* | *Y758* | *Conserved (exit gate as suggested PMID: 23175790)* |
